# Mapping the Functional Landscape and Viral Diversity of 2A Peptides

**DOI:** 10.64898/2026.09.24.754024

**Authors:** Adesh Baral, Aditi Kothiala, Kevin X. Zhong, Janelle Cheung, Stephane Flibotte, Simcha Srebnik, Khanh Duc Dao, Curtis A. Suttle, Eric Jan

**Affiliations:** Department of Biochemistry and Molecular Biology, Vancouver, BC Canada; Life Sciences Institute, University of British Columbia, Vancouver, BC Canada; Department of Ocean, Earth and Atmospheric Sciences, University of British Columbia, Vancouver, BC Canada; Department of Chemical and Biological Engineering, University of British Columbia, Vancouver, BC Canada; Department of Mathematics, University of British Columbia, Vancouver, BC Canada

**Keywords:** ribosome, tRNA, translation, 2A peptide, peptidyl-tRNA hydrolysis

## Abstract

2A peptides are short (∼18–22 amino acid) sequences utilized by a subset of RNA viruses to process their polyproteins in an unusual co-translational event known as “StopGo” or “ribosome skipping”. This process involves ribosome pausing at a conserved C-terminal D(V/I)ExNPG↓P motif, followed by peptidyl-tRNA hydrolysis of the nascent peptide-tRNA^Gly^, and continued translation from the downstream proline codon. However, how nascent 2A peptide interacts with the ribosomal exit tunnel to promote these event(s) remains poorly understood and the molecular determinants governing 2A-mediated StopGo translation remain undefined. Here, we developed a mammalian fluorescence resonance energy transfer (FRET) based high-throughput reporter system to quantitatively measure 2A activity across large libraries of natural and engineered 2A peptides. Systematic mutational analysis of canonical viral 2A peptides from porcine tescovirus and foot-and-mouth disease virus (P2A, F2A) identified key residues required for activity and revealed critical contributions from previously uncharacterized N-terminal residues. Extending this approach to metagenomic viral datasets uncovered conserved sequences governing StopGo efficiency leading to functional ranking of diverse viral 2A peptides in mammalian cells, including identification of a novel subclass that supports high stop-go activity (>99%). Phylogenetic analysis of viral genomes revealed clustering of functional 2A peptides and specific motifs within specific viral lineages branches, suggesting an evolutionary trajectory underlying 2A diversification and optimization. These findings define the sequence and functional landscape of viral 2A peptides and provide mechanistic and evolutionary insights into how nascent peptide–ribosome exit tunnel interactions mediate StopGo translation.

## INTRODUCTION

Translational recoding allows ribosomes to reinterpret codons or the reading frame during protein synthesis, thereby enabling the production of multiple distinct polypeptides from a single mRNA without changes to its primary nucleotide sequence. This phenomenon has been documented across diverse viral and eukaryotic systems. Canonical examples include programmed ribosomal frameshifting, stop codon readthrough, and translational bypass, all of which modulate the ribosome during elongation or termination (Farabaugh, 1996; Gesteland and Atkins, 1996; Baggen *et al*., 2018). Because recoding outcomes depend on coordinated interactions between the mRNA, the nascent polypeptide, and the ribosome, these systems have served as models for dissecting how ribosomes integrate multiple molecular inputs to regulate translation.

One such recoding mechanism is 2A peptide StopGo recoding, in which translation of a single open reading frame yields two discrete protein products through site-specific release of the nascent polypeptide during elongation at a sense codon, followed by continued translation of the downstream sequence. The 2A peptide was first described at the foot-and-mouth disease virus (FMDV) 2A/2B junction, where viral polyproteins are separated without proteolytic cleavage (Ryan, King and Thomas, 1991). Subsequent biochemical and genetic analyses established that this separation reflects an alteration in ribosomal elongation rather than post-translational processing (Donnelly *et al*., 2001). 2A peptides are widely used in biomedical research and biotechnology to co-express multiple genes from a single construct, ranging from monoclonal antibody production to generation of pluripotent stem cells (Fang *et al*., 2005; Szymczak and Vignali, 2005; Luke and Ryan, 2024). However, despite their widespread use, relatively few studies have investigated the underlying mechanism of 2A-mediated recoding or systematically characterized 2A peptide diversity across viral and eukaryotic genomes.

2A-mediated recoding depends entirely on the eukaryotic translation machinery. Functional StopGo activity is observed in eukaryotic cell-free systems and in cells, but no 2A peptide tested to date has shown activity in prokaryotic translation systems (Donnelly *et al*., 2001; Rao *et al*., 2025). Studies in yeast suggest that eukaryotic release factors influence the efficiency and outcome of 2A-mediated separation, whereas reconstituted mammalian translation systems demonstrate that nascent-chain release can occur independently of eRF1 and eRF3 (Doronina *et al*., 2008; Sharma *et al*., 2012; Machida *et al*., 2014). A recent cryo-EM structure of the F2A-StopGo complex confirmed that eRF1-AAQ accommodated in the A-site does not contact the P-tRNA or nascent chain, indicating that the resulting complex reflects the pre-release state independent of eRF1 binding (Li *et al*., 2026). The molecular basis for the broader system-dependent differences in StopGo efficiency is not fully understood.

At the sequence level, functional 2A peptides are typically 18–22 amino acids in length and can be grouped into distinct classes. Class A 2A peptides, typified by picornaviral sequences such as those from FMDV (F2A), porcine teschovirus (P2A), equine rhinitis A virus (E2A), and Thosea asigna virus (T2A), share a conserved C-terminal region essential for activity centered on the motif DxExNPG | P (x denotes any residue), with nascent-chain separation occurring between the glycine and proline residues (Ryan and Drew, 1994; Donnelly *et al*., 2001). Mutational analyses demonstrate that the acidic residues, the invariant asparagine, and the NPGP junction are each critical for efficient recoding, with substitutions at these positions abolishing activity (Sharma *et al*., 2012). In addition, the N-terminal amino acids of 2A peptides (positions 1-12), though not essential, can promote StopGo reactions (Donnelly *et al*., 2001; Kjaer and Belsham, 2018; Rao *et al*., 2025). Class A 2A peptides contain an N-terminal central hydrophobic region enriched in leucine and other aliphatic residues (Rao *et al*., 2025). Although this N-terminal region lacks strict primary-sequence conservation, truncation and scanning-alanine experiments demonstrate that disruption of this hydrophobic segment reduces nascent-chain release (Sharma *et al*., 2012; Minskaia, Nicholson and Ryan, 2013). Conceptually, the Class A 2A peptide is divided into two elements: upstream amino acids that likely interact with the ribosomal exit tunnel to promote StopGO reactions and the essential PG|P motif where the peptide bond is skipped.

As a nascent translated polypeptide of approximately 30–40 amino acids are largely confined within the ribosomal exit tunnel, 2A peptide–ribosome interactions drive StopGo activity(Ryan *et al*., 1999; Voss *et al*., 2006; Doronina *et al*., 2008; Li *et al*., 2026). In a recent report, a cryo-EM structure of a mammalian ribosome stalled at the FMDV 2A site revealed that the F2A nascent chain is positioned deep within the exit tunnel, where the conserved hydrophobic segment (FDLLKL) forms a hydrophobic cluster that packs against tunnel-lining elements formed by 28S rRNA and ribosomal protein uL22 (Li *et al*., 2026). The conserved polar residues D12, E14, and N16 form hydrogen bonds and electrostatic interactions with 28S rRNA nucleotides near the peptidyl transferase center. These contacts likely induce conformational changes in the PTC that shift the P-tRNA CCA end away from its canonical position, inhibiting peptide bond formation at the glycine–proline junction and pre-exposing the peptidyl-tRNA ester bond for hydrolysis (Li *et al*., 2026). Single alanine substitutions at hydrophobic positions (F4A, L6A, L7A, K8A, L9A) reduced activity to varying degrees, while D12A and N16A abolished activity entirely (Li *et al*., 2026). Toeprinting, primer extension, and ribosome profiling assays confirm that ribosomes pause at the StopGo site, with the G18 codon in the P-site and the P19 codon in the A-site (Atkins *et al*., 2007; Doronina *et al*., 2008). Li et al. (2026) further showed that while proline at position 19 is optimal, substitutions with Asp, Gln, or Trp retained over 70% activity in vitro, indicating that the identity of the incoming tRNA aminoacyl moiety is not strictly deterministic for StopGo. Beyond productive StopGo, ribosome drop-off and reduced downstream translation have been observed at some 2A sites, with efficiencies that vary across sequences and experimental systems (Donnelly *et al*., 2001; Doronina *et al*., 2008).

Large-scale bioinformatic analyses have recently revealed that 2A/StopGo peptide sequences are far more widespread than previously appreciated. Rao et al. (2025) used an iterative HMMER-based approach to identify over 2,200 potential unique 2A peptide sequences across viral ORFs in RefSeq, UniProt, and the MGnify metagenomic database, expanding the known repertoire by nearly an order of magnitude. In this study, they identified Class B 2A peptides, which are generally longer and more sequence-divergent than Class A 2A peptides. Class B 2A peptides contain a distinct motif containing an asparagine/histidine immediately upstream of the terminal PGP and a characteristic invariant N-terminal tryptophan, which is critical for activity (Luke *et al*., 2008; Rao *et al*., 2025). Functional tests of 34 representative sequences in HEK293T cells confirmed that the majority supported some level of StopGo activity, with an extrapolated estimate of 1700 active 2A peptides; however, this has yet to be verified (Rao *et al*., 2025). In addition, Li et al. (2026) compiled approximately 870,000 nonredundant viral nucleotide sequences and identified 908 exact matches and 1,557 near-matches to the expanded core motif (D/G/C/N)(V/I)ExNPGP. Analysis of upstream sequences revealed six distinct hexapeptide motifs, and functional assays confirmed that representative upstream motifs are required for StopGo activity. This study suggests that 2A peptide diversity extends beyond the current Class A and Class B classification (Li *et al*., 2026). The molecular basis for the functional differences between these classes of 2A peptides is not known.

StopGo elements are predominantly associated with single-stranded RNA viruses and select eukaryotic lineages but are rare in cellular genomes and absent from mammals, with only sporadic examples in certain eukaryotic retrotransposons (Odon *et al*., 2013; de Lima and Lanza, 2021; Li *et al*., 2026). Notably, despite functional compatibility with plant translation systems, StopGo elements were virtually absent from plant virus genomes, suggesting lineage-specific evolutionary constraints on 2A utilization (Rao *et al*., 2025; Li *et al*., 2026). Furthermore, to our knowledge, there have been no reports that 2A StopGo reactions are supported in prokaryotes (Donnelly *et al*., 2001; Rao *et al*., 2025). Together, these observations indicate that 2A-mediated recoding is predominantly a viral eukaryotic strategy for expanding coding capacity, with limited adoption in host genomes.

Although bioinformatic surveys have identified thousands of candidate 2A peptides, functional validation remains limited to small subsets tested individually by western blot or *in vitro* translation assays(Luke *et al*., 2008; de Lima and Lanza, 2021; Rao *et al*., 2025; Li *et al*., 2026). The extent to which computationally predicted 2A motifs across the virome encode functional StopGo activity has not been comprehensively determined. Furthermore, no comprehensive high-throughput functional mutagenesis screen has been applied to 2A peptides, and previous mutational analyses have relied on targeted substitutions within individual sequences, examining one or a few positions at a time (Doronina *et al*., 2008; Sharma *et al*., 2012; Kjaer and Belsham, 2018; Rao *et al*., 2025; Li *et al*., 2026). In this study, we developed a novel fluorescence-based high-throughput mammalian reporter that quantitatively measures 2A/StopGo activity across extensive libraries of natural and synthetic sequences, including candidates mined from viral genomes and metagenomic datasets. Using this platform, we identify key amino-acid determinants and residue boundaries governing StopGo activity across multiple viral 2A peptides and functionally rank diverse viral sequences in a mammalian translation context. Moreover, these analyses reveal previously unrecognized functional classes of 2A peptides and the functional 2A peptides associated with specific viral lineages. Finally, we provide a framework for elucidating how distinct peptide architectures engage the ribosomal exit tunnel to mediate peptidyl–tRNA hydrolysis and elongation resumption, as well as for tracing the evolutionary diversification of 2A-mediated translational recoding.

## MATERIALS AND METHODS

### Bioinformatic identification of 2A and 2A-like peptides

2A-peptides were identified from the Serratus assembly dataset (s3://serratus-public/assemblies/rdva_v0.2.fa.lz4), comprising assemblies derived from 58,557 NCBI SRA accessions representing diverse hosts and habitats (Edgar *et al*., 2022). Distinct profiles for classes A, B, and X, respectively, were employed to screen assembled contigs for 2A-peptide-like sequences (Figures 7 and 8). Contigs containing putative 2A-peptides were subsequently screened for RNA-dependent RNA polymerase (RdRp) sequences using the RdRp-scan pipeline (Charon *et al*., 2022) with default parameters, and for RdRp-palm sequences using PalmScan (Babaian and Edgar, 2022; Edgar *et al*., 2022) with default parameters. Contigs containing RdRp and/or RdRp-palm sequences were considered to originate from RNA viruses in the kingdom *Orthornavirae*.

Although some RNA virus-derived contigs may lack detectable RdRp or RdRp-palm sequences due to incomplete assembly, these contigs were retained for downstream analyses and taxonomic assignment using complementary taxonomic annotation pipelines, including geNomad v.1.9.0 (Camargo *et al*., 2024), the Contig Annotation Tool (CAT) v.4.6 (von Meijenfeldt *et al*., 2019), and Kaiju v.1.9.1(Menzel, Ng and Krogh, 2016), as described below.

The obtained 2A-peptide sequences, as well as the 2A-peptide-containing contigs, were deduplicated using SeqKit v.2.13.0 (Shen *et al*., 2016) with default parameters (rmdup -s). To assess the novelty of the 2A peptides identified in this study, the obtained 2A-peptide sequences were clustered together with previously reported 2A peptides (Rao *et al*., 2025; Li *et al*., 2026) using CD-HIT v.4.8.1(Li and Godzik, 2006). Clustering was performed at 100% amino-acid sequence identity across the alignment, with the alignment covering 100% of the shorter sequence (-c 1.0 -n 5 -G 1 -aS 1.0 -g 1 -d 0 -M 0). This criterion requires the shorter sequence to be fully aligned to the longer sequence while allowing the longer sequence to contain additional residues, thereby accommodating differences in sequence length among studies (e.g., the flanking residues included in the ∼100 amino-acid sequences reported by Li et al. (2026) (Li *et al*., 2026). The 2A-peptide sequences that did not cluster with any previously reported sequence under these criteria were considered novel.

### Taxonomy assignment

The taxonomy of 2A-peptide-containing contigs was determined using five complementary classification approaches. First, RdRp-palm-based classification (Edgar *et al*., 2022) was performed by searching RdRp-palm sequences of the 2A-peptide-containing contigs against PALMdb (released 26 April 2023; https://github.com/ababaian/palmdb) using DIAMOND v2.0.15 (Buchfink, Reuter and Drost, 2021) blastp (parameters: -d otu_centroids.fa.dmnd --ultra-sensitive --masking 0 -e 1e-5 --max-target-seqs 1). Second, the RdRp-scan pipeline (Charon, 2022) was employed to classify RdRp-containing contigs by searching translated contig sequences against the RdRp-scan_0.90 database using DIAMOND v2.0.15 (Buchfink, Reuter and Drost, 2021) blastx (parameters: -d RdRp-scan_0.90.dmnd --very-sensitive --matrix BLOSUM45 -e 1e-5 --min-orf 600 --max-target-seqs 1). Third, 2A-peptide-containing contigs were further analysed using geNomad v.1.9.0 (Camargo *et al*., 2024) to identify and classify mobile genetic elements, including various types of viruses. Fourth, the CAT pipeline v.4.6 (von Meijenfeldt *et al*., 2019) was employed with default parameters for broad taxonomic classification. Fifth, Kaiju v.1.9.1 (Menzel, Ng and Krogh, 2016) was used with the NCBI nr database to provide additional taxonomic assignments across all organisms.

Taxonomic assignments for each 2A-peptide-containing contig were then integrated using a hierarchical priority order: RdRp-palm-based classification, RdRp-scan, geNomad, CAT, and Kaiju. When conflicting assignments occurred at lower taxonomic ranks, such as the family level, but a consistent higher-level classification, such as the order level, was supported across methods, the shared higher-level taxonomy was retained. Finally, taxonomic assignments of RNA viruses were further manually refined and curated through phylogenetic analyses of RdRp-palm sequences as well as the virus annotation, as described below. The reverse-transcribing elements (RTEs) were defined when detecting the reverse transcriptase in the genome/contig sequences.

### Phylogenic analysis

To elucidate the evolutionary relationship of 2A-peptide-containing contigs to other RNA viruses, we conducted phylogenetic analysis of the RdRp-palm sequence from 2A-peptide-containing contigs, and representative RNA viruses from the NCBI RefSeq (release 216). The RdRp-palm amino-acid sequences were aligned using MAFFT v.7 (Katoh *et al*., 2002) with default parameters. Then, the multiple alignments were trimmed using Clipkit v.1.4.1 (Steenwyk *et al*., 2020) (parameters: -m kpic-smart-gap). Phylogenies using maximum likelihood (ML) were conducted using these multiple alignments. The construction of the ML phylogeny was executed using IQ-TREE v.2.2.0.3 (Nguyen *et al*., 2015) with 1,000 bootstrap replicates (parameters: -m MFP -B 1000 -bnni). This analysis used the optimal model and gamma-distributed substitution rates determined and implemented by IQ-TREE.

To elucidate the 2A-peptides evolutionary history, a phylogenetic analysis was performed using the identified 2A-peptide sequences from the 2A-peptide-containing contigs, as described above. These 2A-peptide sequences were then aligned using MAFFT v.7, trimmed with Clipkit v.1.4.1, and used to construct ML phylogenies with IQ-TREE v.2.2.0.3 employing the same model selection and bootstrap settings as above.

### Virus annotation

Putative coding sequences (CDSs) in 2A-peptide-containing contigs were predicted using the ORF prediction tool implemented in Geneious Prime® v.2026.1.1 (Biomatters Ltd., Auckland, New Zealand), with following parameters (genetic code = Standard; minimum size = 600 bp). Predicted ORFs were used for downstream protein-domain annotation and genomic context analyses. Annotation of these CDSs, including the identification of conserved domains within the polyprotein, was carried out using InterProScan version 5.59-91.0 (Jones *et al*., 2014) employing default parameters. The InterProScan pipeline integrates a range of protein databases, including Pfam, NCBI SUPERFAMILY, Conserved Domains Database (CDD), PANTHER, ProSiteProfiles, ProSitePatterns, and CATH-Gene3D, enabling a detailed exploration of the protein profiles and domain architectures inherent to the viruses. To further identify remote homologues and infer putative structural features, predicted protein sequences were searched against the PDB70 database using HHpred (Soding, Biegert and Lupas, 2005).

### Reverse-transcribing elements (RTEs) classification

Genome or contig sequences encoding a detectable reverse transcriptase (RT) domain were operationally defined as reverse-transcribing elements (RTEs). RT-containing contigs were first classified using TEsorter v.1.5.1 (Zhang *et al*., 2022) and DANTE v.0.1.9 (Novak *et al*., 2024). TEsorter was used to assign retrotransposon lineages based on profile-HMM searches against REXdb (viridiplantae_v4.0 + metazoa_v3.1) and conserved domain organization (parameters: -prob 0.9 -cov 30 -eval 1e-5 -score 1), whereas DANTE was used to provide complementary domain-based classification against the Viridiplantae v.4.0 and Metazoa_v3.1 databases using default settings (Neumann *et al*., 2019). To facilitate classification of RT-containing contigs that were incompletely or ambiguously resolved by these tools, custom script (https://github.com/kevinzhongxu/RTEs_classifier) were used to integrate conserved protein-domain annotations (see below) and genome/domain architecture information. This additional curation was used to distinguish major categories including non-LTR retrotransposon-like, LTR retrotransposon-like, retrovirus-related, group II intron-like, telomerase-like, CRISPR-RT-like, and other RT-containing elements. Final classifications were assigned by integrating evidence from TEsorter, DANTE, and domain-architecture information, with conflicting or insufficiently supported cases conservatively designated as ambiguous or unclassified RT-containing elements.

### Analysis of 2A-peptide StopGo activity across viral taxa

Normalized activity scores were compared among viral taxonomic groups at each hierarchical taxonomic rank (e.g., genus, family, order, class, and phylum) using nonparametric statistical methods implemented in R v.4.5.2 (Team, 2000). Differences among viral groups were evaluated using two-sided Wilcoxon rank-sum tests, with *P* values adjusted using the Benjamini–Hochberg false discovery rate (FDR) procedure. Differences in normalized 2A-peptide activity across viral taxonomic groups were further assessed using permutational multivariate analysis of variance (PERMANOVA) with the adonis function in the R package vegan v.2.5 (Oksanen, 2013), using Bray–Curtis dissimilarities and 999 permutations. Post-hoc pairwise PERMANOVA comparisons were performed using pairwiseAdonis v.0.4 (Martinez Arbizu, 2017), and resulting *P* values were adjusted for multiple testing using the Benjamini–Hochberg false discovery rate procedure.

To identify taxa exhibiting unusually high 2A-peptide activity, one-versus-rest Wilcoxon rank-sum tests were performed at each taxonomic rank. For each taxon, normalized activity scores were compared with those of all remaining taxa combined using a one-sided Wilcoxon rank-sum test, with the alternative hypothesis that the focal taxon had greater activity than the background population, using the R package stats. Resulting *p-values* were adjusted for multiple testing using the Benjamini– Hochberg false discovery rate procedure. Effect size was quantified as the difference in median normalized activity score between the focal taxon and the background population, using the R package rstatix (Kassambara, 2019).

### Motif enrichment analysis

To identify sequence features associated with 2A-peptide activity, peptides were ranked by normalized 2A FRET activity score and partitioned into high- and low-activity groups using either the top and bottom 50% or the top and bottom 25% of sequences. Motif enrichment analyses were performed using DREME from the MEME Suite v.5.5.2 (Bailey *et al*., 2015), with high-activity peptides used as the positive set and low-activity peptides as the control set. Analyses were conducted using either full-length 2A-peptide sequences (26 amino acids) or the conventional C-terminal 20-amino-acid region.

Motif discovery was performed across multiple taxonomic levels, including individual 2A-peptides, genomes, genera, families, orders, classes, and phyla. Selected motifs associated with elevated activity were further characterized by examining their taxonomic distribution, genomic context, and sequence conservation using multiple-sequence alignments (as above in the Phylogenetic Analysis section) and sequence-logo analyses using R package ggseqlogo v.0.2.2 (Wagih, 2017).

### Statistical analysis and visualization

Unless otherwise stated, statistical analyses were performed in R v.4.5.2 (Team, 2000). Data processing and summarization were conducted using the tidyverse package suite (Wickham *et al*., 2019). Wilcoxon rank-sum tests were performed using functions from the stats and rstatix packages (Kassambara, 2019); the multiple-testing correction was performed using the Benjamini–Hochberg false discovery rate procedure, and statistical significance was defined as FDR-adjusted *P* < 0.05. PERMANOVA was performed using the R package vegan v.2.5 (Oksanen, 2013), and post-hoc pairwise PERMANOVA tests were performed using pairwiseAdonis v.0.4 (Martinez Arbizu, 2017). Sequence motif enrichment was conducted using DREME from the MEME Suite v.5.5.2 (Bailey *et al*., 2015). Visualizations were generated using ggplot2 v.4.0.2 (Wickham, 2011) and ggseqlogo v.0.2.2 (Wagih, 2017).

### Plasmids

The 2A-sensing FRET reporter (Myc-mClover-2A-mRuby-FLAG) contains mClover (donor, N-terminal Myc tag) and mRuby (acceptor, C-terminal FLAG tag) flanking the in-frame 2A peptide and was ordered from Twist Biosciences, cloned downstream of a CMV promoter in a lentiviral backbone (Addgene plasmid 131127). The 2A peptide is flanked by BamHI and EcoRI sites. Reporter variants were generated by replacing the wild-type 2A with mutant 2A sequences by restriction cloning with BamHI and EcoRI. Wild-type P2A (ATNFSLLKQAGDVEENPG↓P) and an inactive P2A double mutant (G18V/P19A; ATNFSLLKQAGDVEENPVA), which yields a 100% fused mClover–mRuby protein, were cloned as controls. Single-fluorophore constructs expressing mClover alone or mRuby alone were generated for compensation and gating. All constructs were verified by sequencing (Plasmidsaurus).

### Library construction

All libraries were ordered as oligonucleotide pools from Twist Bioscience, which were PCR-amplified with flanking BamHI and EcoRI restriction sites and ligated into the BamHI/EcoRI-digested lentiviral MYC-mClover-2A-mRuby-FLAG reporter. Ligation was performed with T4 DNA Ligase (New England Biolabs, Cat. No. M0202M) at a 1:3 vector-to-insert molar ratio in 1× T4 DNA Ligase Buffer and incubated at 16 °C for 16 h. Ligation reactions were purified and transformed into NEB stable competent E. coli (C3040H). Transformed cells were recovered in SOC at 30 °C for 2 hours and plated at multiple dilutions on LB-agar plates with ampicillin to calculate the coverage, and the rest were grown in LB broth. The total number of colonies was estimated to confirm at least ∼100× coverage of the library. Plasmid DNA was extracted by midiprep to generate the plasmid library used for lentiviral packaging.

### Lentivirus production

Lentiviral particles were produced in HEK293T cells by co-transfection with the lentiviral reporter construct and the packaging plasmids psPAX2 and VSV-G (pMD2.G) at a ratio of 1:1:1 (vector: psPAX2: VSV-G) using Lipofectamine 2000 (Thermo Fisher Scientific, Cat. No. 11668027) according to the manufacturer’s instructions. Plasmid DNA and Lipofectamine were each diluted in Opti-MEM (Gibco, Thermo Fisher Scientific, Cat. No. 31985062), combined, incubated at room temperature for 10–15 min, and added dropwise to the cells with gentle rocking. The medium was replaced with 10 mL fresh DMEM 24 h after transfection, and supernatant was collected 48 h after transfection and filtered through a 0.45 µm filter. Virus was concentrated using an Amicon Ultra Centrifugal Filter (100 kDa; UFC910096) by centrifugation at 3,000 rpm for 10–20 min at 4 °C; the flow-through was discarded, and the concentrated virus was aliquoted and stored at −80 °C. Lentiviral MOI was determined by transducing HEK293T cells with a dilution series of viral supernatants and analysing the percentage of cells expressing green and red fluorescence using flow cytometry. For each pooled library screen, cells were transduced at a scale that maintained at least 200× coverage of the library. The volume of virus needed was calculated from the lentiviral MOI, keeping the virus-to-cell ratio constant at an MOI of 0.3-04, and transduced populations were expanded and selected before FACS sorting.

### FRET analysis

FRET was measured on the CytoFLEX SRT cell sorter (Beckman Coulter) using the mClover (Ex: Em 488:525 nm), mRuby (560:610 nm), and FRET (488:610 nm) channels. Live cells were gated on forward and side scatter (FSC/SSC) and compensated for mClover and mRuby. Double-positive mClover + mRuby cells were gated first, and a gate was then applied to exclude false-positive FRET signals arising from direct excitation of mRuby by the 488 nm laser, and a second gate (FRET versus mClover) was set on the FRET-negative co-transfected mClover + mRuby population to define FRET-positive cells. Cells expressing mClover or mRuby alone were used as single-fluorophore controls, and the fused mClover–mRuby protein was used as the FRET-positive control, adapted from (Trümper *et al*., 2019). Low, Medium, and High fluorescence cell populations were analyzed by FRET and isolated.

### Cell sorting, library sequencing, and FRET activity scoring

For pooled library screens, transduced cells were sorted for FRET-positive (inactive 2A) and no-FRET (active 2A) populations on the CytoFLEX SRT cell sorter (Beckman Coulter), collecting at least 100× coverage of the library in each gate. Genomic DNA was extracted from each population using the QIAamp DNA Mini Kit (Qiagen). The 2A peptide sequence was amplified in a two-step PCR with Phusion polymerase master mix: the first PCR of 18 cycles using Illumina-adapter primers, followed by a second indexing PCR of 14 cycles seeded with 5% of the first-round product, adapted from (Rehfeld *et al*., 2023). The final products were purified by Zymo oligo cleanup and concentrator (D4060) and sent to Novogene for Illumina sequencing at a minimum of 100× coverage of the library. Reads were assigned to individual variants and normalized to the total read counts (individual variant reads / total number of reads). A per-variant FRET activity score was then calculated as No-FRET reads (No-FRET + FRET reads).

### Immunoblotting Analysis

For individual 2A variant validation, fragments were ordered as oligonucleotides (GenScript, Twist Bioscience, ThermoFisher) and cloned into the myc-mClover-2A-mRuby-FLAG reporter by Gibson assembly. Clones were miniprepped and confirmed by sequencing (Genewiz, Plasmidsaurus). Each construct was transfected into HEK293T cells using Lipofectamine 2000 according to the manufacturer’s protocol for 24 h. Cells were lysed in RIPA buffer with protease inhibitors, and protein concentration was determined by the Bradford assay. Equal amounts of lysate were resolved by 12% SDS-PAGE and transferred to a PVDF membrane (Amersham HyBond Cat. No.10600023). Membranes were blocked in 5% milk in TBST (0.2% Tween-20) and probed with mouse anti-FLAG (downstream product, DOWN) (Millipore Sigma Cat. No. F4042), rabbit anti-Myc (upstream product, UP) (Invitrogen Cat. No. A190-105A, 1:1,000), and mouse anti-β-actin (loading control) (Abcam #ab8224, 1:1,000). Blots were then incubated with LI-COR IRDye fluorescent secondary antibodies (IRDye 680 anti-mouse (926-68070) and IRDye 800 anti-rabbit (926-32211) at 1:5,000 and imaged on the Amersham Typhoon Biomolecular Imager. Bands corresponding to the fused (∼58.8 kDa), upstream (Myc-mClover, ∼30.7 kDa), and downstream (mRuby-FLAG, ∼28.0 kDa) products were quantified by densitometry in ImageQuant, and 2A activity was calculated as Down/(Down+FL) × 100%. All immunoblot quantifications are from three independent experiments (N = 3).

### 2A Sensitivity score

To define the 2A sensitivity score, we consider at each mutation the empirical logit transform of the FRET response with the Haldane-Anscombe correction (Weber *et al*., 2020) :

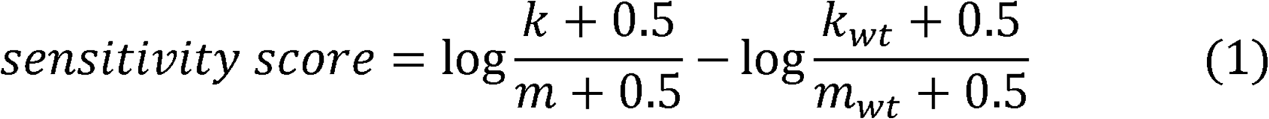

where *k* and *m* are the cleaved and uncleaved read counts, and *k_WT_* and *m_WT_* are the corresponding counts for wild type. The sum of cleaved and uncleaved counts greater than 10 were used. A zero-sensitivity score means wild-type cleavage efficiency, negative values indicate loss of cleavage, and a 2.30 sensitivity score drop equates to a tenfold reduction in the odds of cleavage.

### Nested Multiple Regression

Nested multiple regression was used to analyze determinants of the 2A sensitivity score. To assess the impact of physicochemical properties, and to compare F2A and P2A we considered the following model:

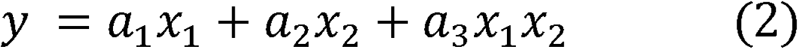

where y is the 2A sensitivity score*, x_1_* encodes for the position, and *x_2_* encodes for either the species, or different physicochemical properties of amino acids that were tested. The *a_i_*’s are the inferred parameters, where *a_1_* and *a_2_* account for independent effects and *a_3_* interaction between variables. The models were fitted using ordinary least squares (OLS). When testing for physicochemical properties, we used the Grantham distance and its components --polarity, molecular volume, and molecular composition (with values taken from (Grantham, 1974). The nested F-tests were then used to determine if the inferred *a_2_*and *a_3_* parameters are significant in the fit.

### Structural Visualization

The Cryo-EM structure of the F2A within the ribosomal exit tunnel reported by (Li *et al*., 2026) PDB: 9RHU was visualized using PyMOL (3.1) and the ribosomal exit tunnel was extracted following the pipeline outlined in (Yu *et al*., 2026), available on GitHub (https://github.com/bioshape-analysis/).

### Data availability statement

Sequences of 2A-peptides and 2A-peptide-containing contigs, as well as other supplementary materials, such as trees, alignments, and relevant codes, can be found in figshare (Baral, et al., 2026) via this link: https://doi.org/10.6084/m9.figshare.33359493. Custom scripts used for RTEs classification are available on GitHub at https://github.com/kevinzhongxu/RTEs_classifier.

## RESULTS

### Development of a 2A Peptide FRET-FACS Reporter for Single-Residue Mutagenesis Screening

Despite the widespread use of 2A peptides for polycistronic expression in mammalian cells, the contribution of individual residues to ribosomal skipping efficiency and the full diversity of naturally occurring active 2A sequences remain incompletely characterized (Sharma *et al*., 2012; Rao *et al*., 2025; Li *et al*., 2026). To address this gap and enable quantitative, high-throughput screening of 2A peptide activity in mammalian cells, we developed a novel fluorescence resonance energy transfer (FRET)–based reporter compatible with fluorescence-activated cell sorting (FACS) and deep sequencing (Figure 1A). The reporter consists of an N-terminal myc-tagged mClover donor and a C-terminal mRuby acceptor flanked by a FLAG tag, separated by an in-frame 2A peptide of interest (mClover-2A-mRuby). When the 2A peptide is functional, ribosomal skipping uncouples mClover from mRuby, producing two physically separated fluorophores and abolishing FRET (Figures 1B, 1C). When the 2A peptide is inactive, the two fluorophores remain tethered as a single fusion protein, generating FRET upon donor excitation. We evaluated FRET in mammalian cells expressing the mClover-2A-mRuby reporter. To optimize gating by FACS analysis, HEK293T cells stably expressing mClover or mRuby alone, both mClover and mRuby, or a fused mClover–mRuby protein were analyzed (Supplementary Figure S1A). We gated cells on forward and side scatter (FSC/SSC) and compensated for mClover and mRuby fluorescence to specifically evaluate FRET in double-positive mClover and mRuby and fused mClover–mRuby cells. We gated and sorted the double-positive mClover + mRuby cells into Low, Medium, and High fluorescence populations and evaluated FRET in each population (Supplementary Figures S1B, S1C). The fraction of FRET-positive cells was 63.4%, 74.0% and 83.9% in the expanded Low, Medium and High fluorescence cell populations, respectively. To benchmark the dynamic range of the reporter, we compared cells expressing the reporter with the wild-type P2A with an inactivating double mutant (G18V/P19A) (Figures 1B, 1C). We initially chose highly fluorescent cells for analysis as medium and lower fluorescent cells resulted in inconsistent and subpar subsequent FRET analysis (Supplemental Figure S1). Wild-type P2A yielded efficient ribosomal skipping, with greater than 90% of cells displaying no FRET and fewer than 10% displaying FRET (Figure 1B, left), whereas the G18V/P19A mutant produced almost exclusively FRET-positive cells (>99%; Figure 1B, right), confirming that the assay resolves active from inactive 2A peptides on a per-cell basis. Immunoblot analysis of cells expressing the reporters validated the FACS readout: wild-type P2A produced predominantly the cleaved downstream product (∼25 kDa, anti-FLAG) and the upstream product (∼30 kDa, anti-myc), whereas the G18V/P19A mutant accumulated as the uncleaved full-length fusion protein (∼58.8 kDa) (Figure 1C). Thus, we have established a robust mammalian FRET-based 2A-sensing reporter assay.

**Figure 1.**
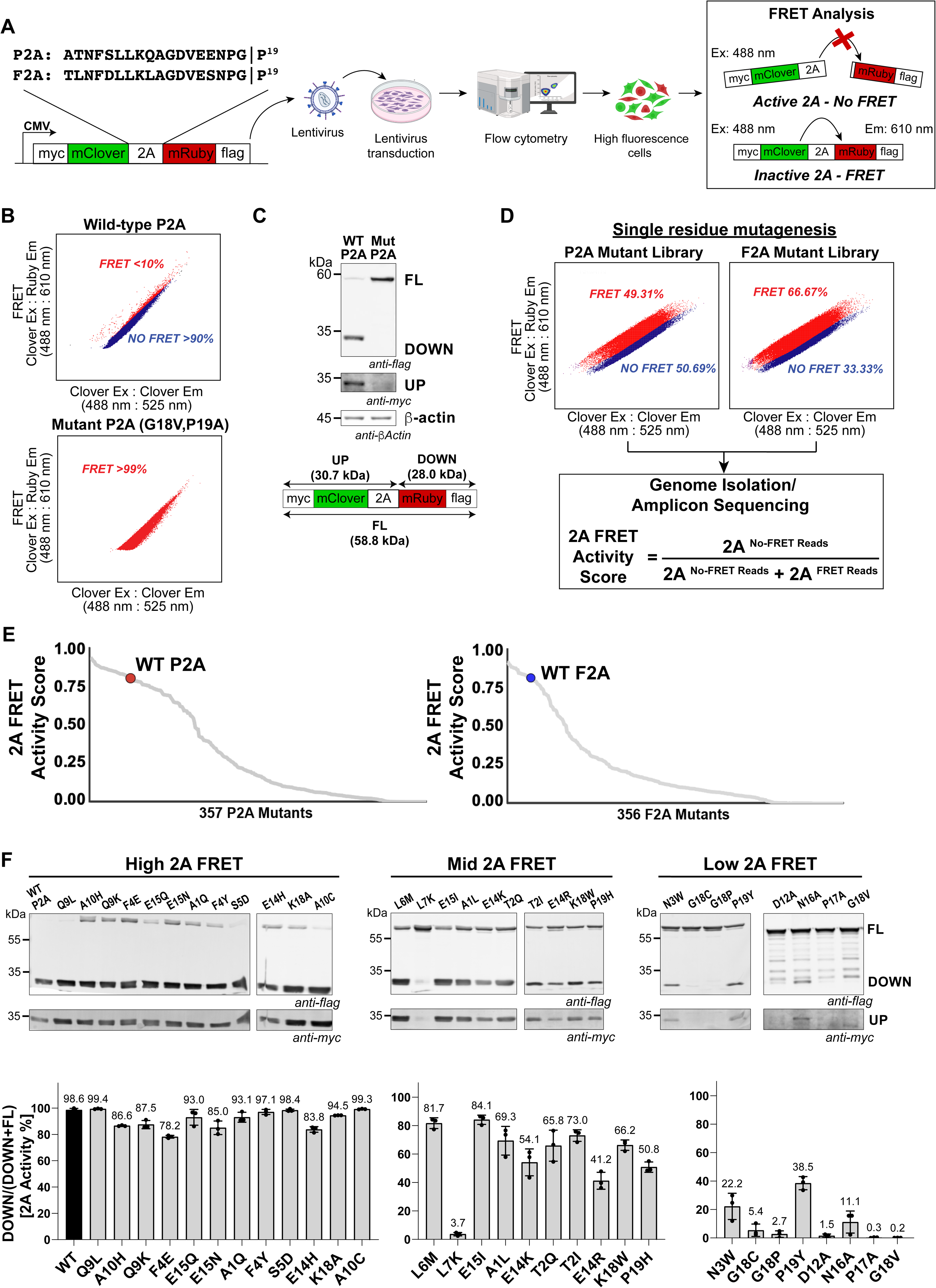
Development of a FRET-FACS mammalian reporter that senses 2A peptide stop-go activity. (A) Pipeline of 2A-sensing FRET-FACS. Schematic of the lentiviral dual-fluorescence reporter containing the mClover and mRuby FRET pair, with the 2A peptide cloned in-frame between the two fluorophores under a CMV promoter (myc-mClover-2A-mRuby-Flag). The P2A (ATNFSLLKQAGDVEENPG↓P¹□) and F2A (TLNFDLLKLAGDVESNPG↓P¹□) sequences are shown. Lentivirus-transduced HEK293T cells were selected with puromycin and sorted by flow cytometry based on high mClover and mRuby expression. When 2A is active, the ribosome separates the two fluorophores (no FRET); when 2A is inactive, the fluorophores remain in a single fused protein and produce a FRET signal (488 nm excitation, 610 nm emission). Cells displaying FRET (inactive 2A) or no FRET (active 2A) were sorted by FACS, genomic DNA was extracted, and the 2A-encoding region was amplified and sequenced. (B) FRET-FACS scatter plots (FRET Clover Ex: Ruby Em 488:610 nm versus mClover Ex: Em 488:525 nm) of HEK293T cells stably expressing wild-type P2A (top) or the inactive P2A double mutant G18V/P19A (bottom). Wild-type P2A shows less than 10% FRET-positive and more than 90% no-FRET cells, while the mutant shows more than 99% FRET-positive cells. (C) Immunoblots of HEK293T lysates expressing wild-type or G18V/P19A mutant P2A. The full-length fused protein (FL; myc-mClover-P2A-mRuby-Flag, 58.8 kDa) is shown at the top, the separated downstream product (DOWN; mRuby-Flag, 28.0 kDa) in the middle, and the upstream product (UP; myc-mClover, 30.7 kDa) below. (D) FRET-FACS scatter plots of the pooled P2A and F2A single-residue mutant libraries. The P2A library shows 49.31% FRET and 50.69% no-FRET, and the F2A library shows 66.67% FRET and 33.33% no-FRET. The sorted FRET and no-FRET populations were subjected to genomic DNA isolation and amplicon sequencing, and a per-variant 2A FRET activity score was calculated as 2A no-FRET reads / (2A no-FRET reads + 2A FRET reads). (E) Ranked 2A FRET activity scores for 357 P2A mutants (left) and 356 F2A mutants (right). Wild-type P2A (red) and wild-type F2A (blue) are highlighted at their respective rank positions. (F) Immunoblots of selected P2A mutants ranked by FRET activity, drawn from the high (left), mid (center), and low (right) FRET activity regions. 2A activity was quantified from immunoblot band intensities measured in ImageQuant as Down/(Down + FL), where higher values indicate greater 2A efficiency. Error bars represent average ± s.d.

We applied the FRET-FACS 2A-sensing reporter to map the contribution of each residue to 2A activity using single-residue saturation mutagenesis libraries spanning all 19 positions of P2A and F2A and plotted the resulting activity scores by position (Figure 2A, 2B). Saturation single-residue mutagenesis libraries of P2A and F2A (∼400 single mutations for each 2A peptide) were cloned into the reporter, packaged into lentivirus, and transduced into HEK293 cells. Following high fluorescence cells FACS sorting, the libraries displayed broad distributions of FRET versus non-FRET populations (Figure 1D), consistent with the expectation that single-residue substitutions produce a continuum of 2A activities. Genomic DNA was isolated from FRET and no-FRET cell populations, and the 2A-encoding region was identified by amplicon sequencing. For each variant, a 2A *FRET activity score* was computed as the ratio of no-FRET reads to total reads (no-FRET + FRET). This score, which is scaled from 0 (fully inactive) to 1 (fully active), provides a quantitative readout of 2A peptide function across thousands of variants in a single experiment.

**Figure 2.**
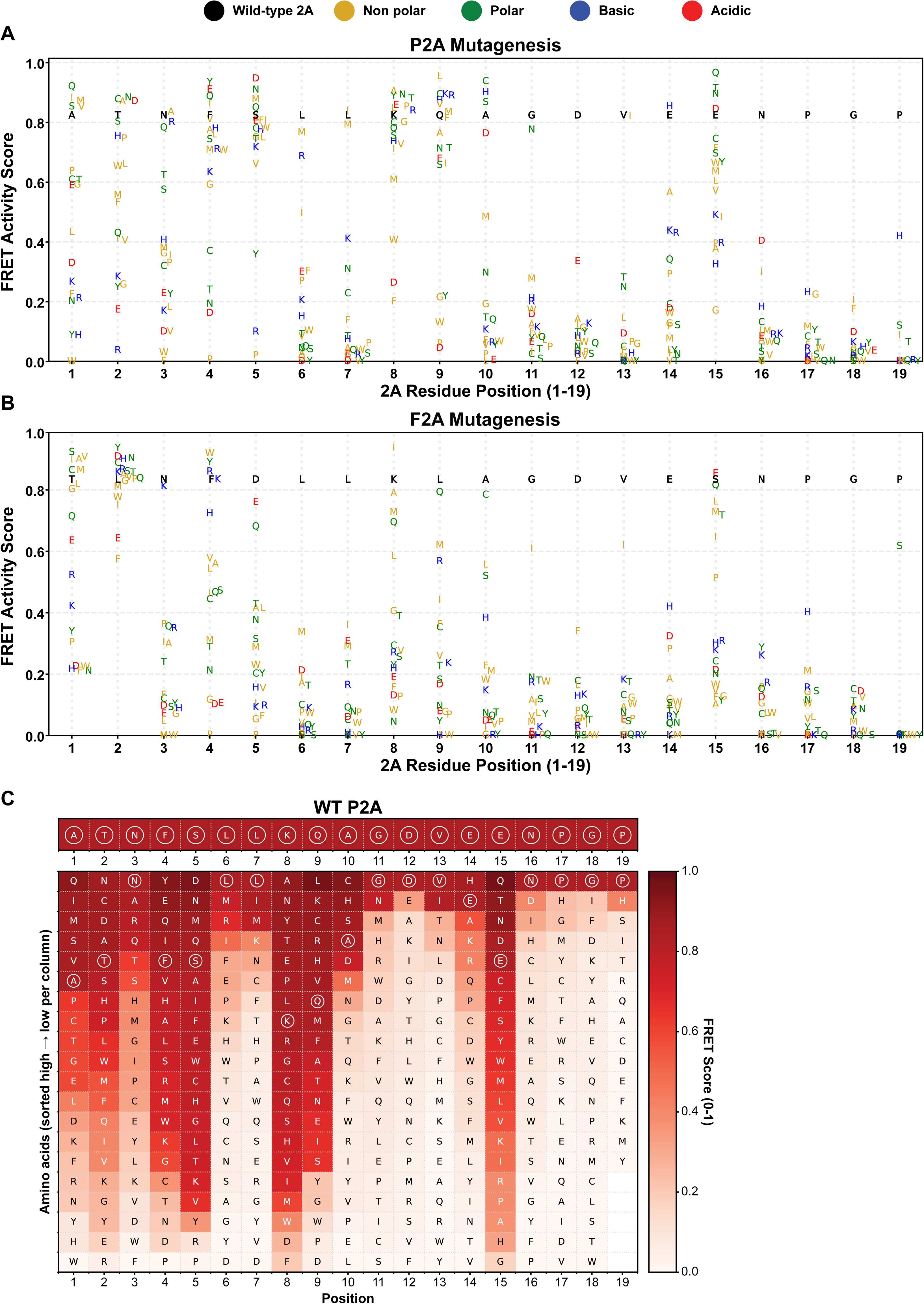
Scatter plot of relative FRET activity scores for all single-residue mutagenesis across the 2A peptide. (A–B) Scatter plots of position-specific single amino-acid mutagenesis effects on P2A and F2A activity. Each letter represents the activity of a single amino-acid substitution at the indicated position in the 2A peptide. The x-axis shows the 2A residue position (1–19), and the y-axis shows the relative FRET activity score for each mutant. Different colors represent distinct amino-acid characteristics, allowing visualization of position-specific tolerance and substitution effects across the peptide. Wild-type residue positions are shown in black for reference. (C) Heat map of FRET activity scores for amino-acid substitutions across the P2A peptide. The x-axis shows the residue position in P2A (1–19), and the y-axis shows the substituted amino acids, sorted from high to low activity score within each column. The color scale indicates the FRET score (0–1), with dark to light red representing higher to lower 2A FRET activity, respectively. The wild-type P2A sequence is shown at the top for reference.

From the single residue mutagenesis experiment, both peptide libraries displayed a range of FRET activities ranging from 0-99% (Figure 1E). To validate the single-mutagenesis FRET activities, select 2A mutants were selected by binned high, mid, and low FRET 2A activity scores and individually assessed for StopGo activity via immunoblotting of the N-myc and C-FLAG tagged reporter proteins following transfection into HEK293 cells (Figure 1F). StopGo activity for each mutant was quantified as the percentage of downstream protein (DOWN) detected by anti-FLAG relative to the combined total of downstream and full-length protein. In general, FRET activity bins broadly correlated with immunoblotting outcomes (Figure 1F). Wild-type P2A and F2A exhibited ∼99% and ∼75% StopGo activity, respectively; high-FRET P2A mutants ranged from 78–99% and high-FRET F2A mutants from 68–73%. Intermediate- and low-FRET P2A mutants showed correspondingly reduced activities of 41–82% and 0–39%, respectively. There were exceptions such as L7K that resulted in very low 2A activity despite binned in the mid-FRET group (Figure 1F). Despite this exception, these results demonstrate that FRET-FACS faithfully reports 2A StopGo activity.

The mutagenesis analysis of both P2A and F2A peptides displayed an asymmetric pattern of StopGo activity: the N-terminal domain was broadly permissive and the C-terminal domain progressively constrained (Figures 2A-2C). Positions 1–5 of the 2A peptides accommodated substitutions across all chemical classes, with many retaining near or relatively high wild-type activity (FRET score > 0.6) (Figures 2A-2C). Tolerance collapsed sharply closer to the conserved D-(V/I)-E-x-N-P-G-P motif. Consistent with these assignments, no substitution at conserved D12, E14, or N16 reached a FRET 0.6 threshold in either peptide, with the single exception of E14H in P2A (0.85) (Figures 2A, 2B). The hydrophobic-cluster leucines, L6 and L7, and the C-terminal -NPGP residues, were similarly intolerant in both peptides, that resulted in reduced 2A FRET activity to <0.2, in line with previous reports (Sharma *et al*., 2012; Li *et al*., 2026). Furthermore, mutations of G11, in general, also abolished 2A FRET activity in both peptides suggesting the importance of G11 for 2A reactions, however, G11 in both P2A and F2A tolerated N and I substitutions (FRET > 0.6). Although residue changes were more tolerant near the N-terminus of the 2A peptides, there were some residue changes that had similar properties. For instance, F4 of F2A, a peripheral aromatic, tolerated only aromatic and basic residues (F4 to W, Y, R, K, H resulted in 0.72-0.92 2A activity scores). V13 of F2A tolerated only β-branched I (2A activity score 0.62), and the same was true for V13 to I in P2A (2A activity score 0.82). The variable position 15, was the most permissive C-terminal position in both peptides and displayed tolerances to similar amino acid changes (F2A S15 to E, Q, L, M, T, I all > 0.6 activity; P2A E15: 11 substitutions including S, Q, L, M, T, > 0.6 activity). Interestingly, there were residue changes at N3, K4, and Q9 of P2A and F4, K8, and S15 of F2A that can be partitioned into active vs inactive 2A residues.

The FRET-FACS screen not only pinpointed the key positions but assigned each a quantitative, physicochemical basis for its effect on 2A StopGo activity. The strongest evidence for context-dependence came from a single comparison: the same residue at the same position behaved differently in P2A and F2A. Although the two peptides share the -DxExNPGP core and identical residues at several upstream positions, their tolerance profiles diverged sharply. At position 8, F2A permitted only four small-to-medium substitutions (I, A, M, Q) while bulky, charged, or aromatic residues abolished activity, whereas the identical lysine in P2A tolerated sixteen substitutions spanning every chemical class (Figures 2A, 2B). To compare the two peptides systematically, we assigned each position a sensitivity score—the mean loss of StopGo activity (in FRET log-odds) across all 19 substitutions, where a lower score denotes greater loss upon substitution (Figure 3A; Methods). Sensitivity spanned −7.1 to 0 log-odds in F2A and −6.4 to −0.2 in P2A, decreasing toward the C-terminal core, where essentially no substitution is tolerated in either peptide, while the N-terminal half remained broadly permissive. The two peptides diverged most at positions 5, 8, and 9 (S, K, Q in P2A; D, K, L in F2A), which were locally the most sensitive and thus the most predictive of activity. Because the side chains are identical at shared positions yet their tolerance differs, these differences reflect the local structural environment rather than the residue itself and thus pointing to context-specific roles for these residues within the ribosome exit tunnel.

**Figure 3.**
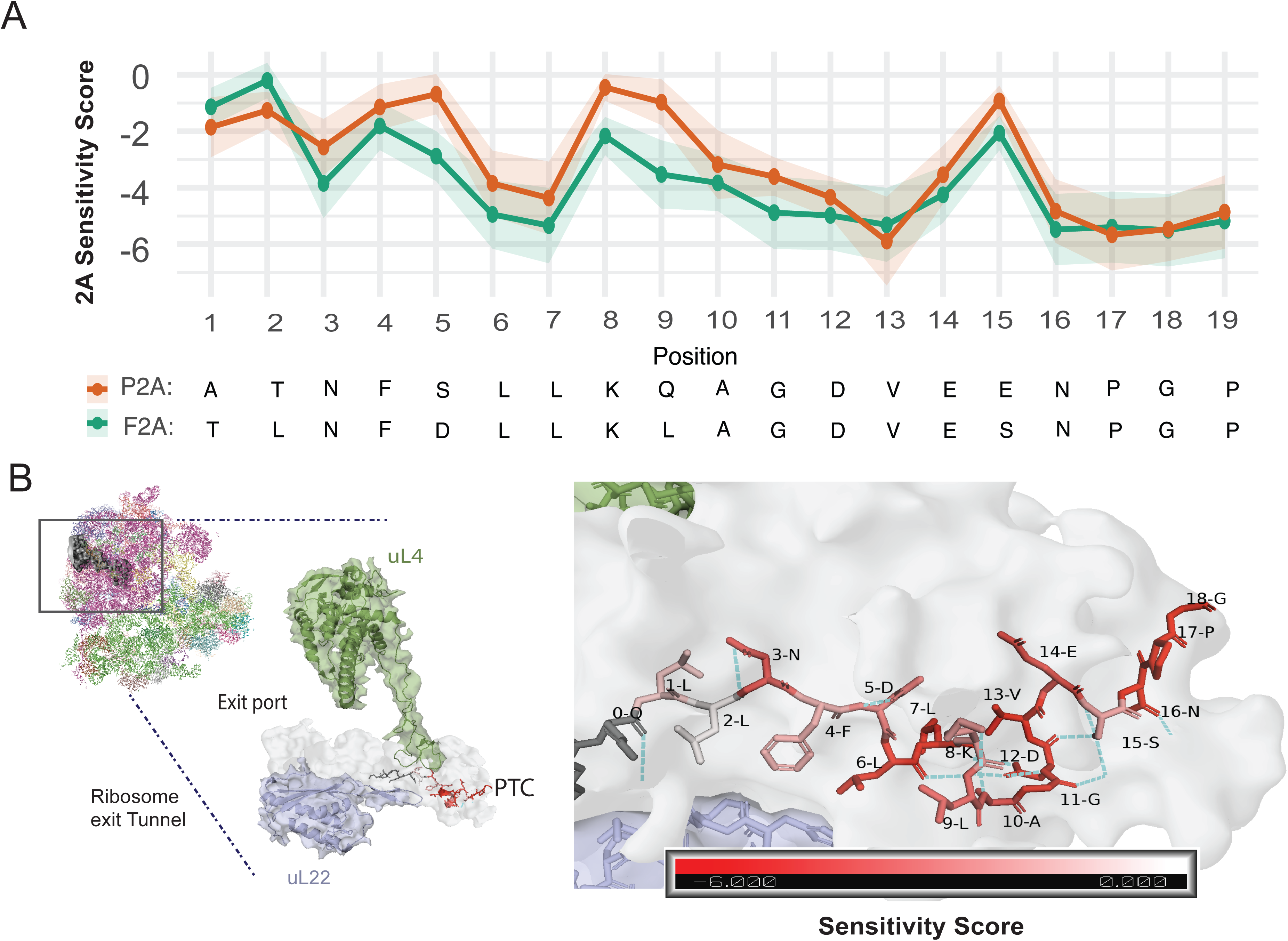
2A Positional sensitivity and structure-function association of the F2A peptide with the ribosome exit tunnel. (A) Positional sensitivity score for P2A and F2A. (B) Cyro-EM structure of F2A complexed with the rabbit ribosome (PDB: 9RHU) is shown on the left, with the ribosomal exit tunnel mesh and the constriction proteins of the ribosomal exit tunnel (uL22 and uL4). On the right, F2A is color-coded by sensitivity score, according to the color bar shown at the bottom.

We next performed statistical analyses on the sensitivity score at each point mutation (equation (1) in Methods) to assess the contribution of position, species and later physicochemical properties. First, we ran nested multiple regression with position and species (see Methods) and found significant effect (F value =14.2 p=0.000177), suggesting that subsequent analysis of physicochemical properties should be done separately for P2A and F2A. We therefore separately ran a multiple regression analysis for F2A and P2A, with position and physicochemical properties, captured by the Grantham matrix and its components (see Methods). We found for P2A a significant independent effect for Grantham distance and molecular volume (F=7.32 p = 0.007138, F=11.09 p=0.00096, respectively), while F2A showed significant independent effect with polarity only (F=3.93 p=0.0071). The interaction coefficient was significant only in F2A for Grantham and composition (F= 4.84 p=0.029, F=6.37 p=0.012, respectively).

To place the sensitivity scores in a structural context, we analyzed the cryo-EM structure of a mammalian ribosome stalled on F2A (Li *et al*., 2026) and examined the nascent chain within the exit tunnel (Fig 3B; Methods). F2A occupies the region between the peptidyl transferase center (PTC) and the constriction site formed by uL4 and uL22 (Dao Duc *et al*., 2019): of the 19 residues resolved, residues 4 through 19 lie between the PTC and the constriction, with residues 5–7 forming a short helical segment (Figure 3B). The three most sensitive positions (residues 4, 9, and 15; Figure 3A) each make a defined tunnel contact: residue 4 π-stacks with rRNA, Leu9 hydrogen-bonds within the α-helical backbone, and residue 15 hydrogen-bonds to the tunnel wall. These positions are distinguished by their structural contacts rather than by residue size or volume, consistent with our variance decomposition, indicating that sensitivity tracks specific contacts more than global property changes. The sharp drop in sensitivity across residues 5 to 9 (Figure 3A) coincides with the helical segment, linking the helix to StopGo activity. No cryo-EM structure of P2A is currently available; whether the same helical segment forms in P2A remains open.

### Delineation of the boundaries of functional 2A peptides

The N-terminal residues of 2A peptides are generally poorly conserved, yet studies indicate that N-terminal sequence and length can promote 2A activity (Sharma *et al*., 2012; Minskaia et al., 2013;Rao *et al*., 2025). To investigate this more thoroughly, we generated a deletion library of two 2A peptides, P2A and the Cricket paralysis virus 2A (C2A) (26 amino acids each) using the FRET 2A-sensing reporter. P2A and C2A represent high and low StopGo activity (∼90-95% and ∼50-60% 2A efficiency), respectively. We systematically introduced deletions from the N-terminus and at internal positions upstream of the conserved C-terminal domain. P2A FRET activity was maintained upon deletion of up to 9 N-terminal residues but dropped precipitously with further deletions of 10–15 residues (Figure 4). In contrast, the majority of internal deletions reduced P2A FRET activity. Notably, deletion of a single internal leucine or a ten-residue internal sequence still preserved relative P2A FRET activity. For C2A, which displays a relatively low baseline FRET score (∼0.2), progressive N-terminal deletions broadly maintained FRET activity, while larger N-terminal and internal deletions yielded FRET activities of 0–0.1 (Supplemental Figure S4). Of note, deletion of 12 internal residues resulted in an elevated P2A FRET score. To further delineate the boundaries of 2A peptide function, we systematically tested chimeric P2A/C2A sequences (Figure 4). Using P2A as the backbone, replacement of its six N-terminal residues with the corresponding C2A sequence retained 2A FRET activity, whereas more extensive C2A substitutions into P2A progressively reduced activity. Using C2A as the backbone, substitution of up to 12 N-terminal residues with P2A sequence maintained low FRET activity (<0.2), but replacement of 13 or more residues dramatically increased FRET activity. Taken together, both deletion and chimeric analyses converge in the central region of P2A and C2A as the inflection point between active and inactive 2A FRET activity.

**Figure 4.**
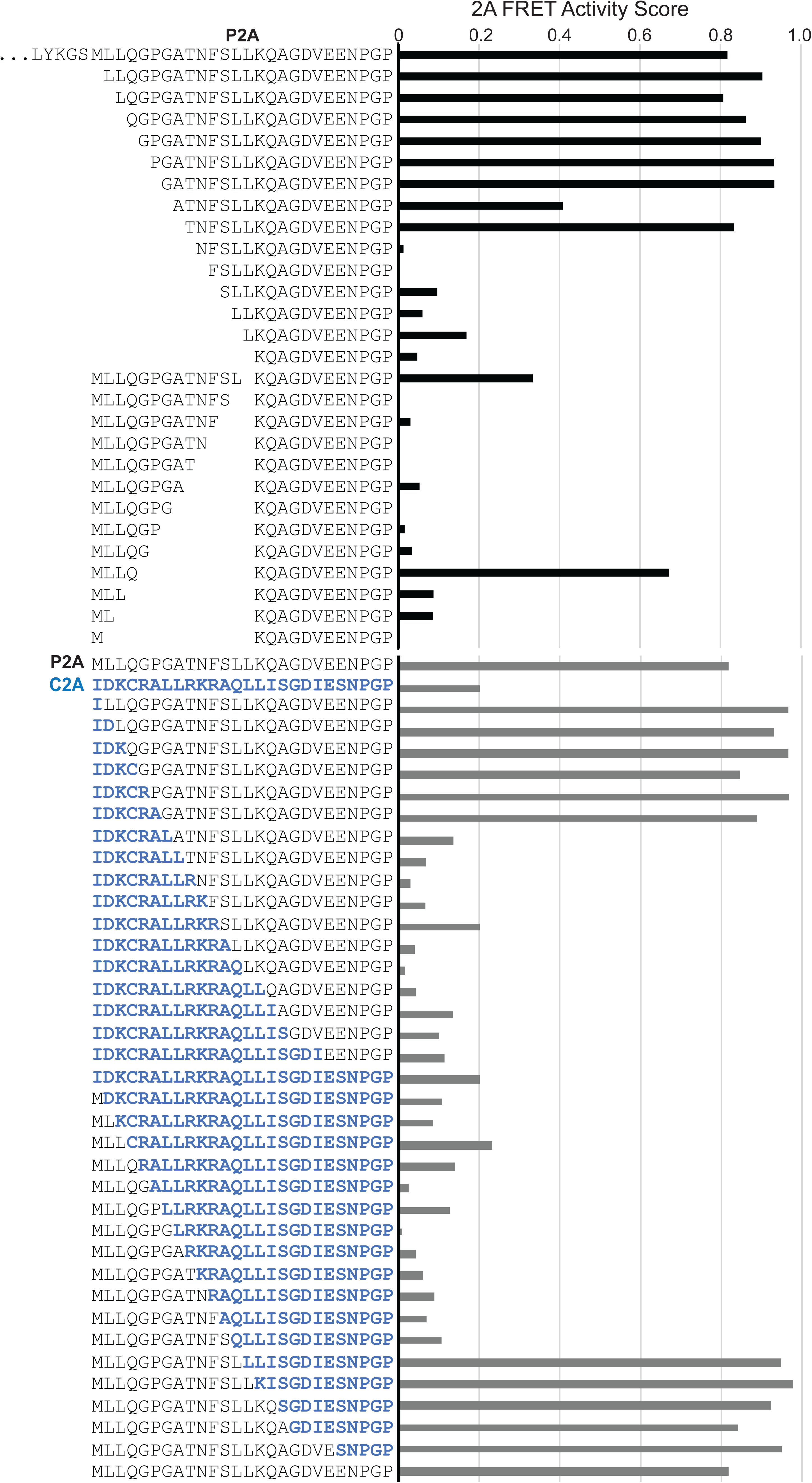
Deletion and chimeric libraries define the functional boundaries of 2A peptides. 2A FRET activity scores (0 = inactive, 1 = active) measured by FRET-FACS for deletion and chimeric variants of P2A and C2A. N-terminal and C-terminal deletions of the indicated 2A peptides. Each bar shows the 2A FRET activity score for the sequence shown on the left, with wild-type P2A at the top. Chimeric P2A/C2A peptide 2A activities are shown with P2A residues in black and C2A residues in blue. 2A FRET activities are shown average ± s.d.

### 2A StopGo Activity Is Tolerant of Residue Substitutions at Proline 19

Although the FRET-FACS approach broadly reflected StopGo activity, inspection of the P2A saturation mutagenesis dataset revealed that many P19 substitutions either failed to recover during cell sorting or returned too few sequencing reads to be reliably scored (Figures 2A-2C, Supplemental Data 2). To fully characterize P19 mutations on P2A activity, we cloned each of the 19 individual P19 substitutions and assayed them in HEK293T cells, rabbit reticulocyte lysate (RRL), and wheat germ extract (WGE) (Figures 5A-5C). In cell-free extracts, we monitored translation by [35S]-methionine incorporation of the upstream mClover (UP), the downstream mRuby (DOWN) and full-length (FL) polyprotein by SDS-PAGE analysis. Quantification of the radioactive proteins showed that the reporter containing the wild-type P2A resulted in >99% 2A activity. In contrast, all P19 substitutions in general reduced 2A activity compared to the wild-type P2A (Figure 5A). However, P19 substitutions to D, E, F, G, H, K, M, Q, S, T, V, W, and Y still retained >80% 2A activity in both RRL and WGE extracts, indicating that P19 of 2A is not absolutely essential for StopGo activity (Figure 5A, 5C). These results are in line using F2A-containing reporters in RRL (Li *et al*., 2026). In contrast, immunoblot analysis of HEK293T cells transfected with 2A-containing reporters, all P19 substitutions led to a 40–100% reduction in 2A activity from quantifying the DOWN/(DOWN+FL) immunoblots (Figures 5B, 5C). While no P19 substitution fully matched wild-type activity (P19, 2A activity ∼99%), several mutants retained substantial activity (e.g., P19W, P19M, P19E, P19Q), whereas others such as P19 to I, K, L, and R were severely impaired. Specifically, P19 substitutions to I, K, L, and R were particularly severe, yielding little detectable downstream product (DOWN) by immunoblotting, yet the upstream product (UP) was readily detected at levels comparable to other P19 mutants (Figures 5B, 5C). A potential mechanistic explanation for this asymmetry may be due to the N-end rule: ribosomal skipping produces a downstream protein whose new N-terminal residue is the residue immediately C-terminal to the skip site, which in the P19K and P19R mutants is lysine or arginine, respectively. Both lysine and arginine are primary destabilizing residues recognized by UBR-family E3 ubiquitin ligases of the N-end rule pathway (Hwang *et al*., 2010; Kim *et al*., 2021). To test this, HEK293T cells were treated with the proteasome inhibitor MG132 prior to transfection of the 2A-containing reporter, and 2A activity monitored by immunoblotting. MG132 treatment rescued downstream protein expression leading to 2A activity from near-zero to 6.4% for P19K and from 0.4% to 24.0% for P19R, while wild-type P2A remained similar (92.8% to 98.1%), confirming that its loss was in part due to N-end rule-mediated proteasomal degradation (Figures 5D, 5E). Together, these results demonstrate that 2A peptide activity can be supported by residues other than P19 *in vitro*, but that N-end rule degradation masks this tolerance in cells.

**Figure 5.**
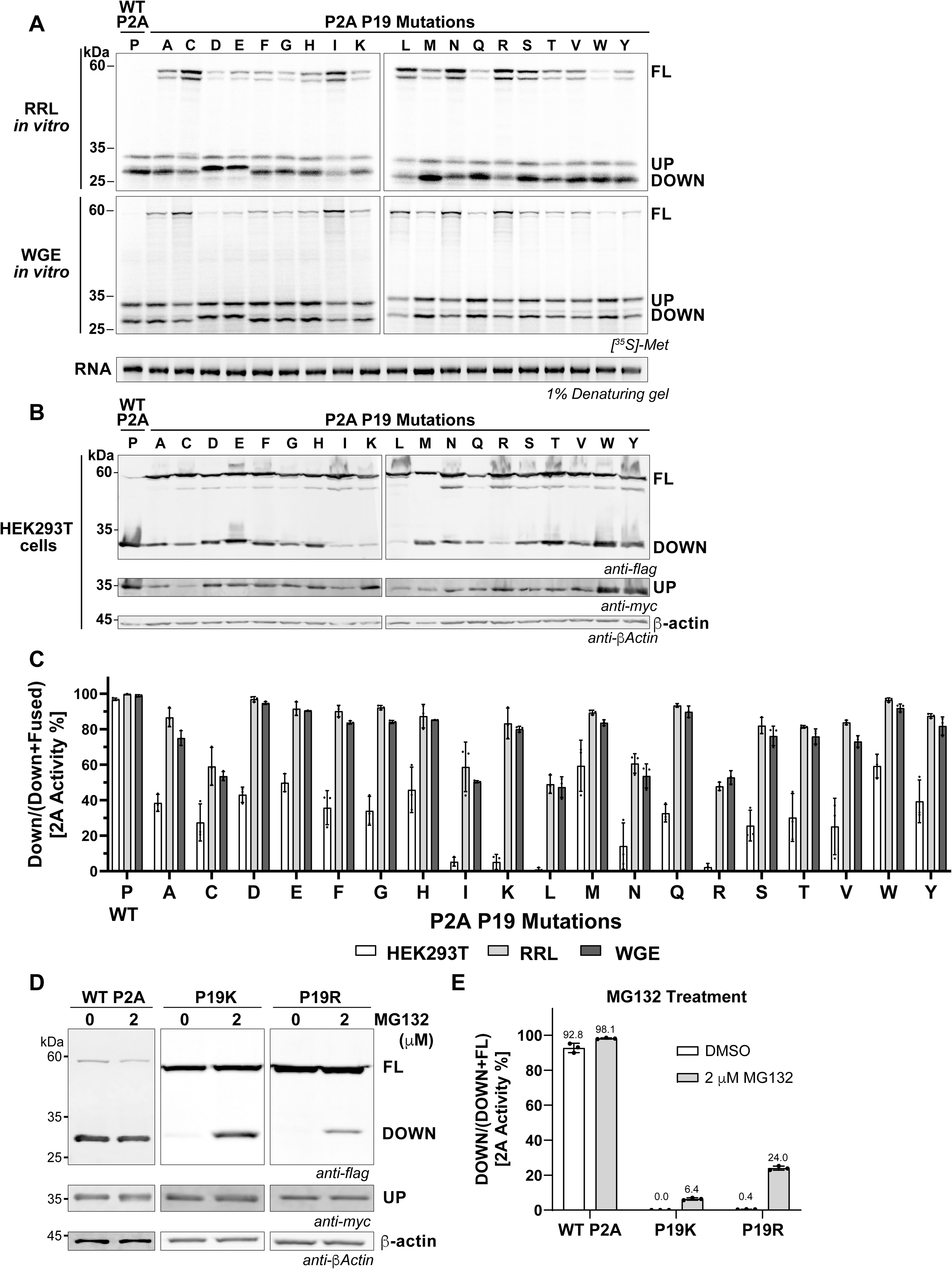
Evaluation of the effect of Proline 19 (P19) mutations on P2A activity. (A) Immunoblots of transfected reporter constructs containing P2A P19 mutants in HEK293T cells. Plasmid constructs containing the indicated P19 mutations were transfected into HEK293T cells for 24 hours followed by immunoblotting. (B) Autoradiographs of in vitro transcribed reporter RNAs containing P2A P19 mutants translated in rabbit reticulocyte lysate (RRL, top) and wheat germ extract (WGE, middle) cell-free systems with [³□S]-methionine and reactions were loaded and analyzed by SDS-PAGE analysis. (C) Quantification of P19 mutant activity from (A) and (B) calculated as relative 2A activity % = Down/(Down + FL). Error bars represent average ± s.d. (D) Immunoblots of HEK293T cells transfected with reporter constructs (24 hours) containing wild-type P2A, P2A P19K, or P2A P19R treated with DMSO (0) or 2 µM MG132. Error bars represent average ± s.d.

### Identification of functional virome-derived 2A peptides

Having validated the 2A-sensing FRET-FACS platform, we next applied it to identify functional viral 2A peptides using a virome-derived sequence library (Figure 6). To establish the evolutionary breadth of class A 2A peptides, we first screened the Serratus petabase-scale assembly resource, comprising sequences derived from 58,557 NCBI Sequence Read Archive (SRA) datasets spanning diverse hosts and ecosystems. We searched the viral Serratus.io database (Edgar *et al*., 2022) for genomes containing Class A 2A peptides using the canonical (G/H)D(I/V)ExNPGP motif. We identified 1,717 putative Class A 2A peptide sequences distributed across 1,547 nonredundant contigs (unique at the nucleotide level) (Supplementary Data 1). Among these, 1024 (66.2%) belonged to RNA viruses within the kingdom *Orthornavirae*, whereas 330 (21.3%) were associated with reverse-transcribing elements (RTEs), predominantly non-LTR retrotransposon-like sequences based on the genomic architecture and reverse-transcriptase content. The remaining 193 contigs (12.5%) represented unclassified viral sequences and/or sequences putatively derived from eukaryotic or prokaryotic hosts, consistent with previous investigation^1^ (Supplemental Figures S5A).

**Figure 6.**
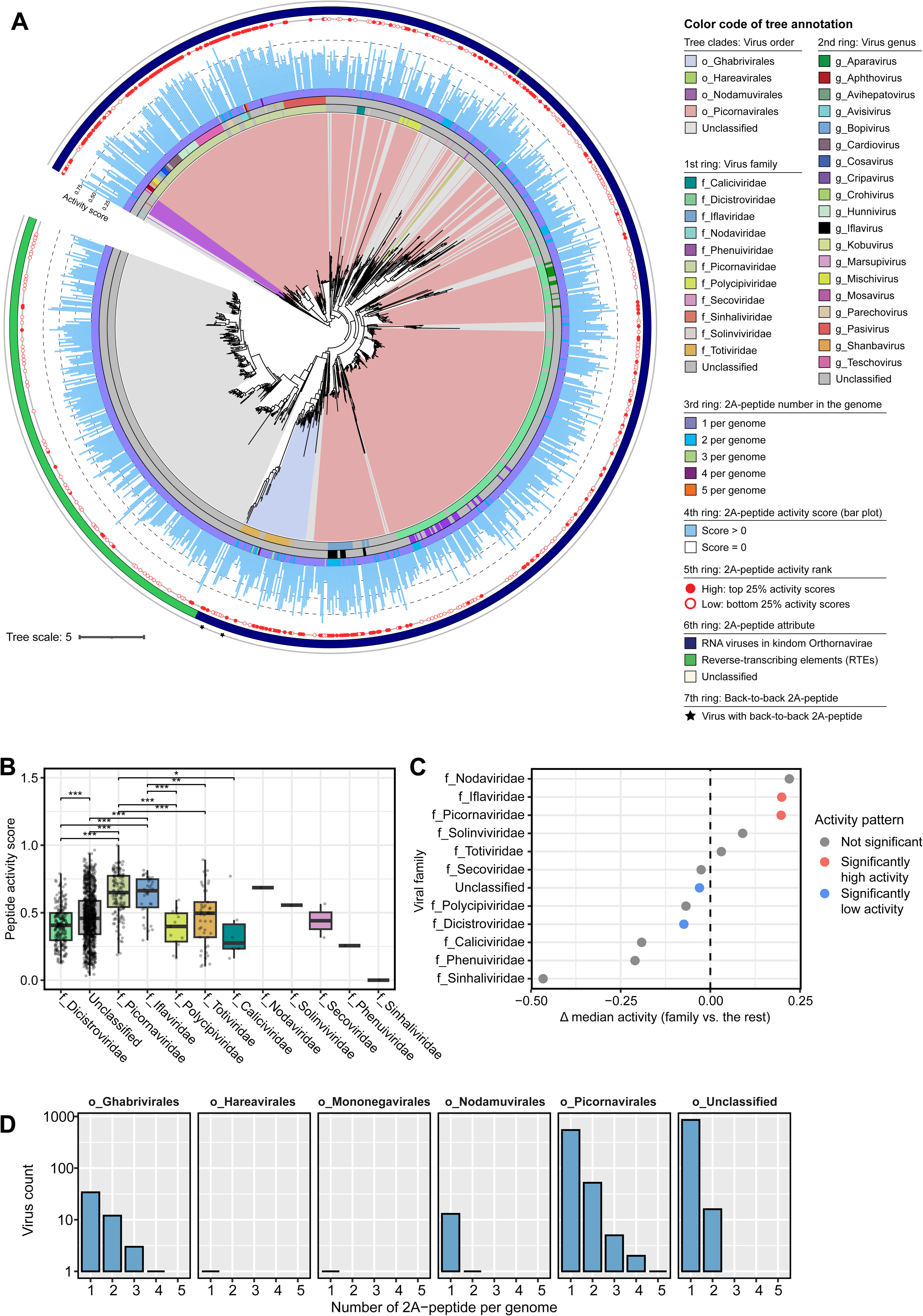
Widespread of predicted and functional Class 2A-peptides in RNA viruses. (**A**) Phylogenetic relationships of viruses that contain the Class A 2A-peptides, identified from the Serratus dataset. The unrooted maximum-likelihood phylogenetic tree was constructed based on palm motif sequences of RNA-dependent RNA polymerases (RdRp). Tree leaves are annotated by viral family (1^st^ ring), genus (2^nd^ ring), number of 2A-peptides detected per genome (3^rd^ ring), normalized 2A-peptide activity score on average for the genome (4^th^ ring), activity-score rank (5^th^ ring), putative 2A-peptide source (6^th^ ring), and presence of back-to-back 2A-peptides (7^th^ ring). Tree clades are colored according to viral order from which the 2A-peptides were derived. The scale bar indicates evolutionary distance. (**B**) Average normalized activity scores of 2A-peptides across viral families. Boxplots show the median (center line), interquartile range (box), and values within 1.5 times the interquartile range (whiskers); points beyond the whiskers represent outliers. Pairwise comparisons between families were performed using pairwise two-sided Wilcoxon rank-sum tests; significant differences are indicated by asterisks (\**P* < 0.05, \*\**P* < 0.01, \*\*\**P* < 0.001). (**C**) Identification of viral families with unusually high or low 2A-peptide activity. For each viral family, the median normalized activity score was compared with that of all remaining families combined using one-versus-rest Wilcoxon rank-sum tests with Benjamini–Hochberg correction. Points represent the difference in median activity score between the indicated family and all other families (Δ median activity). Positive values indicate higher activity, whereas negative values indicate lower activity relative to the remaining viral families. Significant families are highlighted according to directionality of the effect. (**D**) Histogram showing the distribution of viral contigs encoding multiple class A 2A-peptides. The x-axis indicates the number of the 2A-peptides encoded per contig, and the y-axis shows the corresponding number of contigs.

Phylogenetic reconstruction based on RdRp-palm sequences revealed that class A 2A-peptide-containing viruses are distributed throughout multiple deeply divergent RNA virus lineages, including positive-sense ssRNA viruses within the *Picornavirales* and *Nodamuvirales*, negative-sense ssRNA viruses within the *Hareavirales*, and double-stranded RNA viruses within the *Ghabrivirales* (Figure 6A). These observations are consistent with the broad taxonomic distribution recently reported by Rao et al. (2025) (Rao *et al*., 2025), while substantially expanding the number of candidates with 2A-peptide-containing genomes recovered from environmental sequence space (Supplementary Figure S5B). Notably, putative class A 2A peptides were also detected in reverse-transcribing elements, extending their distribution beyond canonical RNA viruses in the kingdom *Orthornavirae* (Figure 6A).

Beyond their broad phylogenetic distribution, the putative Class A 2A peptides exhibited several previously unrecognized genomic architectures. Among the 1,547 2A peptide-containing viruses identified, 39 (2.52%) encode multiple class A 2A-peptides within a single genome (Figures 6A, 6D, Supplemental Figures S6). These viruses were distributed across several evolutionary lineages, including the orders *Picornavirales*, *Ghabrivirales*, and *Nodamuvirales*. The most extreme example was an *Avihepatovirus* genome (family *Picornaviridae*), which encodes five distinct 2A peptides (all Class A), representing, to our knowledge, the largest number observed in a single viral genome (Supplemental Figures S7). The genomes with multiple 2A-peptides adopted two principal organizational architectures (Supplemental Figure S7). In the first, two or more 2A-peptides occurred consecutively upstream of the same protein-coding region, forming tandem ribosome-skipping modules as observed in representatives of *Picornaviridae*, *Dicistroviridae*, *Totiviridae*, and viruses within *Nodamuvirales*. In the second, multiple 2A-peptides were positioned at distinct locations within a viral polyprotein, partitioning different functional regions into separate protein products. This arrangement was particularly evident in *Iflaviridae*-like viruses, in which multiple 2A peptides occurred at different positions within the capsid protein region (Supplemental Figure S7). Finally, we identified tandem back-to-back 2A-peptide arrangements in five viral genomes, including two dsRNA viruses belonging to the family *Totiviridae* (order *Ghabrivirales*) and three previously unclassified RNA viruses (Figures 6A, Supplemental Figures S6-S8).

To determine which predicted viral 2A peptides are functional, we cloned the Class A 2A peptide library (1602 predicted unique 2A peptides) within the FRET-based mClover-mRuby reporter. Following lentiviral transduction, FRET-FACS analysis showed 18.64% and 81.36% in the no-FRET (active 2A) and FRET (inactive 2A) cell populations, respectively (Figure 7B) and after sorting, genome purification and amplicon sequencing, we recovered a total of 1502 2A peptide sequences. Ranking the predicted Class A 2A peptides based on the FRET activity score (as described in Figure 1) yielded a distribution spanning active to inactive variants (Figure 7C, Supplemental Data 3). The three 2A peptides (P2A, F2A, C2A) as internal benchmarks displayed the expected high to low 2A activity scores and by immunoblotting (Figures 7C, 7D). We validated representative peptides from high (∼0.9 FRET), mid (∼0.5 FRET), and low (∼0 FRET) FRET-score bins by immunoblotting, showing that high-, mid and low-bin peptides yielded 75–99%, ∼18% to ∼90%, and <60% activities (several near baseline) (Figures 7E, 7F). Thus, the FRET-based reporter reported a range of functional viral Class A 2A peptides.

**Figure 7.**
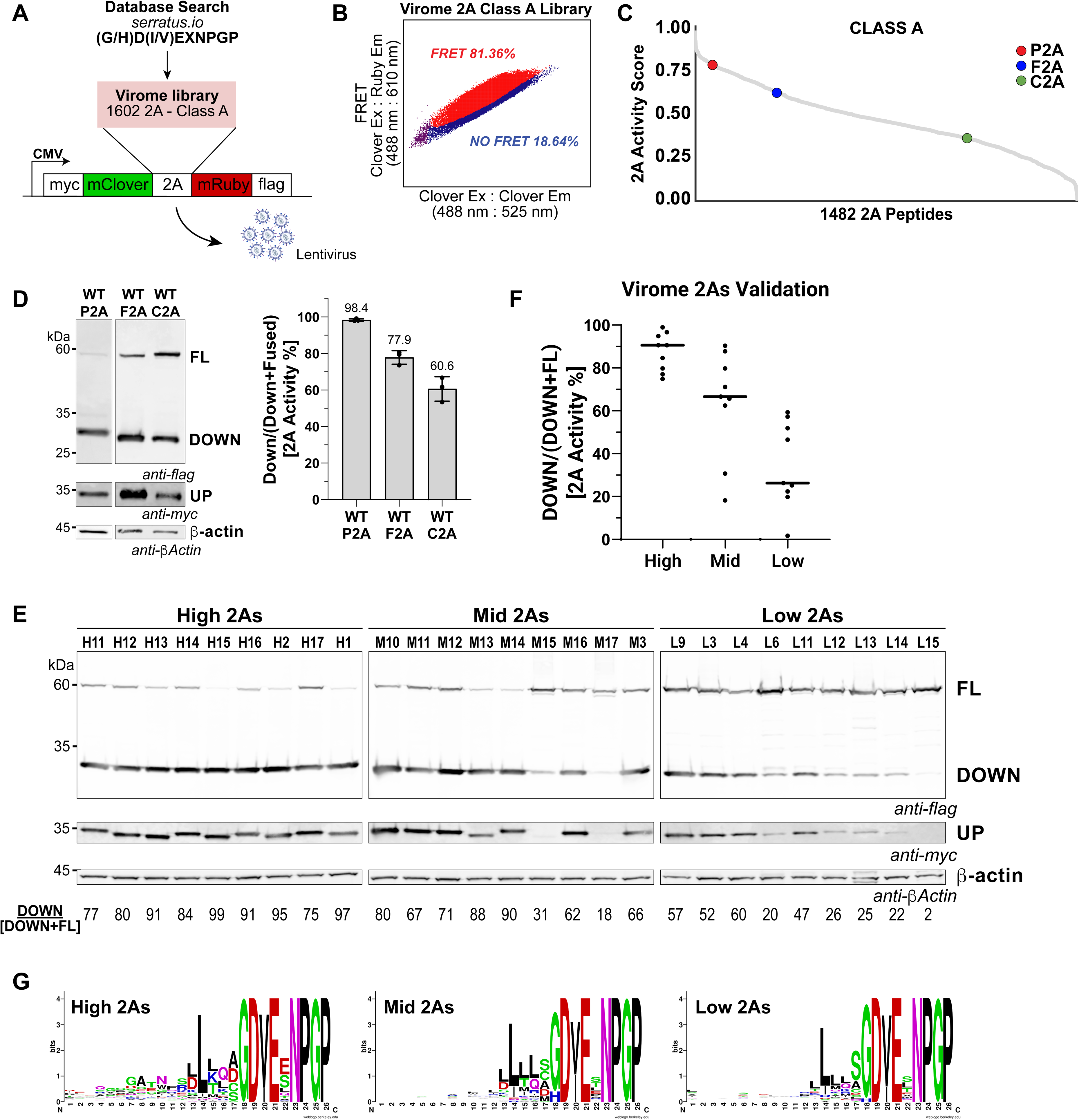
FRET-FACS screening of functional viral 2A peptides. (A) Schematic of the virome 2A library. Candidate 2A peptides were mined from the Serratus.io database using the canonical (G/H)D(I/V)ExNPGP motif. F<u>rom 1,547 genomes</u> that contained a predicted 2A peptide, <u>we identified 1,602 unique putative 2A-encoding nucleotide sequences, corresponding to 1,488 distinct peptides at the amino acid level.</u> The peptides were cloned in-frame between mClover and mRuby in the CMV-driven myc-mClover-2A-mRuby-Flag reporter and packaged into lentivirus for transduction into HEK293T cells. (B) FRET-FACS distribution of the pooled Class A virome library, showing 18.64% of cells in the no-FRET (active) gate and 81.36% in the FRET (inactive) gate. (C) Ranked 2A activity scores for all 1,482 virome peptides that were recovered by NGS, plotted from high to low activity. P2A (red), F2A (blue), and C2A (green) are included as internal benchmarks and highlighted at their respective rank positions. (D) Immunoblots of transfected reporter constructs containing P2A, F2A, and C2A in HEK293T cells (24 hours). 2A activity was calculated as relative 2A activity % = Down/(Down + FL) (right). Error bars represent average ± s.d.; values above each bar indicate mean percent activity (P2A 98.4%, F2A 77.9%, C2A 60.6%). (E) Immunoblots of transfected HEK293T cells with reporter constructs containing virome 2A peptides selected from the high (left, 9 peptides), mid (center, 9 peptides), and low (right, 9 peptides) FRET activity regions. (F) Per-bin distribution of validated 2A activities for the peptides shown in (E). Each dot represents one validated peptide, and horizontal bars indicate the median per bin. (G) Ice-logos of the top 50 high, mid 50, and low 50 activity peptides of the virome library. Letter height is proportional to information content (bits) at each position, and letter color indicates amino-acid chemistry.

Given the broad evolutionary distribution of class A 2A peptides, we next addressed whether their ribosome-skipping activities are associated with their viral lineages. Towards this, we mapped the 2A FRET activity scores onto the phylogeny of 2A peptide-containing viruses and examined activity distributions across taxonomic groups (Figure 6A). Substantial variation in 2A peptide activity was observed across viral families (PERMANOVA: *r*^2^ = 0.330, *p* = 0.001). Such significant difference (post-hoc pairwise PERMANOVA and two-sided Wilcoxon rank-sum tests, *p* < 0.05) not only occurred between viral families from same order (e.g., *Picornavirales*), but also between viral families from different orders (e.g., *Totiviridae* in the order *Ghabrivirales vs. Picornaviridae* or *Iflaviridae* in the order *Picornavirales*) (Figure 6B). Comparing the 2A peptide StopGo activity scores across taxonomic ranks showed that *Picornaviridae* and *Iflaviridae* in the order *Picornavirales* displayed significantly high 2A-peptide activity, while *Dicistroviridae* in the order *Picornavirales* and those unclassified contigs (including RETs and other host-related sequences) exhibited significantly low 2A-peptide activity (Figure 6C). These results suggest that translational StopGo recoding efficiency itself may represent an evolvable trait subject to lineage-specific selective pressures.

To identify potential candidate determinants of StopGo efficiency, we performed motif enrichment analyses comparing high- and low-StopGo activity peptides across multiple taxonomic levels, including individual peptides, contigs, genera, families, orders, classes, and phyla. Analyses using both the top-versus-bottom 50% and the more stringent top-versus-bottom 25% activity partitions revealed five motifs that were significantly enriched among highly active peptides, from which, the motif GATNFSLL located within the N-terminal of the 2A peptide identified as the most consistently associated with elevated activity across all taxonomic levels (Supplemental Figure S9). GATNFSLL-containing 2A peptides were enriched within the genera *Teschovirus* and *Hunnivirus* of the family *Picornaviridae* (Supplemental Figure S10A). Notably, these same lineages exhibited the highest activity distributions observed across the entire dataset (Supplemental Figure S10D), linking motif occurrence with lineage-specific activity phenotypes. The consistency across active viral 2A peptides indicates that GATNFSLL may represent a key determinant of highly efficient translational StopGo activity.

### Identification of novel functional 2A-like 2A peptides

Recent reports described 2A-like peptide variants, a few of which were tested and found to be functional (Sharma *et al*., 2012; Rao *et al*., 2025; Li *et al*., 2026). Inspired by these findings, we mined the Serratus.io database for 2A-like peptides. For one set, we searched for guided Class B peptides (Rao *et al*., 2025), which are distinguished by an invariant N-terminal tryptophan and a divergent central motif in place of the canonical GDVE stretch as observed in Class A 2A peptides (Rao *et al*., 2025), specifically the motif WXXX(L/V)XXEG(I/V)EX(N/H)PGP. In the second set, we searched for 2A sequences that fall outside the canonical core by searching for motifs that permit a substitution at a single conserved position of the (G/H)D(I/V)ExNPGP core, which we termed Class X 2A-like peptides (Figure 8A). These analyses identified predicted 260 class B and 1,999 class X 2A-peptides distributed across a diverse collection of viral and virus-associated genomes (Figures 8A, 8B), spanning nine RNA virus orders including *Bunyavirales*, *Durnavirales*, *Hepelivirales*, *Ghabrivirales*, *Hareavirales*, *Mononegavirales*, *Nodamuvirales*, *Ortervirales* and *Picornavirales* (Supplemental Figure S11A). Comparison with the recent search of 2A-like peptides conducted in IMG/VR and Swissport databases (Rao *et al*., 2025), our study in the Serratus database were in broad agreement regarding the taxonomic distribution of Class A and B 2A peptides across major RNA virus lineages (Supplemental Figure S5B). However, our study also uncovered 2875 previously undescribed 2A-like peptide sequences. Among all predicted 2A peptide-containing genomes, 2.3% (83/3654) harbour putative 2A peptides from multiple classes (e.g., class A, class B, and class X); only a single viral genome (belonging to unclassified virus) encoded representatives of all three classes (Supplemental Figures S11B-S11D). Of the predicted Class X 2A peptides (1999 peptides), a fraction (323; 16%) could be clustered into at least five distinct sub-classes (X1-X5) that spanned several viral families including in endogenous retrovirus-related sequences (Figure 8A, Supplemental Figure S13B). All clustered Class X 2A-like motifs contained the C-terminal NPGP and a V or I at position 20 and distinct adjacent residues at positions 18 and 19.

**Figure 8.**
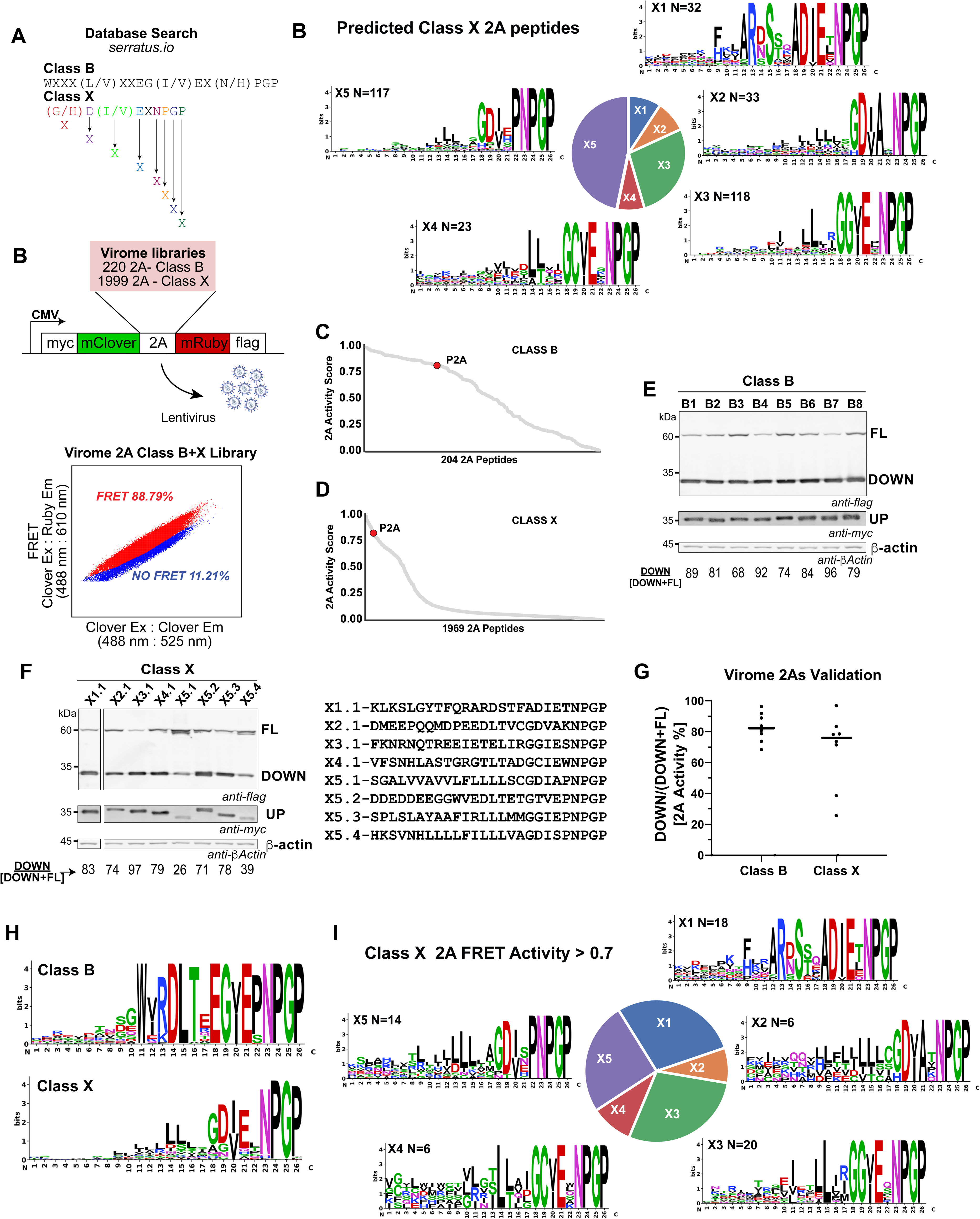
Screening for functional virome-derived Class B and Class X 2A-like peptides. (A) Candidate 2A peptides were mined from the Serratus.io database using a Class B motif (WXXX(L/V)XXEG(I/V)EX(N/H) PGP) and eight Class X motifs, which allow a substitution at one conserved position of the canonical (G/H)D(I/V)ExNPGP core; the resulting Class X 2A peptides were clustered into five different sub-classes, ice logos of each sub-class and their distribution are shown (right). (B) The resulting libraries (220 Class B and 1,999 Class X peptides) were cloned in-frame between mClover and mRuby in the CMV-driven myc-mClover-2A-mRuby-Flag reporter and packaged into lentivirus. FRET-FACS scatter plot of the pooled Class B + X virome library (FRET Clover Ex: Ruby Em 488:610 nm versus mClover Ex: Em 488:525 nm), showing 11.21% of cells in the no-FRET (active) gate and 88.79% in the FRET (inactive) gate. (C) Ranked 2A activity scores for the 204 Class B peptides, plotted from high to low activity, with P2A (red) as an internal benchmark. (D) Ranked 2A activity scores for the 1,969 Class X peptides, plotted from high to low activity, with P2A (red) as an internal benchmark. (E) Representative Immunoblots of transfected reporter constructs containing indicated Class B or X 2A peptides in HEK293T cells (24 hours). (F) Sequences of the eight Class X 2A-like peptides (X1-X5.4) that were validated in (E). Relative 2A activity % = Down/(Down + FL) is indicated below. (G) Scatter plot of the 2A peptide activity scores quantified from immunoblot analysis; horizontal bars indicate the median 2A activity per class. (H) Icelogos of the 2A peptides with a FRET score >0.7 for Class B (top left) and Class X 2A-like peptides (bottom left). (I) Pie chart showing the distribution of the active Class X 2A-like peptides (FRET activity score >0.7). Icelogos of the five clusters of Class X 2A-like peptides are shown.

To identify functional 2A-like peptides, we cloned the predicted Class B and X 2A-like sequences (combined library) in-frame into the 2A-sensing FRET-based reporter, packaged the libraries into lentivirus, transduced HEK293T cells, and sorted the pooled populations by FRET-FACS (Figure 8A). From FACS analysis, 11.2% and 88.8% of cells were in the no-FRET (active) and the FRET (inactive) populations, respectively (Figure 8B). After FRET-FACS sorting and amplicon sequencing, 204 Class B and 1969 Class X 2A sequences were recovered across all replicates (2A peptides that were absent from at least one replicate were excluded). The Class B 2A peptides displayed a continuous gradient from fully active to inactive, with the benchmark P2A ranking high in activity (Figure 8C). In contrast, the Class X 2A-like peptides distribution dropped steeply, with only a small subset of high-activity peptides (n=139, 2A FRET activity score >0.7) (Figure 8D, Supplementary Figure S5C). Mapping highly active Class B and X 2A peptides (>0.7) onto their viral genomes revealed that the 2A-like peptides are distributed throughout the polyprotein (Supplemental Figure S13A).

The FRET-FACS screen recovered many highly active Class B peptides (n=81, >0.7). Selected Class B 2A peptides with high 2A activity scores (>0.7) were individually validated by immunoblotting analysis (B1–B8, Supplemental Data 3) showing a range of 68-96% StopGo activity (Figure 8E), thus demonstrating that highly functional Class B 2A peptides are more abundant than previously appreciated (Rao *et al*., 2025). To identify the residues underlying activity, we generated Icelogos of Class B 2A peptides with a FRET activity score > 0.7, revealing an enriched functional motif W(Y/I) RDLTEEG(V/I)EPNPGP (Figure 8H) for high 2A StopGo activity.

Similar to Class B 2A peptides, we identified multiple functional Class X 2A-like peptides. By immunoblotting in HEK293T cells, we validated select Class X1-5 2A-like peptides (Figure 8F), showing 26%-97% 2A Stop-Go activity. These results revealed a number of features. The 2A-like peptides containing deviations within the core (G/H)DxExNGP still retained relatively high Stop-Go activity. For example, Class X3.1 and X5.2 2A-like peptides containing a G19 and T19, respectively, still displayed high StopGo activities (97% and 71%, respectively) (Figures 8F, 8G). Similarly, X2.1 and X5.1, both having A21 and X5.4 containing a S21, all showed active StopGo activities (74% and 39% activity, respectively). Alignment of highly active Class X 2A peptides for each subclass (>0.7 2A FRET activities) revealed distinct motifs within the core C-terminal domain but also within the central domain from residues 12-18 (Supplemental Figure S13B). Together, these findings reveal unexpected plasticity in the 2A core: non-canonical residues can substitute at otherwise conserved positions without loss of activity and in fact, can retain high StopGo activity. The Class X 2A-like peptides thus represent novel classes of highly functional 2A sequences.

### Functional characterization of a viral tandem 2A peptide

From the viral 2A activity screen, the H15 2A peptide displayed one of the highest StopGo activities (Figure 7E). Upon closer inspection, the H15 2A peptide that was tested (26 amino acid in length) contains two DVESNPGP motifs, one at the N- and C-terminus of the 2A peptide. This raised the possibility that the natural locus encodes a back-to-back tandem 2A peptide architecture, which we have named H15A and H15B (Figure 9A). Notably, both H15A and H15B carry a W(V/I) RDL sequence near their N-terminus and a GDVESNPGP motif at their C-terminus. Interestingly, W(V/I) RDL is characteristic of Class B 2A peptides, whereas GDVESNPGP is characteristic of Class A 2A peptides, suggesting that the tandem 2A peptides may represent a novel subclass containing a hybrid Class A/B chimera. Searching the virome for this arrangement, we identified multiple 2A peptides sharing the same chimeric Class A/B 2A-like motif (n=54) (Figure 9B, Supplemental Data 3). The source viral genome encoding H15 contains three 2A peptides (orange): one located further upstream and the two tandem motifs, H15A and H15B, that together make up H15 (Figure 9A). All three lie within ORF1, which encodes the capsid/coat protein (blue), while ORF2 encodes the RNA-dependent RNA polymerase (RdRp, red) together with its RdRp-palm subdomain (dark red).

**Figure 9.**
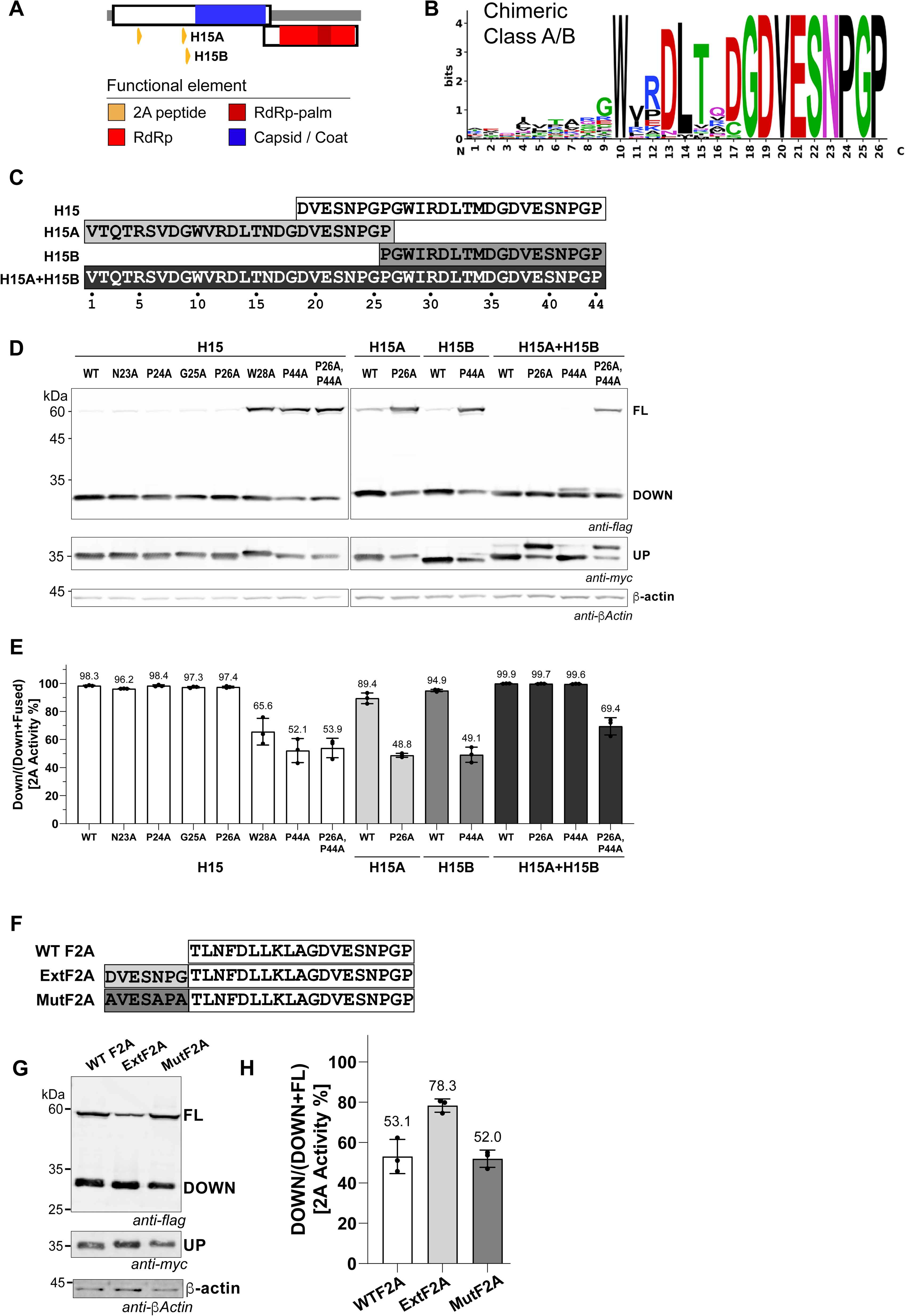
Characterization of a natural functional viral tandem 2A peptide. (A) Genomic location of H15 within its source viral genome, showing the two tandem 2A peptides H15A and H15B. Functional elements are colored as indicated: 2A peptide (orange), RdRp (red), RdRp-palm (dark red), and capsid/coat (blue). (B) Icelogo of the chimeric Class A/B 2A-like peptides across the 26-residue peptide (positions 1–26); letter height is proportional to information content (bits) at each position. (C) Sequence architecture of H15 and derivative constructs. H15, the original virome hit (positions 19–44 of the full-length region), contains a single C-terminal -DVESNPGP motif. Inspection of the upstream genomic context revealed an additional -DVESNPGP-like motif located further upstream, indicating that the natural locus encodes a back-to-back (tandem) 2A architecture. Three additional constructs spanning the full 44-residue region were generated: H15A (positions 1–26), comprising the upstream motif; H15B (positions 26–44), comprising the downstream motif; and H15A+H15B (positions 1–44), the full back-to-back tandem reconstitution. Boxed regions highlight each ribosomal skipping motif. (D) Immunoblots of transfected reporter constructs containing either the wild-type or mutant H15, H15A, H15B and H15A+H15B 2A peptides in HEK293T cells (24 hours). (E) Quantification of 2A activity for the constructs in (D) as relative 2A activity % = Down/(Down + FL). Error bars represent average ± s.d.; values above each bar indicate mean percent activity. (F) Immunoblots of HEK293T cells transfected with reporter constructs containing the indicated F2A wild-type 2A peptides (19 amino acids in length) or ExtF2A 2A pepdies with the H15 upstream motif (DVESNPG) (26 amino acids in length) or a MutF2A (AVESAPA) added to their N-terminus. (G) Quantification of (F) as relative 2A activity % = Down/(Down + FL). Error bars represent average ± s.d.; values above each bar indicate mean percent activity.

To test the functionality of the H15A/B 2A peptides, we generated three mCLover-2A-mRuby reporter constructs and tested in HEK293T cells followed by immunoblotting: (i) H15A, comprising the upstream 2A peptide (positions 1–26); H15B, comprising the downstream 2A peptide (positions 26–44); and both H15A+H15B, the tandem 2As (positions 1–44). We also introduced alanine substitutions at DXEXNPGP within each motif to distinguish contributions of each 2A peptide (Figures 9C, D). The wild-type H15 yielded high cleavage activity (98.3%), comparable to P2A. Single substitutions, N23A, P24A, G25A, P26A within H15 were well tolerated and still retained high 2A StopGo activity (96.2–98.4%), indicating that the upstream DXEXNPGP motif present (positions 19– 44) contributes to little measurable StopGo activity. In contrast, substitutions at P44 to alanine of the C-terminal DXEXNPGP motif reduced H15 2A activity (52.1% activity) and the W28A substitution reduced activity to 65.6%. The double mutant P26A+P44A, which simultaneously disrupts the upstream and the C-terminal prolines of DXEXNPGP, resulted in 53.9% activity, comparable to the single P44A mutant, further confirming that the upstream motif within H15 is non-functional and that activity in this construct is driven by the C-terminal DXEXNPGP motif.

Analysis of the individual H15A and H15B 2A peptides clarified the contribution of each motif. Immunoblotting analysis of the reporter containing H15A alone led to 89.4% StopGo activity, and P26A substitution reduced activity to 48.8%; H15B alone produced 94.9% activity, and P44A reduced it to 49.1% (Figures 9D, 9E). Thus, each motif is independently functional, and each is sensitive to disruption of its critical proline. Combining both motifs in tandem (H15A+H15B) as in its natural arrangement resulted in high StopGo activity (99.9%), and either P26A or P44A mutations in this context was similarly active (∼99%). Only simultaneous disruption of both motifs (P26A+P44A) reduced activity to 69.4%. Direct evidence that both motifs are independently active in the back-to-back context was obtained from the upstream (UP) protein blot which can distinguish usage of the H15A or H15B 2A peptide activity.

The high StopGo activity of H15 raised the question of whether the upstream DXEXNPGP motif can increase the activity of another 2A peptide. To address this, we inserted the DVESNPG or a mutated version (AVESAPA) upstream of F2A and measured 2A activity by immunoblotting (Figures 9E, 9F). While F2A alone displayed 53.1% StopGo activity, inserting DXEXNPGP upstream of F2A increased StopGo activity to 78.3% whereas inserting the mutant version did not affect StopGo activity (52.0%). These results demonstrated a novel arrangement whereby an upstream DXEXNPGP stimulates StopGo activity of a downstream functional DXEXNPGP motif.

### 2A peptide activities in diverse mammalian cells

To test whether the high-activity 2A peptides retain their activity across distinct cell types, we transfected the mClover-2A-mRuby reporters containing P2A, F2A, H1, or H15 across five mammalian cell lines: HEK293T, Jurkat, MDA-MB-231, primary keratinocytes, and normal human bronchial epithelial (NHBE) cells (Supplemental Figures S14A, S14B). P2A was highly active in all five cell lines (∼98-99%) and F2A StopGo activity was lower and more variable across cell lines (44.4% in KC to 67.3% in MDA-MB-231). The virome-derived 2A peptides H1 and H15 were highly active in all cell types (97-99%). These results demonstrated that the StopGo activities are intrinsic within each 2A peptide sequence.

### Functional 2A peptides in reverse-transcribing elements (RTEs)

A substantial proportion of class A, class B, and class X 2A peptide-containing contigs were associated with reverse-transcribing elements (Figure 10A; total 581 predicted 2A peptides) (Supplemental Figures S11A, S12). RTE-associated contigs accounted for 21.3% (330/1,547) of putative Class A, 6.3% (13/206) of putative Class B, and 11.9% (237/1,985) of putative Class X 2A-peptide-containing contigs (Figure 10A). These sequences encompassed diverse RTEs, including group II intron-like, telomerase-like, and CRISPR-RT-like elements, as well as retroelements comprising non-LTR retrotransposon-like, LTR retrotransposon-like, and retrovirus-related sequences (Figure 10A, Supplementary Figure S12, Supplemental Data 4. Across all three predicted 2A-peptide classes, RTE-associated sequences were dominated by non-LTR retrotransposon-like elements (∼79%), whereas predicted Class X 2A peptides were additionally detected in endogenous retrovirus-related sequences, and Class A 2A peptides were uniquely associated with telomerase-like and CRISPR-RT-like elements (Figure 10A, Supplementary Figure S12).

**Figure 10.**
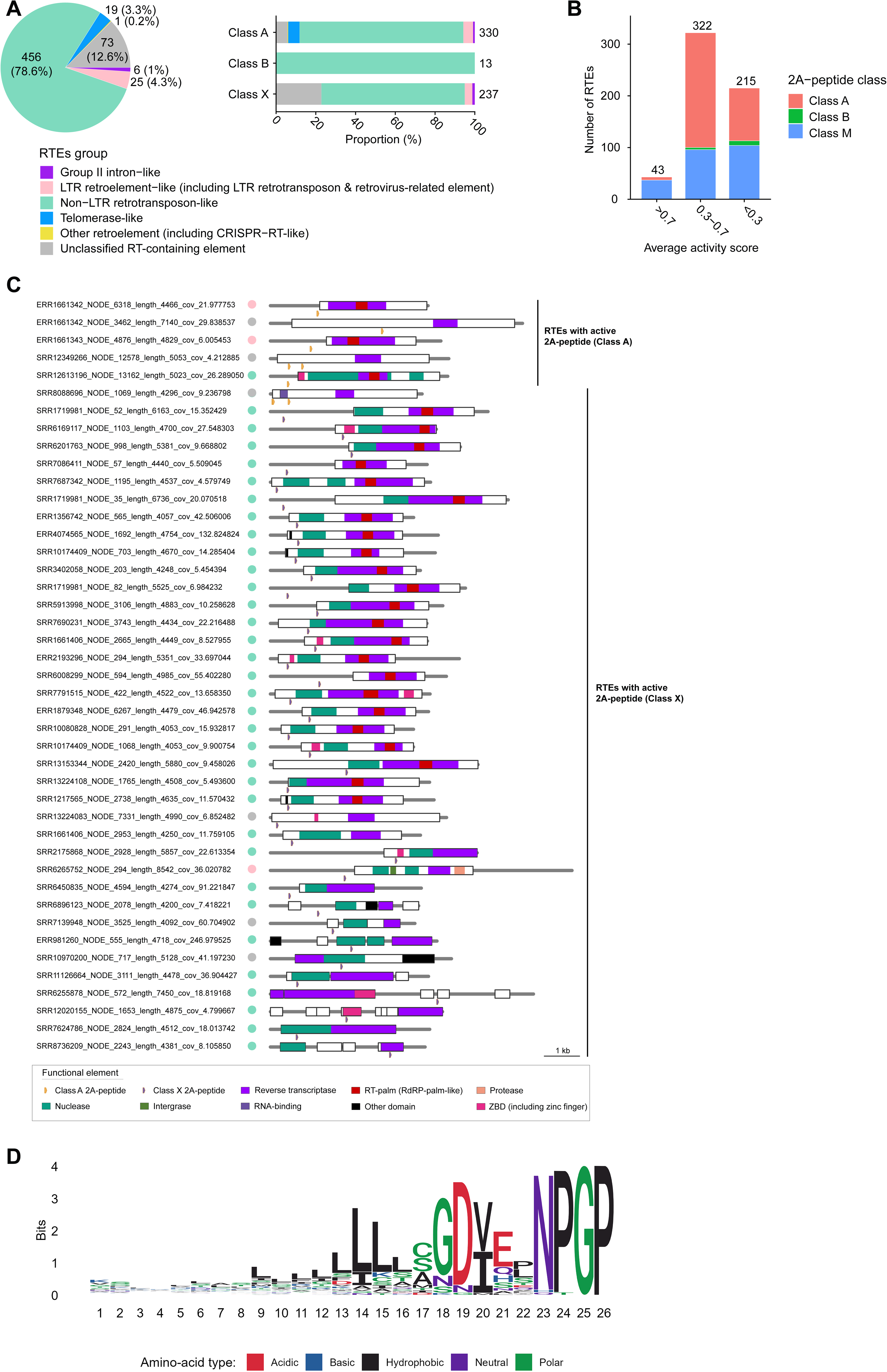
Identification of 2A peptides in reverse-transcribing elements (RTEs) (A) Distribution of different types of the RTEs according to the presence of Class A, Class B, and/or Class X 2A-peptides. (B) Distribution of RTEs across different activity groups based on the activity scores of their 2A peptides. (C) Genomic organization of RTEs containing active 2A-peptides. A 2A-peptide activity score greater than 0.7 was considered indicative of an active 2A-peptide. RTE types are indicated by colored circles, with colors corresponding to the legend in panel A. (C) Putative hosts or habitats of the identified 2A-peptide-containing RTEs. Putative host or habitat assignments were inferred from the sample metadata associated with the original sequences, retrieved from the NCBI SRA, EMBL ENA, and DDBJ DRA databases. The left panel shows the number of 2A-peptide-containing RTE contigs identified across different host or habitat groups, while the right panel shows their proportional distribution among all identified 2A-peptide-containing RTEs.

Of the 580 RTE-associated contigs that contained a putative 2A sequence, only a fraction (7.4%, 43/580) contained 2A peptides that were highly active (2A FRET activity score >0.7; Figure 10B) and most (56%) displayed mid-range FRET activity scores. The highly functional RTE-associated 2A peptides (>0.7) belonged to Classes A and X, but not Class B. Mapping the location of the RTE-associated active Class A and X 2A peptides (>0.7) onto the genome showed a distributed arrangement across the viral open reading frames, suggesting distinct 2A-mediated viral polyprotein processing across RTE-containing lineages (Figure 10C). Together, these findings reveal a broader distribution of functional 2A elements across diverse RTEs than anticipated (Odon *et al*., 2013).

We next examined the biological contexts in which 2A-peptide-containing RTEs were detected. Because metagenomic and transcriptomic assemblies cannot unambiguously establish the cellular host of an RTE, we used the source metadata of the corresponding sequencing datasets as a proxy for their plausible biological and environmental origins. RTE-associated 2A-peptides were recovered from a remarkably broad range of sample types (Supplementary Figure S15). In addition to environmental and complex host-associated metagenomes for which a putative host could not be assigned, these RTEs were detected in datasets derived from diverse metazoans, including amphibians, annelids, arachnids, cnidarians and corals, crustaceans, echinoderms, fishes, humans and other mammals, insects, molluscs, nematodes, and sponges, as well as from protist- and bacterium-associated datasets. Because these sequences could originate either from the sampled organisms themselves or from associated microbiota, parasites, or other organisms present in the samples, their definitive host range cannot be inferred from sequencing metadata alone. Nevertheless, their recovery across phylogenetically and ecologically diverse datasets demonstrates that 2A-peptide-containing RTEs are broadly distributed across biological systems and suggests that StopGo-like translational recoding is considerably more widespread among reverse-transcribing elements than previously appreciated.

## DISCUSSION

Emerging evidence indicates that the ribosomal exit tunnel serves as a key regulatory hub, controlling translation and co-translational folding through interactions with the nascent peptide (Voss *et al*., 2006; Bhushan *et al*., 2010; Wilson and Beckmann, 2011; Ito and Chiba, 2013; Ito, 2014; Dao Duc *et al*., 2019). The 2A peptide exploits this property through one of the most unusual recoding mechanisms whereby the nascent 2A peptide interacts with the exit tunnel to direct peptidyl-tRNA hydrolysis and translation resumption, releasing the upstream protein while allowing synthesis of the downstream protein to continue. Although the conserved C-terminal D(V/I)ExNPG↓P motif of Class A 2A peptides is essential for this activity, how the remaining residues dictate its efficiency has remained unclear. This is increasingly relevant as the diversity of predicted 2A sequences continues to expand (Luke *et al*., 2008; de Lima and Lanza, 2021; Rao *et al*., 2025; Li *et al*., 2026), yet only a small fraction has been tested for function. To address this, we developed a high-throughput 2A-sensing FRET-FACS reporter assay that quantifies StopGo activity across thousands of natural and engineered variants in a single experiment, assigning each a continuous activity score. Using single-residue libraries of two 2A peptides, P2A and F2A, we comprehensively mapped their sequence-to-function landscapes in mammalian cells, linking activity to specific residue properties at the key positions that direct StopGo. Applying the same system to virome-derived libraries, we functionally ranked thousands of natural 2A peptides, mapped their activity onto the viral phylogeny, and uncovered new 2A motifs and classes, revealing that StopGo efficiency varies both across and within viral lineages.

The comprehensive functional landscape provides molecular insight into the 2A peptide– ribosome exit tunnel interactions that promote StopGo, and substantially extends previous efforts that mined 2A peptides bioinformatically and tested them one by one (Rao *et al*., 2025; Li *et al*., 2026). Using P2A and F2A as models, the mutagenesis libraries revealed an asymmetric landscape: the C-terminal core is strictly required, as expected, whereas the N-terminal half is broadly permissive yet contains specific residues that still tune activity. The positions our sensitivity analysis marked as most constrained (residues 4, 9, and 15) correspond to key contacts in the cryo-EM structure of a ribosome stalled on F2A (Li *et al*., 2026): residue 4 can π-stack with rRNA, residue 9 hydrogen-bonds within the nascent helix backbone, and residue 15 hydrogen-bonds to the tunnel wall. The sharp drop in sensitivity across positions 5 to 9 overlaps the short helical segment resolved in the structure, tying skipping to a defined nascent-chain conformation (Fig 4). Beyond the core, functional mining of viral 2A peptides likewise converged on the N-terminal half as a positive determinant of efficiency (Figs 4, 7; Fig S9). Motif-enrichment analysis comparing high- and low-activity peptides across taxonomic levels identified five motifs enriched among active peptides, of which the N-terminal “GATNFSLL” was the most consistent signature of high activity at every level (Fig S9). Furthermore, this signature coincides within the same N-terminal segment that our deletion and chimeric libraries pinpointed as the boundary between high- and low-activity peptides: grafting the P2A N-terminal region bearing “GATNFSLL” onto the low-activity C2A backbone was sufficient to convert C2A into a highly active peptide (Fig 4). Together, single-residue mutagenesis, deletion and chimeric analysis, and virome-scale functional mapping converge on a model in which N-terminal residues that line the exit tunnel help set the nascent-chain conformation required for skipping (Figs 2, 3, 4).

Our study both supports and revises the prevailing view of an invariant core: consistent with the recently expanded (D/G/C/N)(V/I)ExNPGP core, many non-canonical variants (Class X 2A-like peptides) remain active, and functional Class B peptides carrying an N-terminal tryptophan are far more abundant than previously appreciated (Rao *et al*., 2025; Li *et al*., 2026). More unexpected, relative to the prevailing “core-centric” model, is that non-core N-terminal residues and local sequence context are strong determinants of activity. This is exemplified by the shared lysine at position 8, which behaves differently in P2A and F2A which is a clear case of context-dependence (Figures 2, 3). Furthermore, substitution of the terminal proline retained significant StopGo activity *in vitro* but was masked in cells, where N-end rule degradation of the resulting downstream protein reduces its levels. These findings have implications for how viral protein stoichiometry is set during infection and suggests that some active 2A-like peptides may have been overlooked. It will be of interest to define the molecular rules governing class B and class X 2A-like peptides and whether they resemble those established here for class A, and how they engage the exit tunnel to impact recoding.

Mapping 2A peptide activity in an evolutionary frame reveals that StopGo efficiency is an evolvable, lineage-associated trait. Activity varied significantly across viral families (PERMANOVA r² = 0.330, p = 0.001), high in *Picornaviridae* and *Iflaviridae* and low in *Dicistroviridae* and unclassified sequences, indicating that recoding efficiency is itself under lineage-specific selection rather than a fixed property of the motif (Figure 6). The tandem H15 2A element shows how this diversity is exploited: two back-to-back motifs, each a chimeric Class A/B motif, lie within the capsid-encoding region and likely generate a shorter, abundant capsid and a minor, longer form from a single stretch of sequence (Figure 9). Producing multiple protein forms from one region is a common strategy by which RNA viruses expand a limited coding capacity. For example, flaviviruses do so by first making a longer membrane-anchored capsid and then cleaving it to the mature form that packages the genome (Tan *et al*., 2020; Barnard *et al*., 2021), and the dicistrovirus PSIV achieves a similar outcome through stop-codon readthrough (Kamoshita and Tominaga, 2019). The tandem 2A architecture of H15 adds ribosomal skipping to this repertoire, a compact, RNA-structure-independent route to two protein forms from one region. The recovery of functional 2A elements in reverse-transcribing elements extends StopGo still further, beyond canonical RNA viruses. Practically, this work delivers a catalogue of natural 2A peptides exceeding 99% activity that may prove valuable for research and therapeutic applications.

In summary, by coupling a quantitative, high-throughput StopGo reporter to single-residue, deletion, chimeric, and virome-scale libraries, we have moved the study of 2A peptides from one-by-one characterization to a comprehensive, sequence-to-function map. Our results reframe StopGo as a property of the entire nascent 2A peptide, not the conserved core alone, in which N-terminal residues lining the exit tunnel help set the nascent-chain conformation required for skipping. Placed in an evolutionary frame, StopGo efficiency emerges as a tunable, lineage-associated trait that viruses exploit to diversify their proteomes. These findings define the molecular rules of 2A activity, expand the known diversity of functional 2A peptides, and provide a validated toolkit and design principles for multicistronic expression.

## LIMITATIONS

The 2A FRET activity scores were measured in a fixed reporter and cell type although we tested several 2A peptides in several cell types. Further, flanking context and cellular environment both could shape StopGo activity, absolute activities may shift in other constructs or cell types even as relative rankings hold. We searched bioinformatically exact sequences of Class A and B 2A peptides and deviations from the conserved core DxExNPGP of Class A peptides; however, an improved search using Hidden Markov modeling similar to that performed by others (Rao *et al*., 2025) would potentially yield more robust predicted 2A-like peptides.

## ACKNOWLEDGEMENTS

We thank Hema Bommadavara for help in cloning and immunoblotting. We are grateful to Artem Babaian for providing the Serratus viral database. We thank Ivan Sadowski for CL2+ lentivirus training. We thank Lucy Feng, Vincent Halim and Cindy Lam for providing cell lines and/or transfection studies on different cell lines. This study was supported by the UBC Life Sciences Institute Cores (Flow Flow Cytometry (ubcFLOW) and Bioinformatics Core facilities), which is supported by the UBC GREx Biological Resilience Initiative. This study was supported by a CIHR Project Grant (PJT-206125 to EJ), DNA 2 RNA Foundational Projects Program Grant to EJ, NSERC Discovery grants (RGPIN-2020-05348 to KDD and RGPIN-2024-04666 to SS).

