## Supplemental Figures S1-S15 for "Mapping the Functional Landscape and Viral Diversity of 2A Peptides"

**Supplementary Figure S1. FRET-FACS gating strategy and the effect of expression level on FRET.** (A) Stepwise gating strategy for the FRET-FACS reporter across four conditions: mClover alone, mRuby alone, co-transfected mClover + mRuby, and the fused mClover–mRuby protein. Panel 1: pre-gating scatter plots (mRuby Ex: Em 560:610 nm versus mClover Ex: Em 488:525 nm) used to gate double-positive cells. Panel 2: a triangular gate (FRET 488:610 nm versus mRuby 560:610 nm) that excludes false-positive FRET signals arising from excitation of mRuby by the 488 nm laser. Panel 3: a second triangular gate (FRET 488:610 nm versus mClover 488:525 nm) used to determine FRET-positive cells, adjusted to the FRET-negative co-transfected mClover + mRuby cells. (B) Histogram of the double-positive mClover + mRuby population used to gate cells into Low, Medium, and High fluorescence. (C) FRET plots (FRET 488:610 nm versus mClover 488:525 nm) for the Low, Medium, and High populations gated in (B). The percentage of FRET-positive cells is indicated for each population.

**Supplementary Figure S2. Validation of the FRET activity score for position-specific single amino-acid mutagenesis of F2A by immunoblotting.** (A) Heat map of FRET activity scores for amino acid substitutions across the F2A peptide. The x-axis shows the residue position in F2A (1–19), and the y-axis shows the substituted amino acids, sorted from high to low activity score within each column. The color scale indicates the FRET score (0–1), with darker red representing higher activity and lighter red to white representing lower activity. The wild-type F2A sequence is shown at the top for reference. (B) Representative Immunoblots of selected F2A mutants ranked by FRET activity. The full-length fused protein (FL; myc-mClover-F2A-mRuby-Flag) is shown at the top, the separated downstream product (DOWN; mRuby-Flag) is shown in the middle, and the upstream product (UP; myc-mClover) is shown below. Immunoblotting was performed using anti-Flag to detect FL and DOWN products, anti-myc to detect the UP product, and anti-β-actin to detect β-actin as a loading control. Mutants in (B) were selected from the higher FRET activity region. (C) Quantification of F2A mutants' activity from immunoblots band intensity. Band intensities were measured using ImageQuant and used to calculate relative 2A activity % as Down/(Down + FL), where higher values indicate greater 2A efficiency. Error bars represent Mean ± SD.

**Supplementary Figure S3:** Variance decomposition F-values for F2A and P2A and all components analyzed shown in the table above.

**Supplementary Figure S4. Deletion library of the CrPV 2A (C2A) peptide.** 2A FRET activity scores measured by FRET-FACS for N-terminal and internal deletion variants of C2A (IDKCRALLRKRAQLLISGDIESNPGP, 26 amino acids); note the 0–0.3 scale, reflecting the low baseline activity of C2A. Top: N-terminal deletions, in which residues were progressively removed from the N-terminus. Bottom: internal deletions, in which residues between the N-terminal region and the conserved C-terminal -ISGDIESNPGP motif were progressively removed. Each bar shows the 2A FRET activity score for the sequence shown on the left, with wild-type C2A at the top and C2A residues shown in blue.

**Supplementary Figure S5. (A)** Distribution of obtained Class A 2A-peptides-containing contigs identified from the Serratus assembly datasets. (B) Novelty of the 2A peptides detected in this study. The 2A peptides identified in this study were clustered with previously reported 2A peptides

(Rao *et al.*, 2025; Li et al., 2026) to identify identical and novel peptide sequences. Clustering was performed using CD-HIT at 100% amino-acid sequence identity, with the alignment covering 100% of the shorter sequence to accommodate differences in sequence length among studies. See *Methods* for details. (C) Distribution of highly functional ( FRET 2A activity > 0.7) Class A, Class B, and Class X 2A peptide containing contigs identified from the Serratus assembly datasets.

**Supplementary Figure S6. Phylogenetic relationships of 2A-peptide** (class A) identified in the Serratus dataset. The unrooted maximum-likelihood (ML) phylogenetic tree was constructed using amino acid sequences of class A 2A-peptides detected in assemblies from the Serratus dataset. Tree leaves represent individual 2A-peptides and are annotated with features of the corresponding viral genome and peptide: predicted viral family of the host virus ($1^{st}$ ring), predicted viral genus of the host virus ($2^{nd}$ ring), total number of 2A-peptides detected in the same viral genome ($3^{rd}$ ring), normalized activity score of the individual 2A-peptide ($4^{th}$ ring), activity-score rank of the individual 2A-peptide ($5^{th}$ ring), predicted source of the 2A-peptide based on the corresponding viral genome annotation ($6^{th}$ ring), and whether the individual 2A-peptide occurs in a back-to-back arrangement ($7^{th}$ ring). Tree clades are colored according to viral order from which the 2A-peptides were derived. The scale bar represents evolutionary distance.

**Supplementary Figure S7. Representative genomic architectures of viral contigs encoding multiple class A 2A-peptides.** Representative examples illustrating the diverse genomic organizations of multiple class A 2A-peptides identified in this study, including tandem and back-to-back arrangements.

**Supplementary Figure S8.** Phylogenetic relationships of 2A-peptide (class A) identified in dsRNA viruses in the order of *Ghabrivirales*. The unrooted maximum-likelihood (ML) phylogenetic tree was constructed using amino acid sequences of class A 2A-peptides detected in Serratus dataset. Node labels indicate bootstrap support values. Tree leaves represent individual 2A-peptides and are annotated with features of the corresponding viral genome and peptide: predicted viral family of the host virus (1$^{st}$ ring), predicted viral genus of the host virus (2$^{nd}$ ring), total number of 2A-peptides detected in the same viral genome (3$^{rd}$ ring), normalized activity score of the individual 2A-peptide (4$^{th}$ ring), activity-score rank of the individual 2A-peptide (5$^{th}$ ring), predicted source of the 2A-peptide based on the corresponding viral genome annotation (6$^{th}$ ring), and whether the individual 2A-peptide occurs in a back-to-back arrangement (7$^{th}$ ring).

**Supplementary Figure S9. Motif enrichment analysis associated with 2A-peptide activity across viral taxonomic levels.** Dot plots summarize sequence motifs identified by DREME from comparisons between high-activity and low-activity 2A-peptides. Analyses were performed using either the top 50% *vs.* bottom 50%, or the top 25% *vs.* bottom 25% of normalized 2A-peptide activity scores. Motif enrichment was evaluated across multiple taxonomic levels, including individual 2A-peptide, genome, and viral genus, family, order, class, and phylum. Analyses were conducted using either full-length 2A-peptide sequences (26 amino acids) or the conventional partial 2A-peptide region corresponding to the C-terminal 20 amino acids. Colored dots indicate significantly enriched motifs identified in each comparison.

**Supplementary Figure S10. Association of the GATNFSLL motif with elevated 2A-peptide activity in viruses from the family *Picornaviridae*.** (**A**) Distribution of GATNFSLL motif-containing 2A-peptides across viral genera. Bar plots indicate the proportion of 2A-peptides harboring the motif within each genus. (**B**) Genomic organization of viruses harboring GATNFSLL motif-containing 2A-peptides. Schematics show viral genomes from the genera *Hunnivirus*, *Teschovirus*, and *Aphthovirus*, with the positions of the 2A-peptide and RdRp palm domain indicated. (**C**) Sequence logo of class A 2A-peptides containing the GATNFSLL motif in the viruses shown in panel B. (**D**) Identification of viral genera with unusually high or low 2A-peptide activity. For each genus, the median normalized activity score was compared with that of all remaining genera combined using one-versus-rest Wilcoxon rank-sum tests with Benjamini–Hochberg correction. Points represent the difference in median activity score between the indicated genus and all other genera. Positive values indicate higher activity, whereas negative values indicate lower activity. Significant genera are highlighted according to effect direction. (**E**) Normalized activity scores of 2A-peptides across viral genera. Points represent the mean activity score for each viral genus, and error bars indicate the standard error of the mean (SEM).

**Supplementary Figure 11. Identification of class B and class X 2A-peptides in the Serratus dataset.** (**A**) Distribution of class A, class B, and class X 2A-peptides across viral orders. The left panel shows the number of viruses harboring each 2A-peptide class, whereas the right panel indicates their taxonomic composition by viral order. (**B**) Venn diagram showing the numbers of class A, class B, and class X 2A-peptides and their overlap. Shared 2A-peptides detected in two or all three classes are shown in the corresponding intersections. (**C**) Genomic organization of a representative virus harboring all three 2A-peptide classes. The virus belongs to an unclassified lineage within the kingdom *Orthornavirae*. (**D**) Sequence logo of 2A-peptides in the virus shown in panel C.

**Supplementary Figure 12. Genomic organization of different types of 2A-peptides–containing reverse-transcribing elements (RTEs) identified in this study.**

**Supplementary Figure 13. Class X 2A sub-motif conservation, genomic context, and source information.** (A) Ice-logos of the virome-derived Class X 2A peptides, grouped by relaxed motif definition: X1 (N = 30), X2 (N = 16), X3 (N = 49), X4 (N = 6), and the combined X5, X6, X7, and X5 group (N = 152). Each logo spans the 26-residue 2A peptide from N- to C-terminus (positions 1–26); letter height is proportional to information content (bits) at each position, and letter color indicates amino-acid chemistry. (B) Genome organization of the source contigs encoding the eight validated Class X 2A peptides (X1–X5). Functional elements are colored as indicated: 2A peptide (orange), capsid/coat (blue), RdRp (red), RdRp-palm (dark red), helicase (light blue), zinc-binding domain (ZBD, magenta), reverse transcriptase (purple), and other domains (black). Scale bar, 5 kb. (C) Source contig accession identifiers and taxonomic lineages for each of the eight Class X peptides (X1–X8).

**Supplementary Figure 14. High activity 2A peptides maintain robust StopGo activity across diverse mammalian cell lines.** Representative Immunoblots of the P2A, F2A, H1, and H15 reporters expressed in Jurkat, MDA-MB-231, KC, and NHBE cells. The full-length fused protein (FL; myc-mClover-2A-mRuby-Flag) and downstream product (DOWN; mRuby-Flag) were detected with anti-Flag, the upstream product (UP; myc-mClover) with anti-myc, and β-actin with anti-β-actin as a loading control. (B) Quantification of 2A activity for each 2A peptide across five cell lines (HEK293T, Jurkat, MDA-MB-231, KC, and NHBE), calculated as relative

2A activity % = Down/(Down + FL). Error bars represent Mean ± SD; values above each bar indicate mean percent activity.

**Supplementary Figure 15. Putative hosts or habitats of the identified 2A-peptide-containing RTEs.** Putative host or habitat assignments were inferred from the sample metadata associated with the original sequences, retrieved from the NCBI SRA, EMBL ENA, and DDBJ DRA databases. The left panel shows the number of 2A-peptide-containing RTE contigs identified across different host or habitat groups, while the right panel shows their proportional distribution among all identified 2A-peptide-containing RTEs.

Figure S1

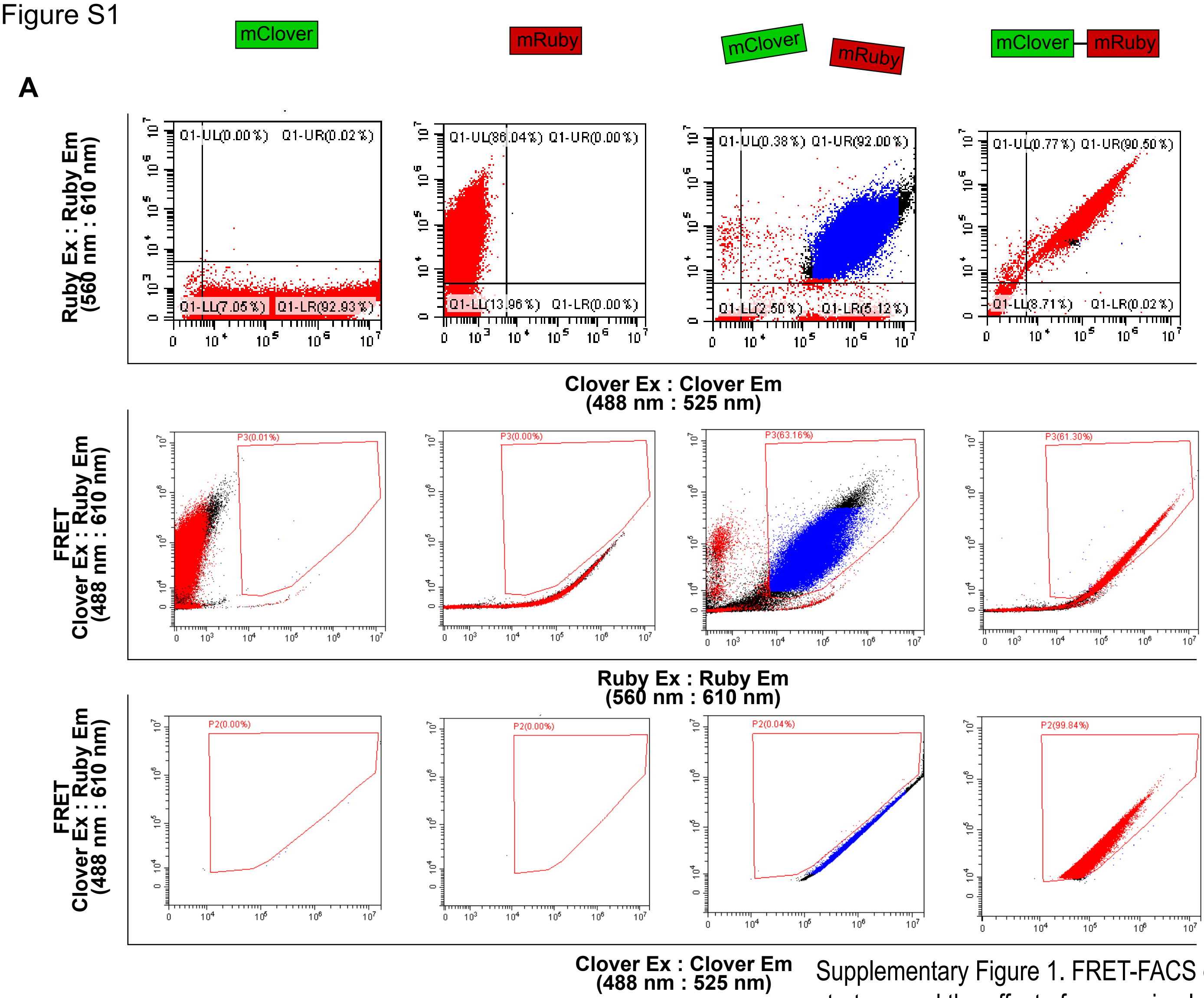


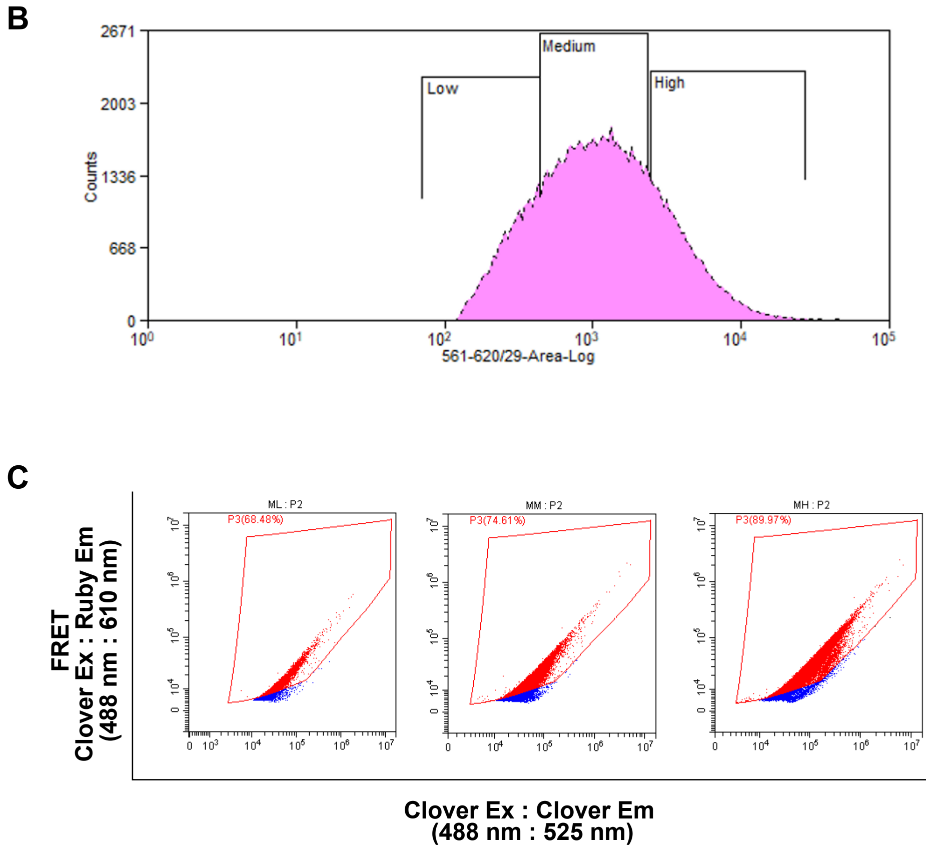


Supplementary Figure 1. FRET-FACS gating strategy and the effect of expression level on FRET. (A) Stepwise gating strategy for the FRET-FACS reporter across four conditions: mClover alone, mRuby alone, co-transfected mClover + mRuby, and the fused mClover–mRuby protein. Panel 1: pre-gating scatter plots (mRuby Ex: Em 560:610 nm versus mClover Ex: Em 488:525 nm) used to gate double-positive cells. Panel 2: a triangular gate (FRET 488:610 nm versus mRuby 560:610 nm) that excludes false-positive FRET signals arising from excitation of mRuby by the 488 nm laser. Panel 3: a second triangular gate (FRET 488:610 nm versus mClover 488:525 nm) used to determine FRET-positive cells, adjusted to the FRET-negative co-transfected mClover + mRuby cells. (B) Histogram of the double-positive mClover + mRuby population used to gate cells into Low, Medium, and High fluorescence. (C) FRET plots (FRET 488:610 nm versus mClover 488:525 nm) for the Low, Medium, and High populations gated in (B). The percentage of FRET-positive cells is indicated for each population.

Figure S2

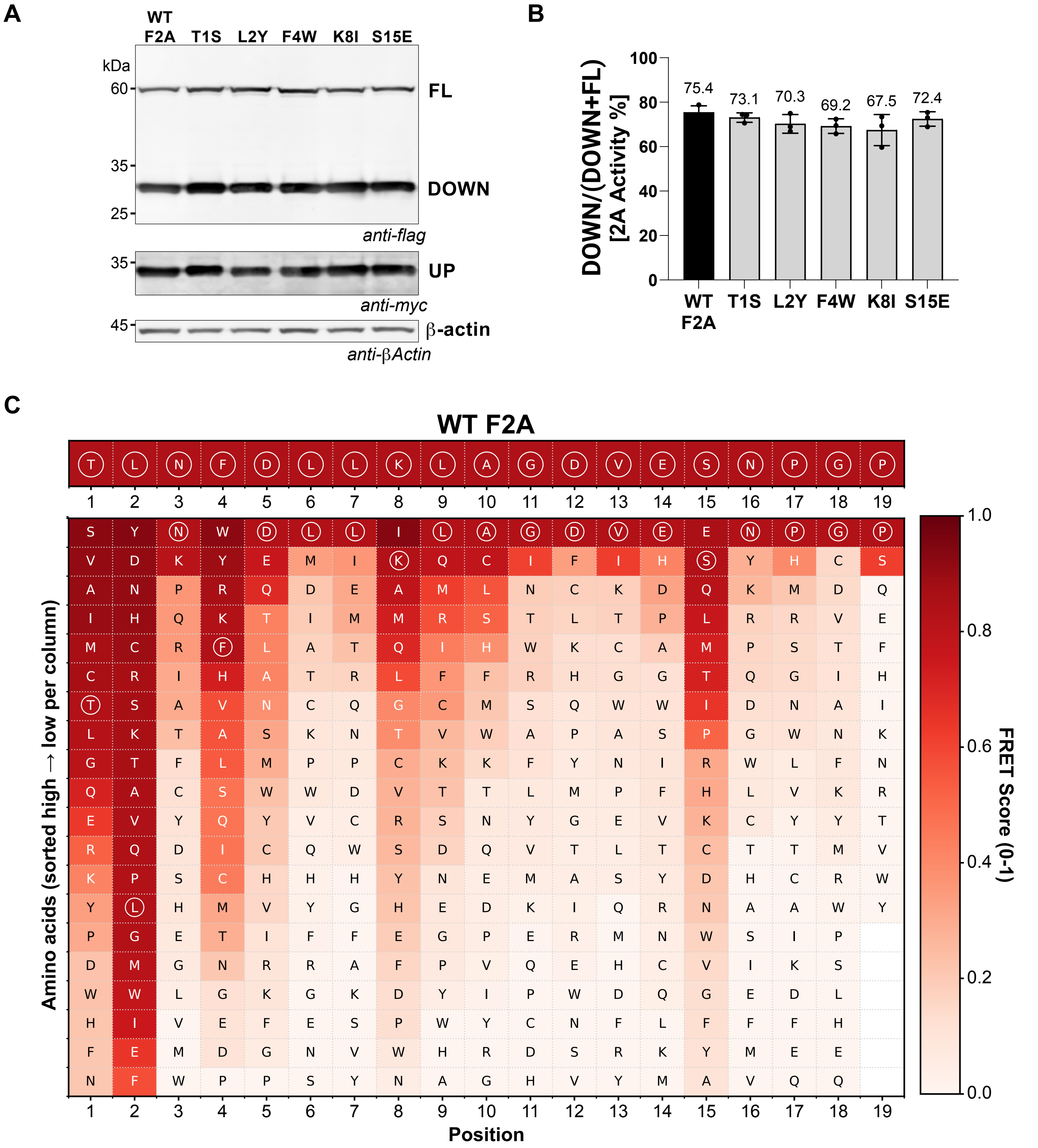


**Supplementary Figure 2. Validation of the FRET activity score for position-specific single amino-acid mutagenesis of F2A by immunoblotting.** (A) Heat map of FRET activity scores for amino acid substitutions across the F2A peptide. The x-axis shows the residue position in F2A (1–19), and the y-axis shows the substituted amino acids, sorted from high to low activity score within each column. The color scale indicates the FRET score (0–1), with darker red representing higher activity and lighter red to white representing lower activity. The wild-type F2A sequence is shown at the top for reference. (B) Representative Immunoblots of selected F2A mutants ranked by FRET activity. The full-length fused protein (FL; myc-mClover-F2A-mRuby-Flag) is shown at the top, the separated downstream product (DOWN; mRuby-Flag) is shown in the middle, and the upstream product (UP; myc-mClover) is shown below. Immunoblotting was performed using anti-Flag to detect FL and DOWN products, anti-myc to detect the UP product, and anti-β-actin to detect β-actin as a loading control. Mutants in (B) were selected from the higher FRET activity region. (C) Quantification of F2A mutants' activity from immunoblots band intensity. Band intensities were measured using ImageQuant and used to calculate relative 2A activity % as Down/(Down + FL), where higher values indicate greater 2A efficiency. Error bars represent Mean ± SD.

| matrix | adj_r2_position | adj_r2_common_slope | adj_r2_varying_slope | f_distance_gain | df1_distance_gain | df2_distance_gain | f_distance p value | f_interact_gain | df1_interact_gain | df2_interact_gain | f_interact_gain p value | common_slope | common_slope_se | species |
|---|---|---|---|---|---|---|---|---|---|---|---|---|---|---|
| Dist_Comp | 0.189733 | 0.1887462 | 0.2009643 | 0.57182129 | 1 | 351 | 0.450043299 | 6.367158302 | 1 | 350 | 0.01206738 | 0.177961496 | 0.235340074 | F2A |
| grantham | 0.189733 | 0.1885353 | 0.1973077 | 0.4804445 | 1 | 351 | 0.488679425 | 4.83601732 | 1 | 350 | 0.02852484 | -0.002056762 | 0.002967306 | F2A |
| Dist_MV | 0.189733 | 0.1890376 | 0.1955267 | 0.69817314 | 1 | 351 | 0.403966636 | 3.831263729 | 1 | 350 | 0.05109943 | -0.003283627 | 0.003929816 | F2A |
| Dist_Polar | 0.200538 | 0.2053548 | 0.2108551 | 3.13365581 | 1 | 351 | 0.077559597 | 3.446457018 | 1 | 350 | 0.06422764 | -0.113181341 | 0.063936548 | P2A |
| Dist_MV | 0.200538 | 0.2228239 | 0.2244055 | 11.09378285 | 1 | 351 | 0.000958239 | 1.715769965 | 1 | 350 | 0.19109754 | -0.012833214 | 0.00385297 | P2A |
| grantham | 0.200538 | 0.2146464 | 0.2145966 | 7.32344471 | 1 | 351 | 0.007138475 | 0.977755955 | 1 | 350 | 0.32343654 | -0.008419548 | 0.003111223 | P2A |
| Dist_Polar | 0.189733 | 0.1964309 | 0.1951641 | 3.9339791 | 1 | 351 | 0.048098728 | 0.447531401 | 1 | 350 | 0.50395059 | -0.119341009 | 0.060169123 | F2A |
| Dist_Comp | 0.200538 | 0.1983102 | 0.1960365 | 0.02182204 | 1 | 351 | 0.882646315 | 0.007323715 | 1 | 350 | 0.93185026 | -0.035967693 | 0.243480893 | P2A |

**Supplementary Figure S3:** Variance decomposition F-values for F2A and P2A and all components analyzed shown in the table above.

Figure S4

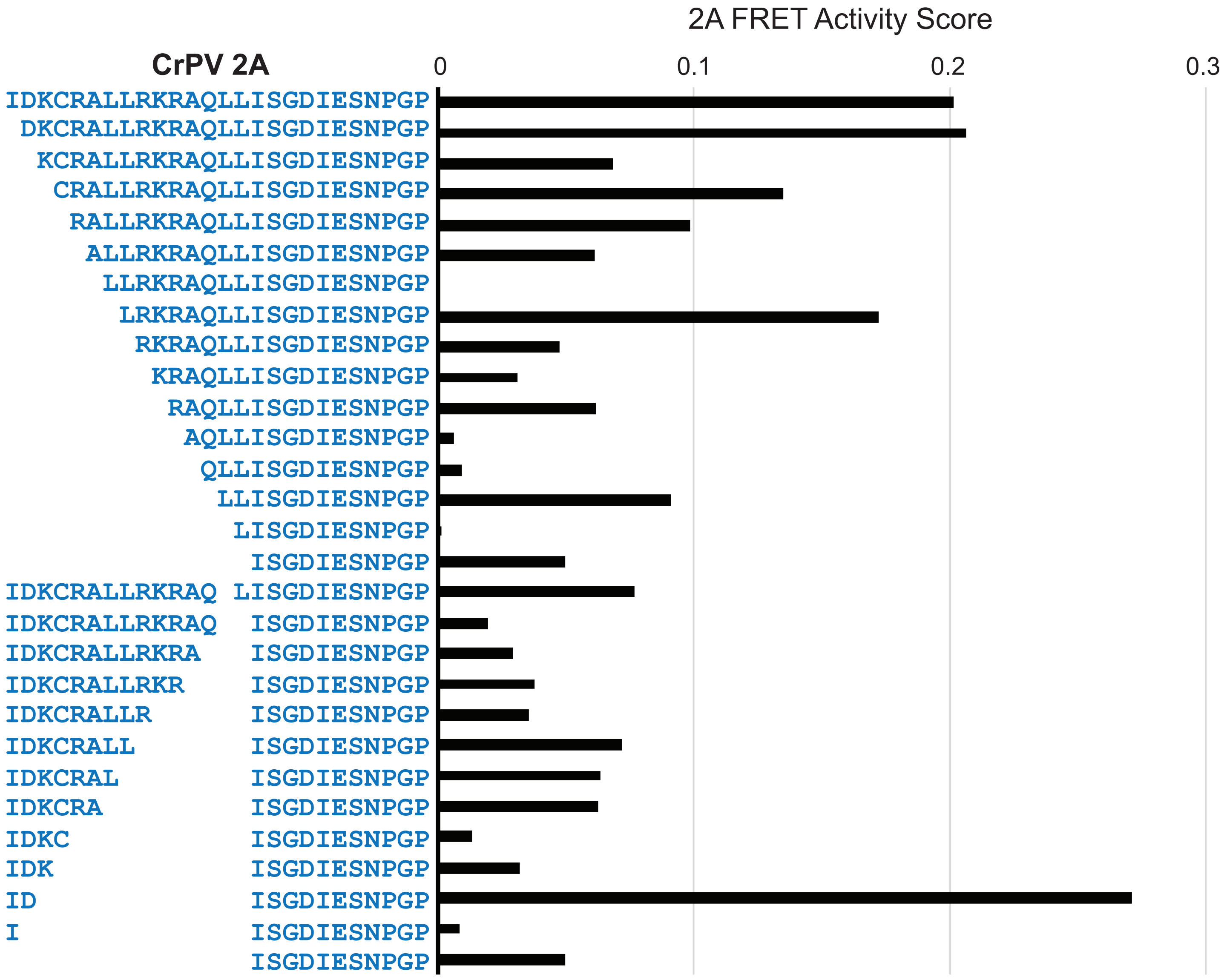


**Supplementary Figure 4. Deletion library of the CrPV 2A (C2A) peptide**. 2A FRET activity scores measured by FRET-FACS for N-terminal and internal deletion variants of C2A (IDKCRALLRKRAQLLISGDIESNPGP, 26 amino acids); note the 0–0.3 scale, reflecting the low baseline activity of C2A. Top: N-terminal deletions, in which residues were progressively removed from the N-terminus. Bottom: internal deletions, in which residues between the N-terminal region and the conserved C-terminal -ISGDIESNPGP motif were progressively removed. Each bar shows the 2A FRET activity score for the sequence shown on the left, with wild-type C2A at the top and C2A residues shown in blue.

Figure S5

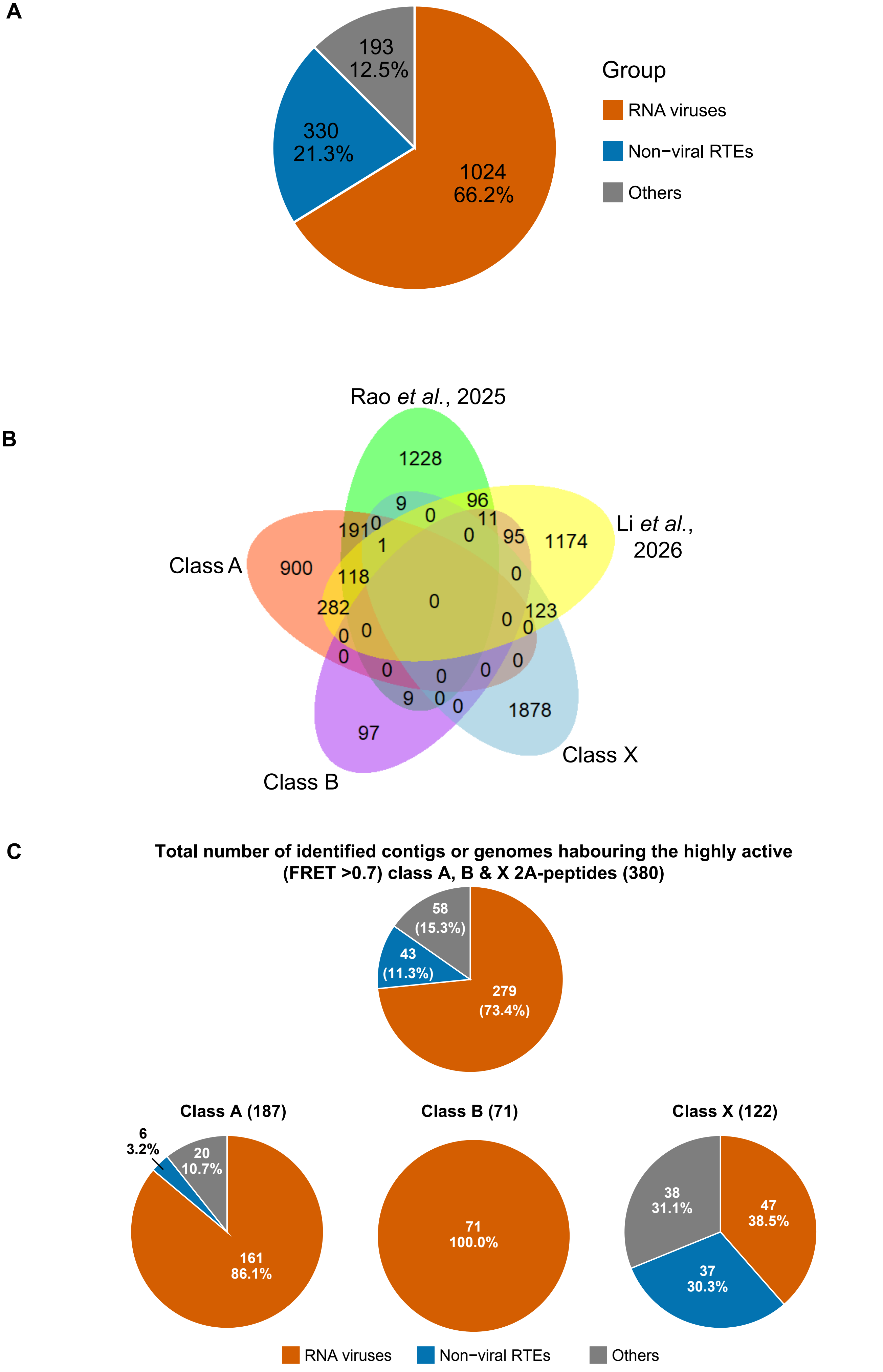
A
193
12.5%
330
21.3%
1024
66.2%
Group
RNA viruses
Non−viral RTEs
Others
B
Rao et al., 2025
Li et al., 2026
Class A
Class B
Class X
1228
1174
900
97
1878
9
96
0
11
191
0
1
0
95
118
0
282
0
123
0
0
0
0
0
0
0
0
0
0
9
0
0
C
Total number of identified contigs or genomes habouring the highly active (FRET >0.7) class A, B & X 2A-peptides (380)
58 (15.3%)
43 (11.3%)
279 (73.4%)
Class A (187)
6 3.2%
20 10.7%
161 86.1%
Class B (71)
71 100.0%
Class X (122)
38 31.1%
47 38.5%
37 30.3%
RNA viruses
Non−viral RTEs
Others

Figure S6

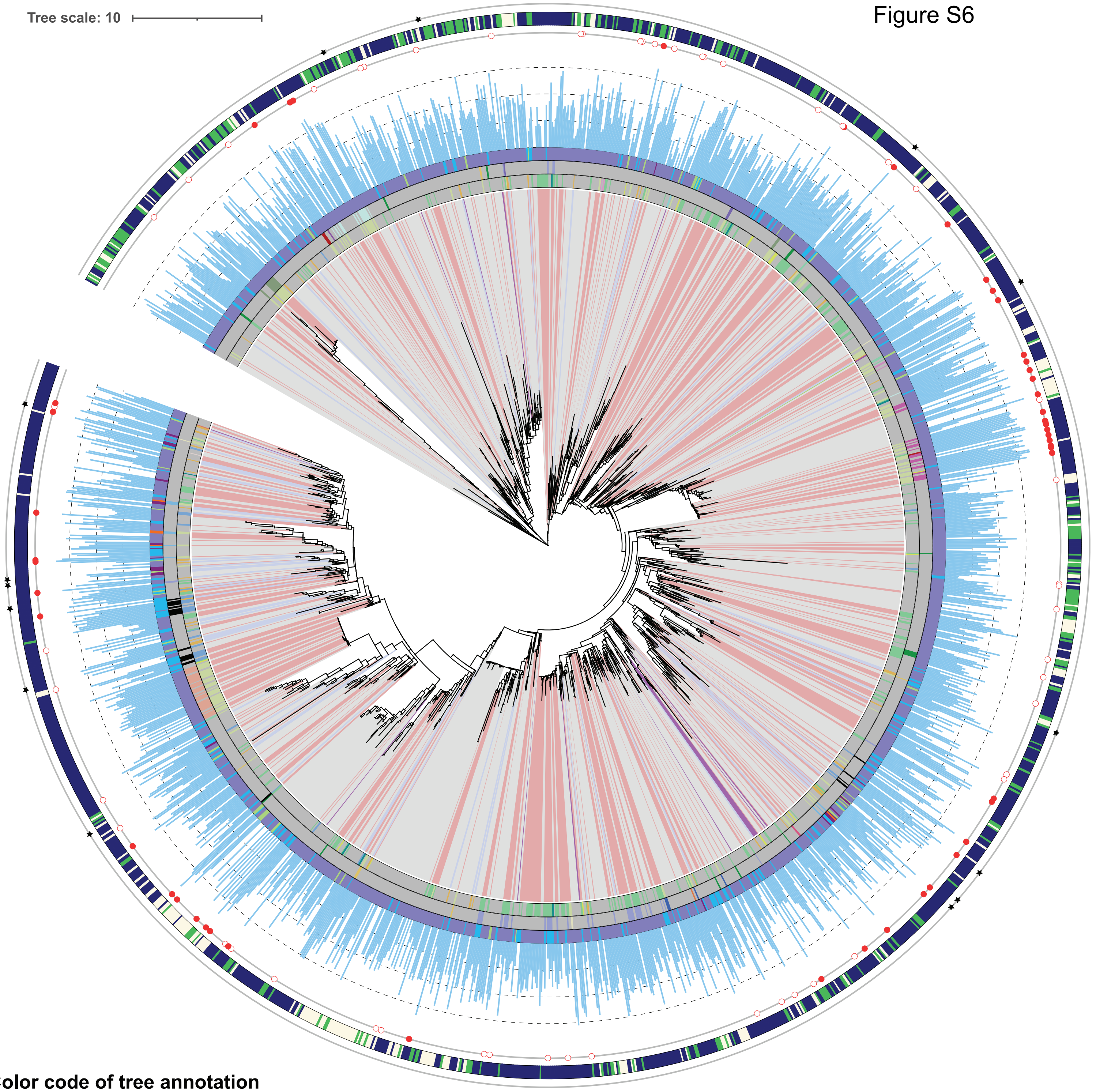


#### Color code of tree annotation

**Tree clades: Virus order**
- o_Ghabrivirales
- o_Hareavirales
- o_Nodamuvirales
- o_Picornavirales
- Unclassified

**1st ring: Virus family**
- f_Caliciviridae
- f_Dicistroviridae
- f_Iflaviridae
- f_Nodaviridae
- f_Phenuiviridae
- f_Picornaviridae
- f_Polycipiviridae
- f_Secoviridae
- f_Sinhaliviridae
- f_Solinviviridae
- f_Totiviridae
- Unclassified

**2nd ring: Virus genus**
- g_Aparavirus
- g_Aphthovirus
- g_Avihepatovirus
- g_Avisivirus
- g_Bopivirus
- g_Cardiovirus
- g_Cosavirus
- g_Cripavirus
- g_Crohivirus
- g_Hunnivirus
- g_Iflavirus
- g_Kobuvirus
- g_Marsupivirus
- g_Mischivirus
- g_Mosavirus
- g_Parechovirus
- g_Pasivirus
- g_Shanbavirus
- g_Teschovirus
- Unclassified

**3rd ring: 2A-peptide number in the genome**
- 1 per genome
- 2 per genome
- 3 per genome
- 4 per genome
- 5 per genome

**4th ring: 2A-peptide activity score (bar plot)**
- Score > 0
- Score = 0

**5th ring: 2A-peptide activity rank**
- High: top 25% activity scores
- Low: bottom 25% activity scores

**6th ring: 2A-peptide attribute**
- RNA viruses in kindom Orthornavirae
- Reverse-transcribing elements (RTEs)
- Unclassified

**7th ring: Back-to-back 2A-peptide**
- Virus with back-to-back 2A-peptide

Figure S7

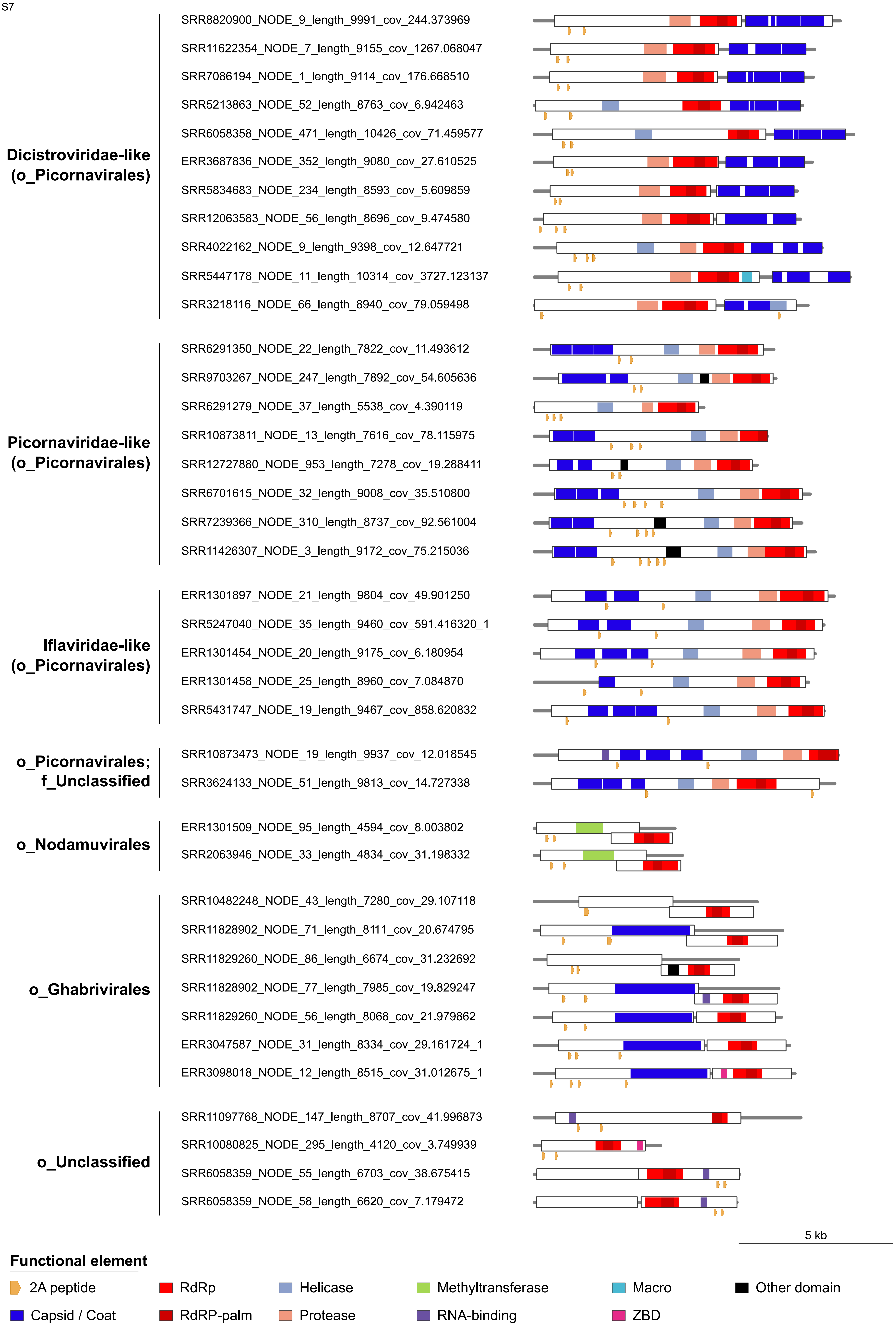
Dicistroviridae-like
(o_Picornavirales)
SRR8820900_NODE_9_length_9991_cov_244.373969
SRR11622354_NODE_7_length_9155_cov_1267.068047
SRR7086194_NODE_1_length_9114_cov_176.668510
SRR5213863_NODE_52_length_8763_cov_6.942463
SRR6058358_NODE_471_length_10426_cov_71.459577
ERR3687836_NODE_352_length_9080_cov_27.610525
SRR5834683_NODE_234_length_8593_cov_5.609859
SRR12063583_NODE_56_length_8696_cov_9.474580
SRR4022162_NODE_9_length_9398_cov_12.647721
SRR5447178_NODE_11_length_10314_cov_3727.123137
SRR3218116_NODE_66_length_8940_cov_79.059498
Picornaviridae-like
(o_Picornavirales)
SRR6291350_NODE_22_length_7822_cov_11.493612
SRR9703267_NODE_247_length_7892_cov_54.605636
SRR6291279_NODE_37_length_5538_cov_4.390119
SRR10873811_NODE_13_length_7616_cov_78.115975
SRR12727880_NODE_953_length_7278_cov_19.288411
SRR6701615_NODE_32_length_9008_cov_35.510800
SRR7239366_NODE_310_length_8737_cov_92.561004
SRR11426307_NODE_3_length_9172_cov_75.215036
Iflaviridae-like
(o_Picornavirales)
ERR1301897_NODE_21_length_9804_cov_49.901250
SRR5247040_NODE_35_length_9460_cov_591.416320_1
ERR1301454_NODE_20_length_9175_cov_6.180954
ERR1301458_NODE_25_length_8960_cov_7.084870
SRR5431747_NODE_19_length_9467_cov_858.620832
o_Picornavirales;
f_Unclassified
SRR10873473_NODE_19_length_9937_cov_12.018545
SRR3624133_NODE_51_length_9813_cov_14.727338
o_Nodamuvirales
ERR1301509_NODE_95_length_4594_cov_8.003802
SRR2063946_NODE_33_length_4834_cov_31.198332
o_Ghabrivirales
SRR10482248_NODE_43_length_7280_cov_29.107118
SRR11828902_NODE_71_length_8111_cov_20.674795
SRR11829260_NODE_86_length_6674_cov_31.232692
SRR11828902_NODE_77_length_7985_cov_19.829247
SRR11829260_NODE_56_length_8068_cov_21.979862
ERR3047587_NODE_31_length_8334_cov_29.161724_1
ERR3098018_NODE_12_length_8515_cov_31.012675_1
o_Unclassified
SRR11097768_NODE_147_length_8707_cov_41.996873
SRR10080825_NODE_295_length_4120_cov_3.749939
SRR6058359_NODE_55_length_6703_cov_38.675415
SRR6058359_NODE_58_length_6620_cov_7.179472
5 kb
Functional element
2A peptide
RdRp
Helicase
Methyltransferase
Macro
Other domain
Capsid / Coat
RdRP-palm
Protease
RNA-binding
ZBD

Figure S8

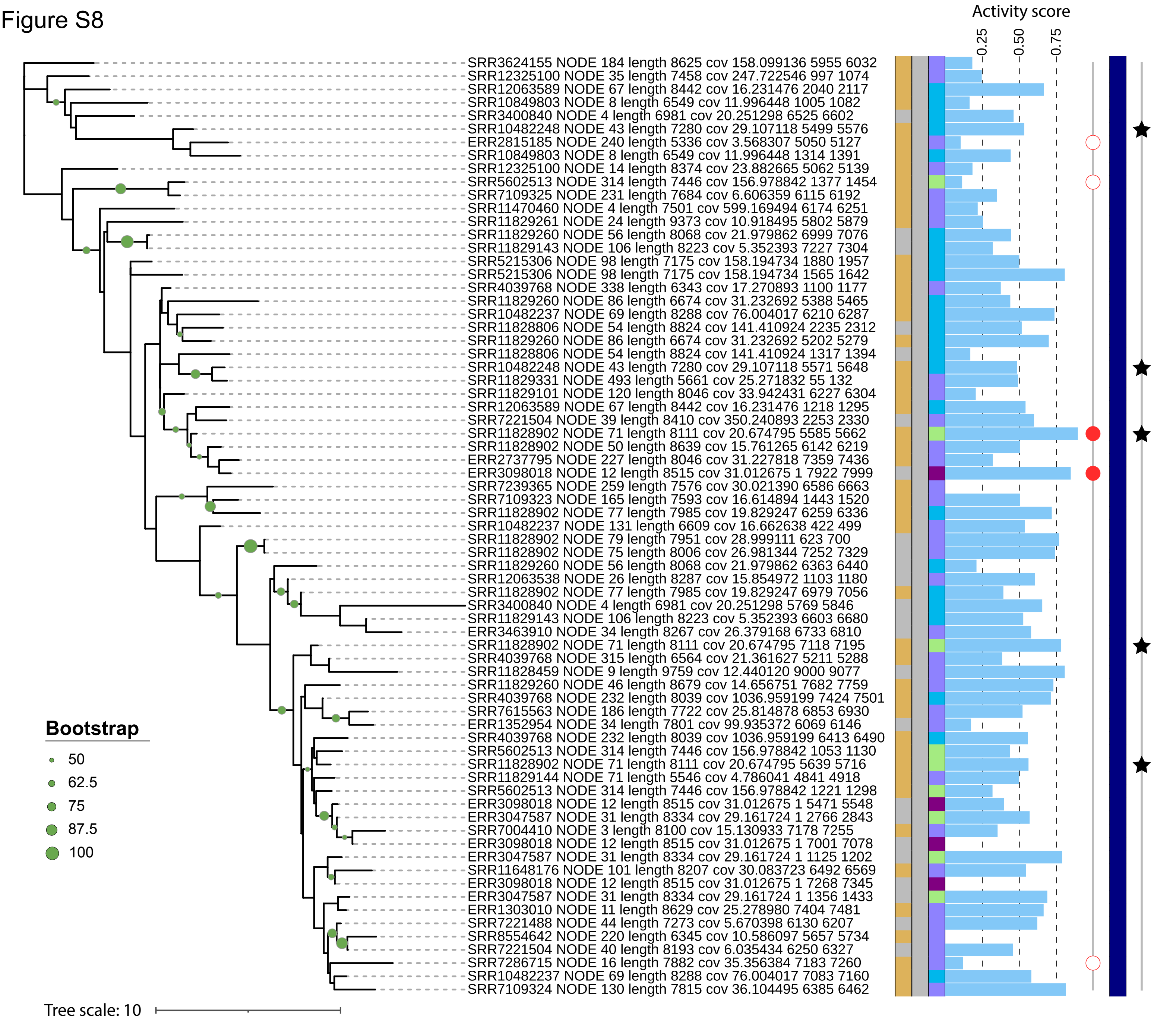


### Color code of annotation

**Tree clades: Virus order**
- o_Ghabrivirales
- o_Hareavirales
- o_Nodamuvirales
- o_Picornavirales
- Unclassified

**1st ring: Virus family**
- f_Caliciviridae
- f_Dicistroviridae
- f_Iflaviridae
- f_Nodaviridae
- f_Phenuiviridae
- f_Picornaviridae
- f_Polycipiviridae
- f_Secoviridae
- f_Sinhaliviridae
- f_Solinviviridae
- f_Totiviridae
- Unclassified

**2nd ring: Virus genus**
- g_Aparavirus
- g_Aphthovirus
- g_Avihepatovirus
- g_Avisivirus
- g_Bopivirus
- g_Cardiovirus
- g_Cosavirus
- g_Cripavirus
- g_Crohivirus
- g_Hunnivirus
- g_Iflavirus
- g_Kobuvirus
- g_Marsupivirus
- g_Mischivirus
- g_Mosavirus
- g_Parechovirus
- g_Pasivirus
- g_Shanbavirus
- g_Teschovirus
- Unclassified

**3rd ring: 2A-peptide number in the genome**
- 1 per genome
- 2 per genome
- 3 per genome
- 4 per genome
- 5 per genome

**4th ring: 2A-peptide activity score (bar plot)**
- Score > 0
- Score = 0

**5th ring: 2A-peptide activity rank**
- High: top 25% activity scores
- Low: bottom 25% activity scores

**6th ring: 2A-peptide attribute**
- RNA viruses in kindom Orthornavirae
- Reverse-transcribing elements (RTEs)
- Unclassified

**7th ring: Back-to-back 2A-peptide**
- ★ Virus with back-to-back 2A-peptide

Figure S9

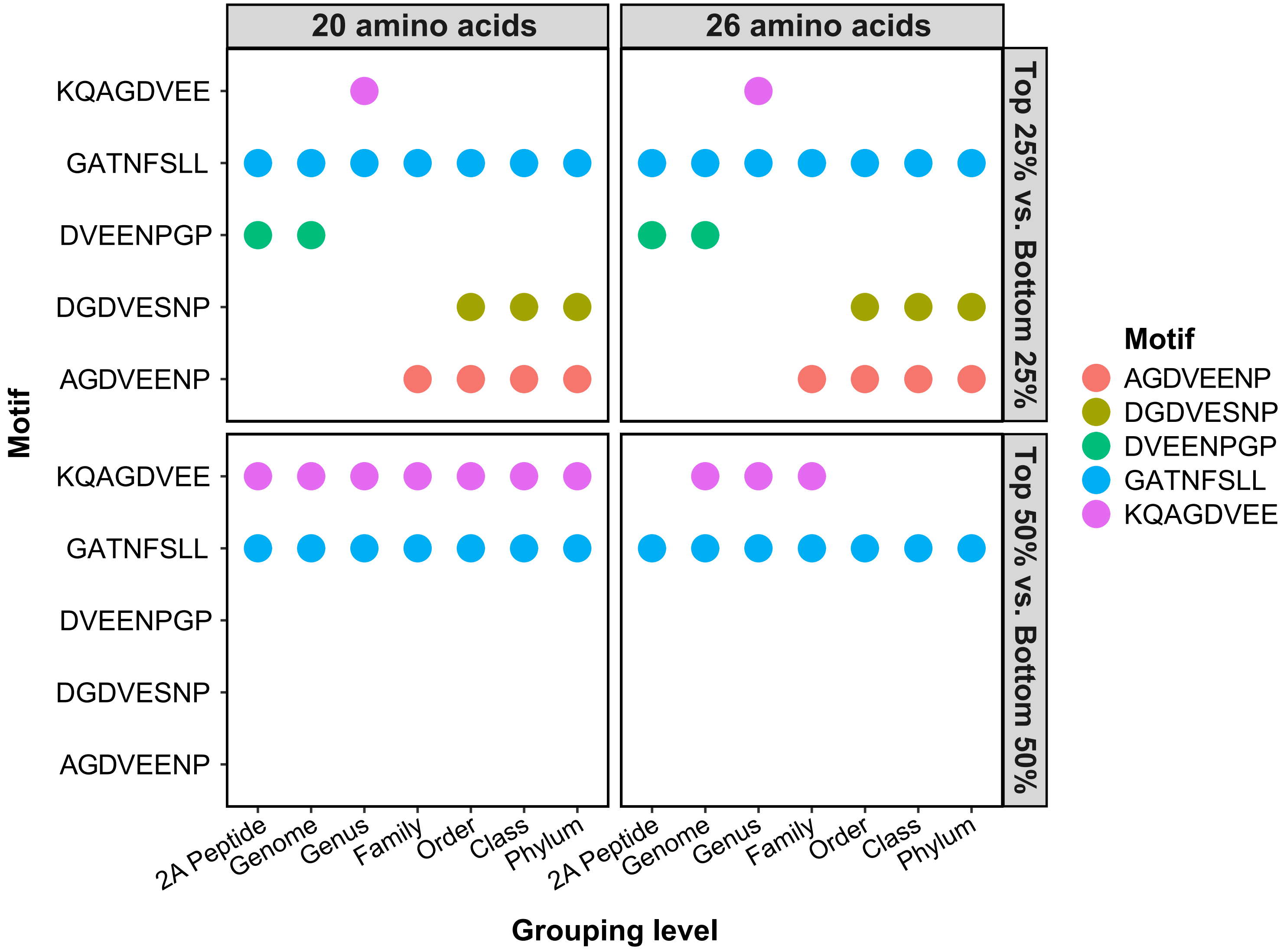
20 amino acids
26 amino acids
Top 25% vs. Bottom 25%
Top 50% vs. Bottom 50%
Motif
KQAGDVEE
GATNFSLL
DVEENPGP
DGDVESNP
AGDVEENP
2A Peptide
Genome
Genus
Family
Order
Class
Phylum
Grouping level
Motif
AGDVEENP
DGDVESNP
DVEENPGP
GATNFSLL
KQAGDVEE

Figure S10

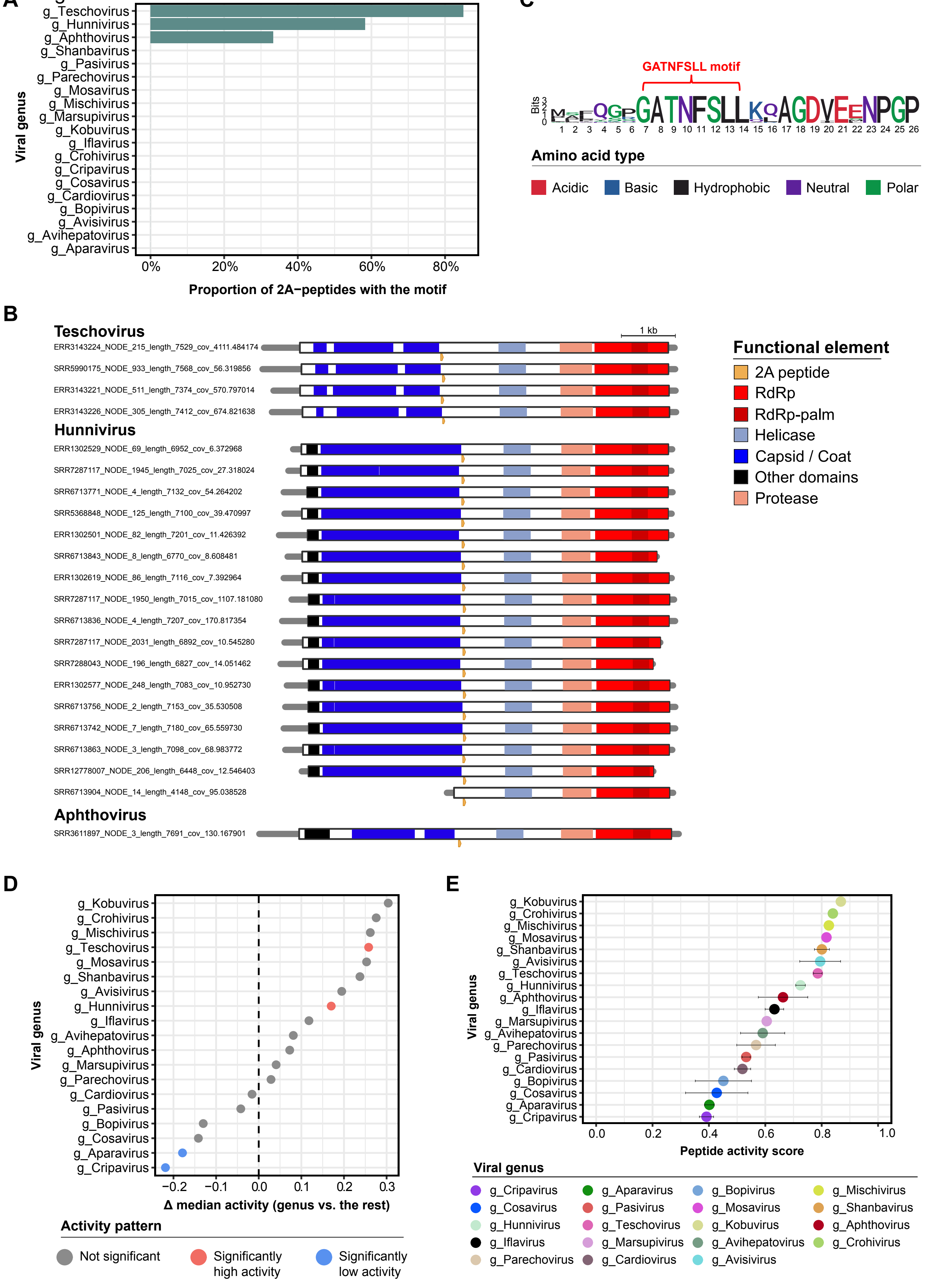

A
g_Teschovirus
g_Hunnivirus
g_Aphthovirus
g_Shanbavirus
g_Pasivirus
g_Parechovirus
g_Mosavirus
g_Mischivirus
g_Marsupivirus
g_Kobuvirus
g_Iflavirus
g_Crohivirus
g_Cripavirus
g_Cosavirus
g_Cardiovirus
g_Bopivirus
g_Avisivirus
g_Avihepatovirus
g_Aparavirus
Viral genus
0%
20%
40%
60%
80%
Proportion of 2A−peptides with the motif
C
GATNFSLL motif
Bits
Amino acid type
Acidic
Basic
Hydrophobic
Neutral
Polar
B
1 kb
Teschovirus
ERR3143224_NODE_215_length_7529_cov_4111.484174
SRR5990175_NODE_933_length_7568_cov_56.319856
ERR3143221_NODE_511_length_7374_cov_570.797014
ERR3143226_NODE_305_length_7412_cov_674.821638
Hunnivirus
ERR1302529_NODE_69_length_6952_cov_6.372968
SRR7287117_NODE_1945_length_7025_cov_27.318024
SRR6713771_NODE_4_length_7132_cov_54.264202
SRR5368848_NODE_125_length_7100_cov_39.470997
ERR1302501_NODE_82_length_7201_cov_11.426392
SRR6713843_NODE_8_length_6770_cov_8.608481
ERR1302619_NODE_86_length_7116_cov_7.392964
SRR7287117_NODE_1950_length_7015_cov_1107.181080
SRR6713836_NODE_4_length_7207_cov_170.817354
SRR7287117_NODE_2031_length_6892_cov_10.545280
SRR7288043_NODE_196_length_6827_cov_14.051462
ERR1302577_NODE_248_length_7083_cov_10.952730
SRR6713756_NODE_2_length_7153_cov_35.530508
SRR6713742_NODE_7_length_7180_cov_65.559730
SRR6713863_NODE_3_length_7098_cov_68.983772
SRR12778007_NODE_206_length_6448_cov_12.546403
SRR6713904_NODE_14_length_4148_cov_95.038528
Aphthovirus
SRR3611897_NODE_3_length_7691_cov_130.167901
Functional element
2A peptide
RdRp
RdRp-palm
Helicase
Capsid / Coat
Other domains
Protease
D
g_Kobuvirus
g_Crohivirus
g_Mischivirus
g_Teschovirus
g_Mosavirus
g_Shanbavirus
g_Avisivirus
g_Hunnivirus
g_Iflavirus
g_Avihepatovirus
g_Aphthovirus
g_Marsupivirus
g_Parechovirus
g_Cardiovirus
g_Pasivirus
g_Bopivirus
g_Cosavirus
g_Aparavirus
g_Cripavirus
Viral genus
−0.2
−0.1
0.0
0.1
0.2
0.3
Δ median activity (genus vs. the rest)
Activity pattern
Not significant
Significantly high activity
Significantly low activity
E
g_Kobuvirus
g_Crohivirus
g_Mischivirus
g_Mosavirus
g_Shanbavirus
g_Avisivirus
g_Teschovirus
g_Hunnivirus
g_Aphthovirus
g_Iflavirus
g_Marsupivirus
g_Avihepatovirus
g_Parechovirus
g_Pasivirus
g_Cardiovirus
g_Bopivirus
g_Cosavirus
g_Aparavirus
g_Cripavirus
Viral genus
0.0
0.2
0.4
0.6
0.8
1.0
Peptide activity score
Viral genus
g_Cripavirus
g_Aparavirus
g_Bopivirus
g_Mischivirus
g_Cosavirus
g_Pasivirus
g_Mosavirus
g_Shanbavirus
g_Hunnivirus
g_Teschovirus
g_Kobuvirus
g_Aphthovirus
g_Iflavirus
g_Marsupivirus
g_Avihepatovirus
g_Crohivirus
g_Parechovirus
g_Cardiovirus
g_Avisivirus

Figure S11

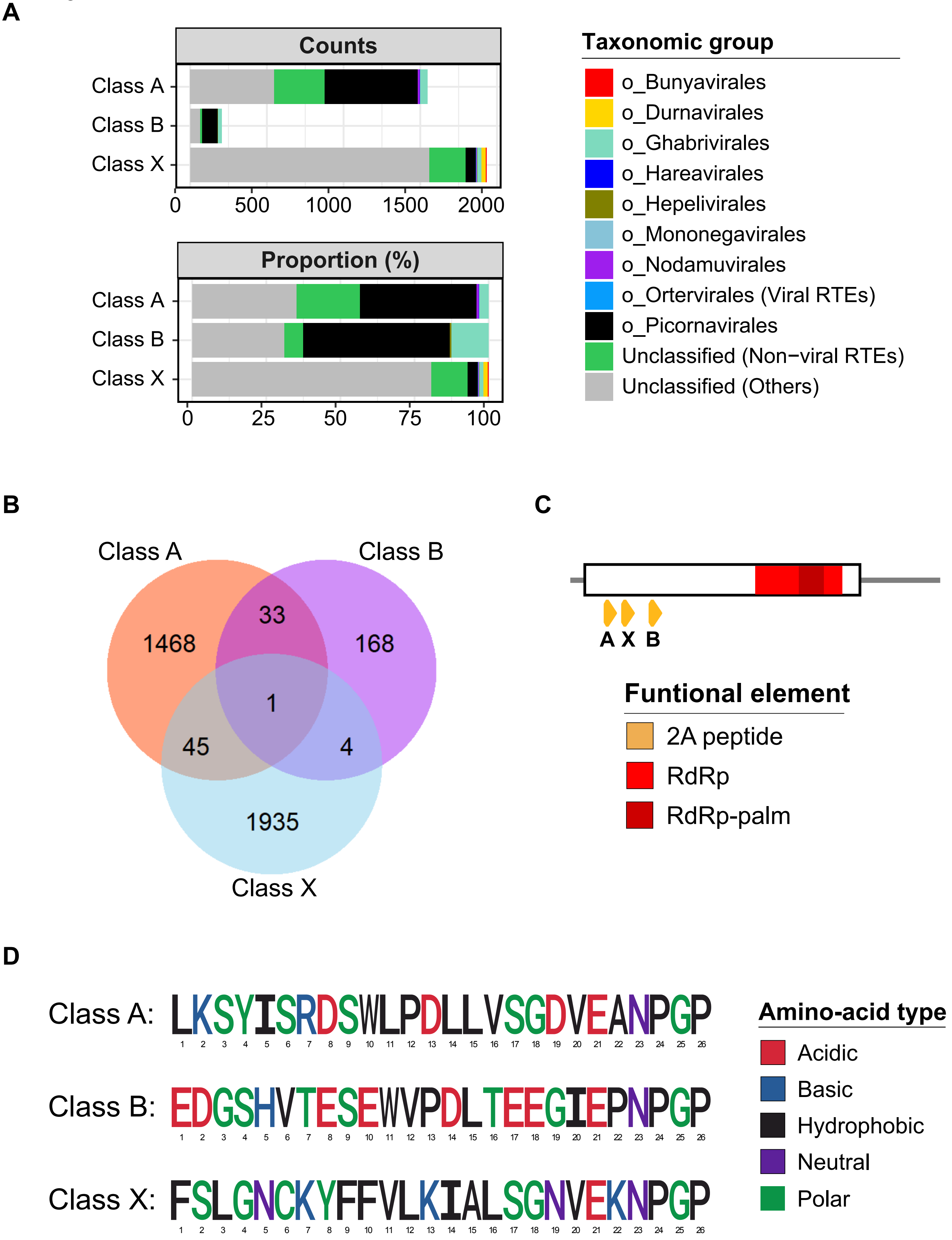
A
Counts
Class A
Class B
Class X
0
500
1000
1500
2000
Proportion (%)
Class A
Class B
Class X
0
25
50
75
100
Taxonomic group
o_Bunyavirales
o_Durnavirales
o_Ghabrivirales
o_Hareavirales
o_Hepelivirales
o_Mononegavirales
o_Nodamuvirales
o_Ortervirales (Viral RTEs)
o_Picornavirales
Unclassified (Non−viral RTEs)
Unclassified (Others)
B
Class A
Class B
1468
33
168
1
45
4
1935
Class X
C
A X B
Funtional element
2A peptide
RdRp
RdRp-palm
D
Class A: LKSYISRDSWLPDLLVSGDVEANPGP
Class B: EDGSHVTESEWVPDLTEEGIEPNPGP
Class X: FSLGNCKYFFVLKIALSGNVEKNPGP
1 2 3 4 5 6 7 8 9 10 11 12 13 14 15 16 17 18 19 20 21 22 23 24 25 26
Amino-acid type
Acidic
Basic
Hydrophobic
Neutral
Polar

Figure S12

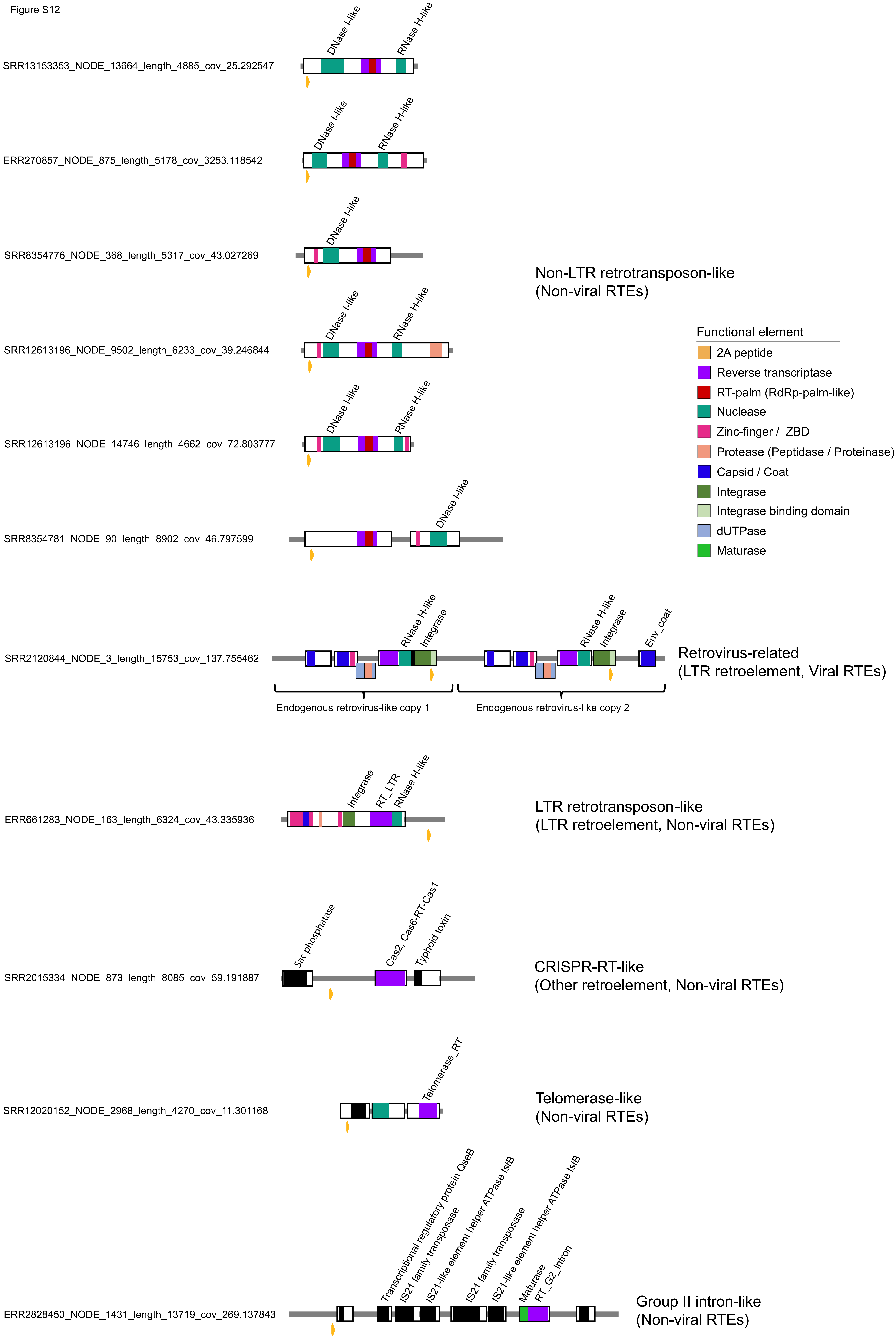

SRR13153353_NODE_13664_length_4885_cov_25.292547
DNase I-like
RNase H-like
ERR270857_NODE_875_length_5178_cov_3253.118542
DNase I-like
RNase H-like
SRR8354776_NODE_368_length_5317_cov_43.027269
DNase I-like
SRR12613196_NODE_9502_length_6233_cov_39.246844
DNase I-like
RNase H-like
SRR12613196_NODE_14746_length_4662_cov_72.803777
DNase I-like
RNase H-like
SRR8354781_NODE_90_length_8902_cov_46.797599
DNase I-like
Non-LTR retrotransposon-like
(Non-viral RTEs)
Functional element
2A peptide
Reverse transcriptase
RT-palm (RdRp-palm-like)
Nuclease
Zinc-finger / ZBD
Protease (Peptidase / Proteinase)
Capsid / Coat
Integrase
Integrase binding domain
dUTPase
Maturase
SRR2120844_NODE_3_length_15753_cov_137.755462
RNase H-like
Integrase
RNase H-like
Integrase
Env_coat
Endogenous retrovirus-like copy 1
Endogenous retrovirus-like copy 2
Retrovirus-related
(LTR retroelement, Viral RTEs)
ERR661283_NODE_163_length_6324_cov_43.335936
Integrase
RT_LTR
RNase H-like
LTR retrotransposon-like
(LTR retroelement, Non-viral RTEs)
SRR2015334_NODE_873_length_8085_cov_59.191887
Sac phosphatase
Cas2, Cas6-RT-Cas1
Typhoid toxin
CRISPR-RT-like
(Other retroelement, Non-viral RTEs)
SRR12020152_NODE_2968_length_4270_cov_11.301168
Telomerase_RT
Telomerase-like
(Non-viral RTEs)
ERR2828450_NODE_1431_length_13719_cov_269.137843
Transcriptional regulatory protein QseB
IS21 family transposase
IS21-like element helper ATPase IstB
IS21 family transposase
IS21-like element helper ATPase IstB
Maturase
RT_G2_intron
Group II intron-like
(Non-viral RTEs)

Figure S13

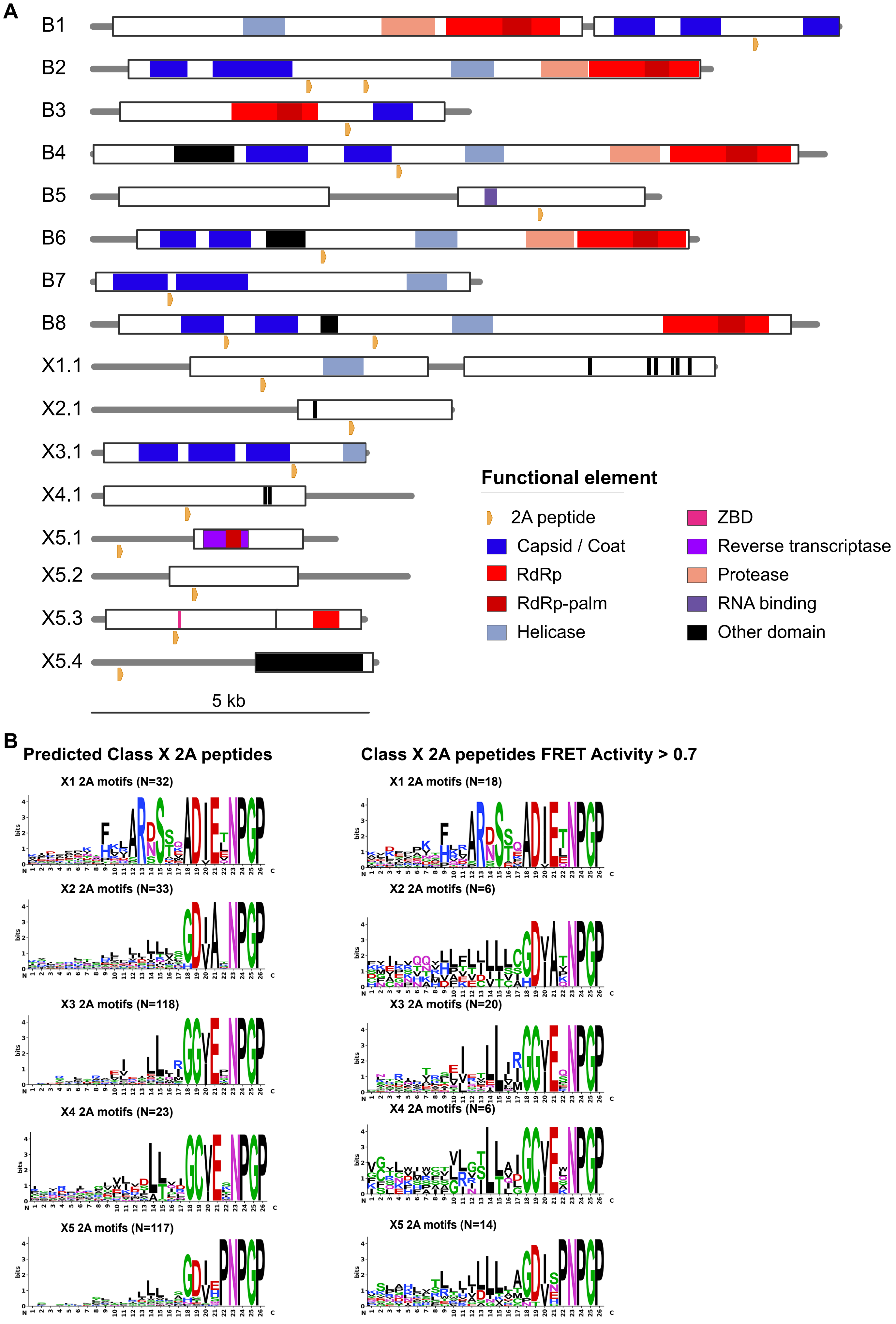
A
B1
B2
B3
B4
B5
B6
B7
B8
X1.1
X2.1
X3.1
X4.1
X5.1
X5.2
X5.3
X5.4
5 kb
Functional element
2A peptide
Capsid / Coat
RdRp
RdRp-palm
Helicase
ZBD
Reverse transcriptase
Protease
RNA binding
Other domain
B
Predicted Class X 2A peptides
Class X 2A pepetides FRET Activity > 0.7
X1 2A motifs (N=32)
X1 2A motifs (N=18)
X2 2A motifs (N=33)
X2 2A motifs (N=6)
X3 2A motifs (N=118)
X3 2A motifs (N=20)
X4 2A motifs (N=23)
X4 2A motifs (N=6)
X5 2A motifs (N=117)
X5 2A motifs (N=14)

Figure S14

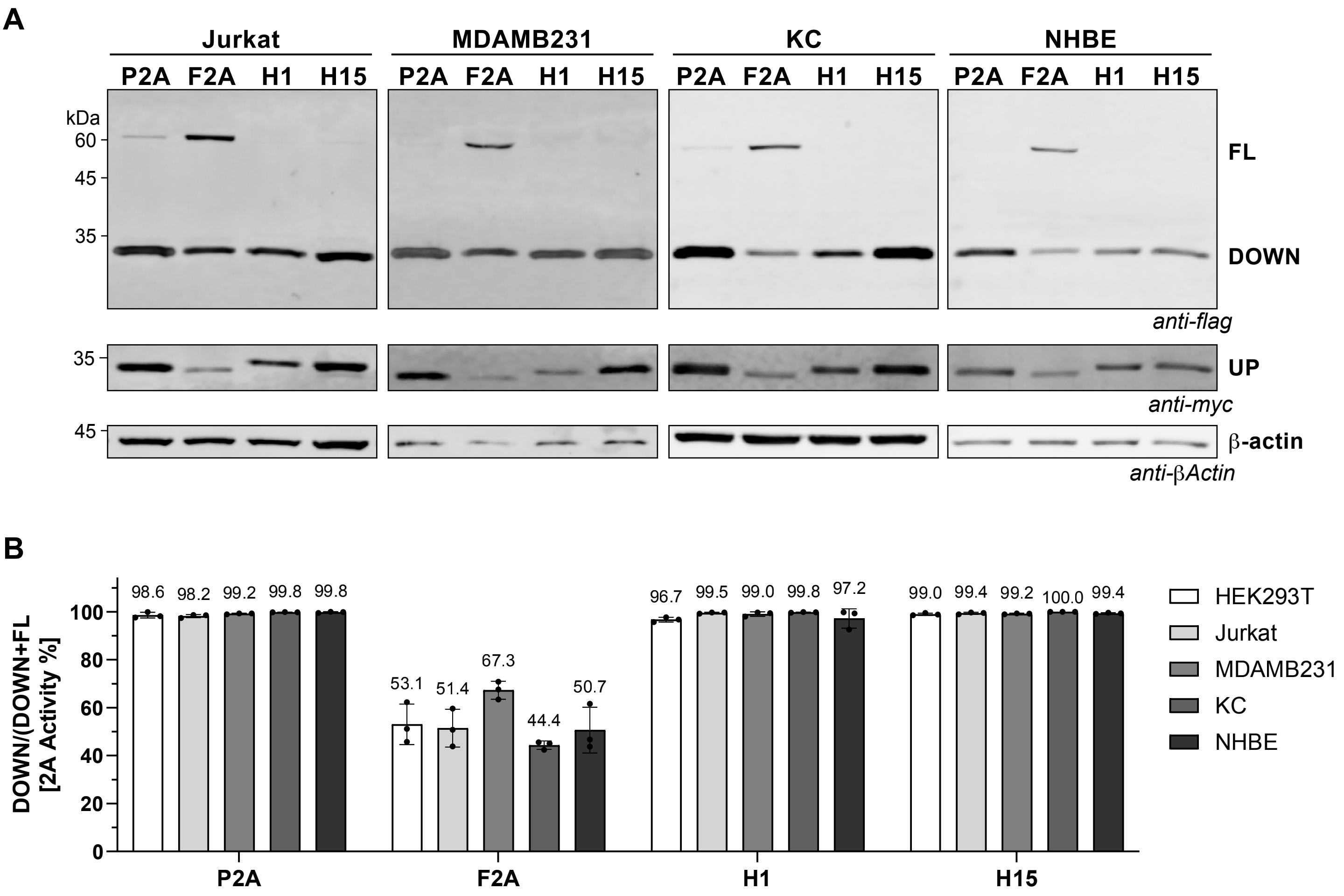


**Supplementary Figure 14. High activity 2A peptides maintain robust StopGo activity across diverse mammalian cell lines**. (A) Representative Immunoblots of the P2A, F2A, H1, and H15 reporters expressed in Jurkat, MDA-MB-231, KC, and NHBE cells. The full-length fused protein (FL; myc-mClover-2A-mRuby-Flag) and downstream product (DOWN; mRuby-Flag) were detected with anti-Flag, the upstream product (UP; myc-mClover) with anti-myc, and β-actin with anti-β-actin as a loading control. (B) Quantification of 2A activity for each 2A peptide across five cell lines (HEK293T, Jurkat, MDA-MB-231, KC, and NHBE), calculated as relative 2A activity % = Down/(Down + FL). Error bars represent Mean ± SD; values above each bar indicate mean percent activity.

Figure S15

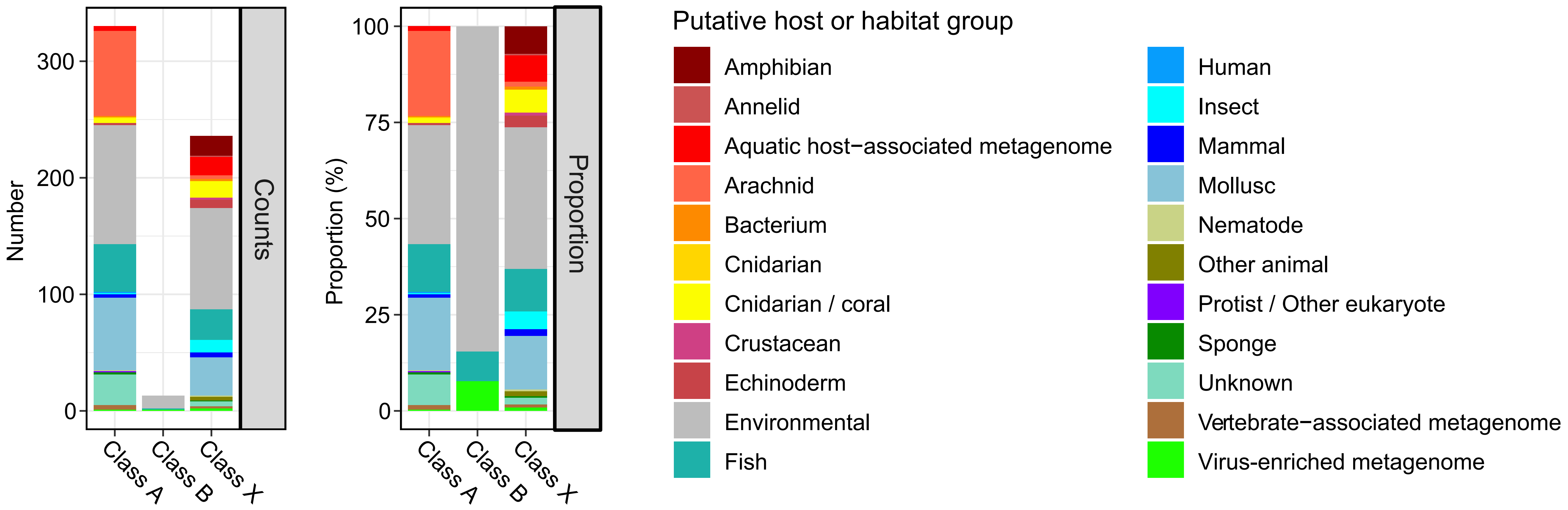


Supplementary Figure 15. Putative hosts or habitats of the identified 2A-peptide-containing RTEs. Putative host or habitat assignments were inferred from the sample metadata associated with the original sequences, retrieved from the NCBI SRA, EMBL ENA, and DDBJ DRA databases. The left panel shows the number of 2A-peptide-containing RTE contigs identified across different host or habitat groups, while the right panel shows their proportional distribution among all identified 2A-peptide-containing RTEs.
